# Structure-Activity Relationship and Target Investigation of Thiophen-2-yl-Pyrimidines against *Schistosoma* species

**DOI:** 10.1101/2025.10.07.681061

**Authors:** Karol R. Francisco, Bobby Lucero, Darius Yohannan, Carmine Varricchio, Anny Lam, Jessica Sebastiani, Lawrence J. Liu, Yujie Uli Sun, Jorge Jacinto, Michele Renzulli, Ludovica Monti, Alexis Lona, Irina Kufareva, Andrea Brancale, Carlo Ballatore, Thibault Alle, Conor R. Caffrey

## Abstract

Chemotherapeutic options for schistosomiasis, a prevalent infectious disease of poverty, are limited to just one drug, praziquantel (PZQ), and alternatives are needed. Our previous studies identified thiophen-2-yl pyrimidines (TPPs), which are structurally derived from microtubule (MT)-active phenylpyrimidines, as potent paralytics of *Schistosoma mansoni*. Although relatively non-toxic to mammalian cells, the progenitor compound, **3**, had poor aqueous solubility and was lipophilic potentially hindering preclinical advancement. To address these issues and expand on the structure-activity and structure-property relationships, 43 new TPP analogs were designed and synthesized, their lipophilicity calculated (cLogP), and their anti-schistosomal activity evaluated in culture. This effort yielded compound **38**, which possessed an oxetane-containing amine moiety at C5, and an *ortho, ortho-*difluoroaniline at C6 of the TPP scaffold. Compared to **3**, compound **38** had better aqueous solubility (46 *vs.* < 0.5 µM) and decreased lipophilicity (logP calc. 4.48 *vs.* 6.81), with toxicity CC_50_ values > 20 µM against three mammalian cell lines. Further, paralytic potency, as measured by the EC_50_ value for adult *S. mansoni* motility, was increased 14.5-fold (538 *vs.* 37 nM), and plasma half-life (t_1/2_) was improved 3-fold, from 0.48 to 1.51 h for a 40% loss in maximum plasma concentration (C_max_). In washout experiments, **38** produced a sustained paralysis of both juvenile and adult *S. mansoni,* possibly suggesting a broader *in vitro* efficacy spectrum compared to PZQ, which is inactive against the juvenile parasite. Also, the two other medically important species, *Schistosoma haematobium* and *Schistosoma japonicum,* were susceptible to **38**. Finally, to identify potential protein targets, we synthesized a TPP photoaffinity labeling (PAL) probe that labeled several *S. mansoni* proteins by SDS-PAGE fluorescence analysis, although, notably, not tubulin, suggesting that the antischistosomal activity of **38** is a function of engaging other targets. Future work with the TPP series will aim to decrease toxicity further while improving PK properties to better support *in vivo* efficacy testing.

## 1. Introduction

Schistosomiasis is caused by trematode flatworms of the genus *Schistosoma*. Transmitted by freshwater snails, an estimated 250 million people are infected with >700 million at risk of infection,^1–4^ most of whom live in sub-Saharan Africa. As a chronic, painful and debilitating disease,^3–5^ schistosomiasis directly contributes to the poverty trap by limiting societal and economic productivity.^6^ ^7^ There is no vaccine, and since the early 1980s, treatment has relied on just one drug, praziquantel (PZQ).^8, 9^ Although safe and reasonably effective, PZQ has both pharmacological and pharmaceutical drawbacks that encourage the search for new drugs.^10–12^

Previously, we identified a series of thiophen-2-yl pyrimidines (TPPs), which produce a potent and sustained paralysis in *ex vivo* adult *Schistosoma mansoni* that is characterized by minor uncoordinated body movements and the inability of the worm’s oral or ventral suckers to operate as holdfasts.^13^ These TPPs share structural similarities with triazolopyrimidines (*e.g*., **1**, Figure 1) and phenylpyrimidines (*e.g*., **2**, Figure 1), investigational anti-tauopathy agents that bind to microtubules (MTs) and modulate MT dynamics.^13–15^ Notwithstanding their structural similarities with these pyrimidines, the anti-schistosomal TPPs do not alter mammalian MT dynamics and exhibit low cytotoxicity to HEK293 cells,^13^ which are typically sensitive to the antimitotic effects of MT-targeting agents.^16^ Furthermore, the paralysis caused by the antischistosomal TPPs, as typified by compound **3** (Figure 1), is not observed after incubation of *S. mansoni* with the MT-active triazolopyrimidines or phenylpyrimidines.^13^

**Figure 1.**
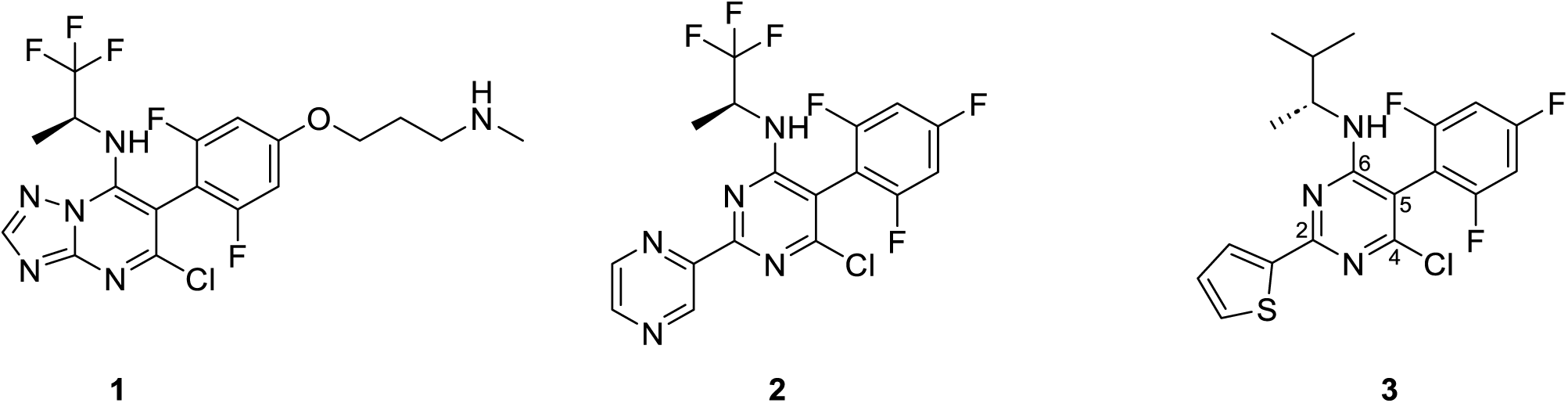
Examples of a triazolopyrimidine, **1**, a phenylpyrimidine, **2**, and a thiophen-2-yl pyrimidine, **3**. Compounds **1** and **2** bind MTs and do not cause paralysis in *S. mansoni*, whereas **3** does not engage MTs but causes paralysis An initial evaluation of the structure-activity relationship (SAR) of TPPs revealed that the presence of a thiophene at C2, a chlorine at C4, a fluorinated phenyl ring at C5, and an amine substituent at C6 were important for antischistosomal activity, with the thiophene as a driver of the paralysis phenotype.^13^ TPP **3** (Figure 1) was the most active congener from these studies; however, its marked lipophilicity,^17^ (a clogP of 6.8) and low aqueous solubility (<0.5 µM at pH 7.4), indicate a poor drug-likeness profile,^18–21^ raising concerns regarding formulation, bioavailability and *in vivo* toxicity.^22^

Accordingly, to further investigate the SAR and the structure-property relationships (SPR), we synthesized a series of 43 TPP derivatives (**4**–**46**, Table 1), calculated their lipophilicities (cLogP), and evaluated their activity against *ex vivo S. mansoni*. We identified novel TPP derivatives that demonstrate decreased lipophilicity, improved aqueous solubility and enhanced potency compared to the initial lead, **3**. In addition, exploration of the mechanism of action through photoaffinity labelling (PAL) experiments revealed that the anti-schistosomal TPPs, unlike their progenitor MT-binding triazolopyrimidines, do not engage tubulin, suggesting that other targets are involved in the paralysis produced by these compounds.

**Table 1.**
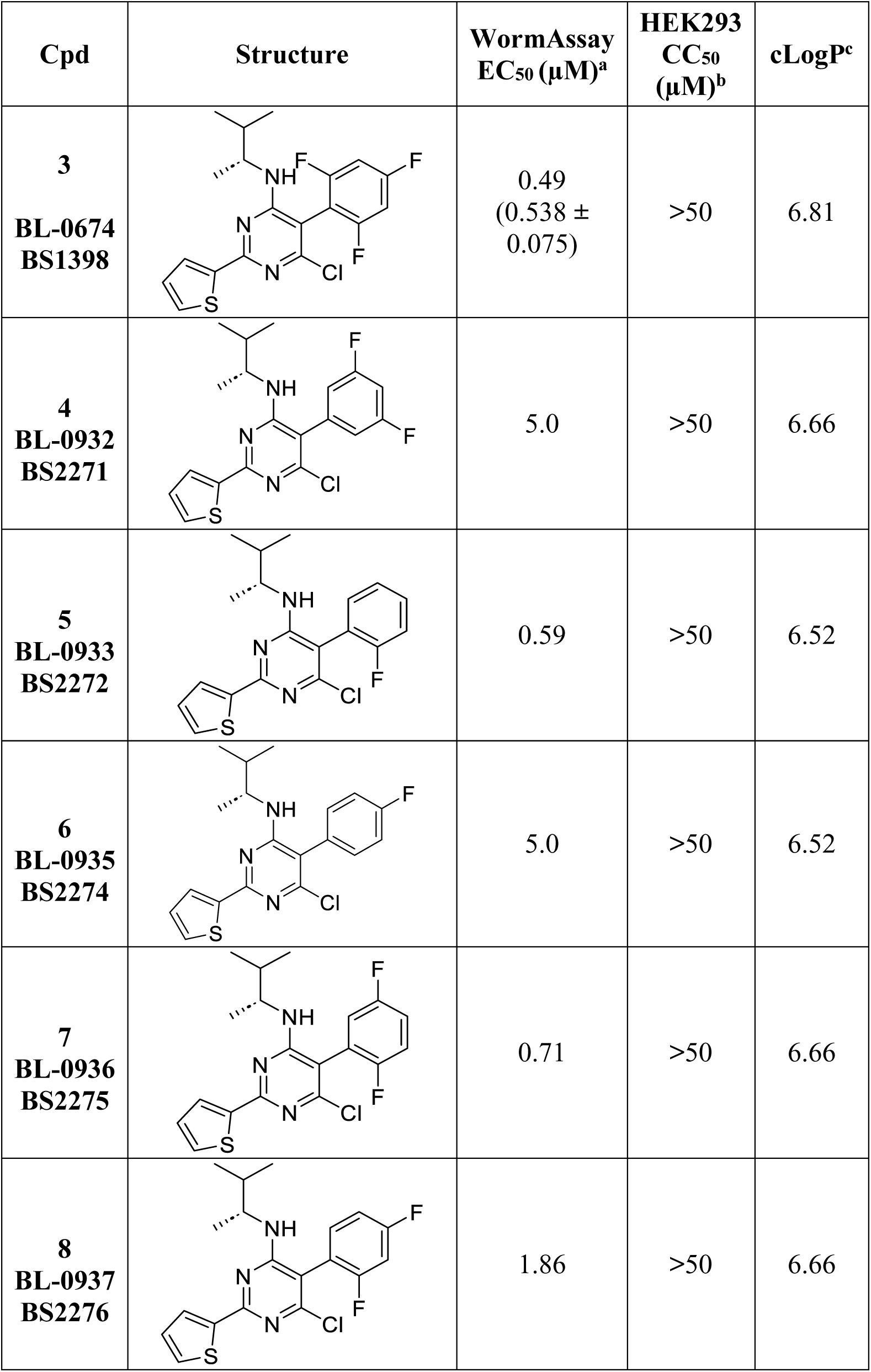
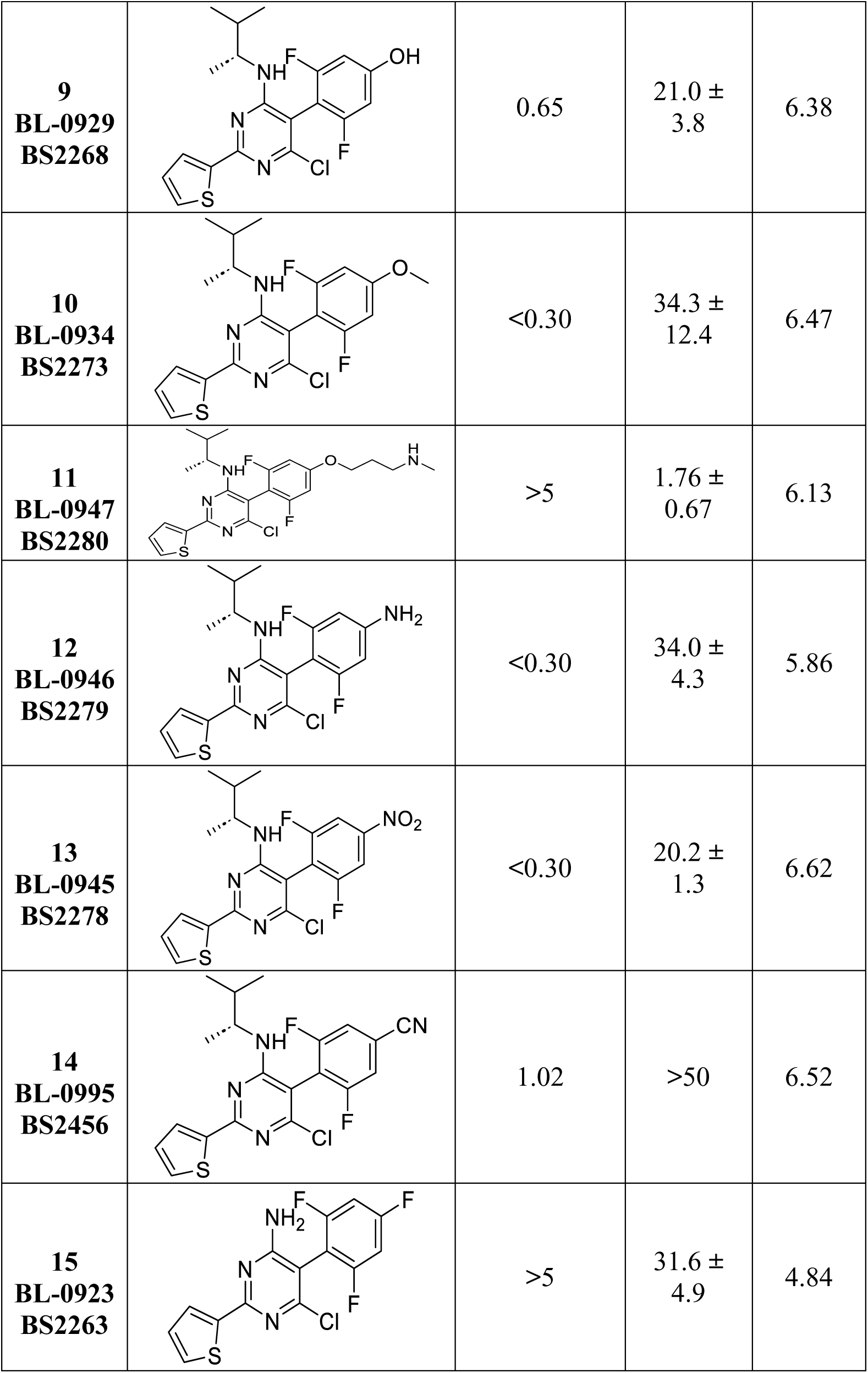
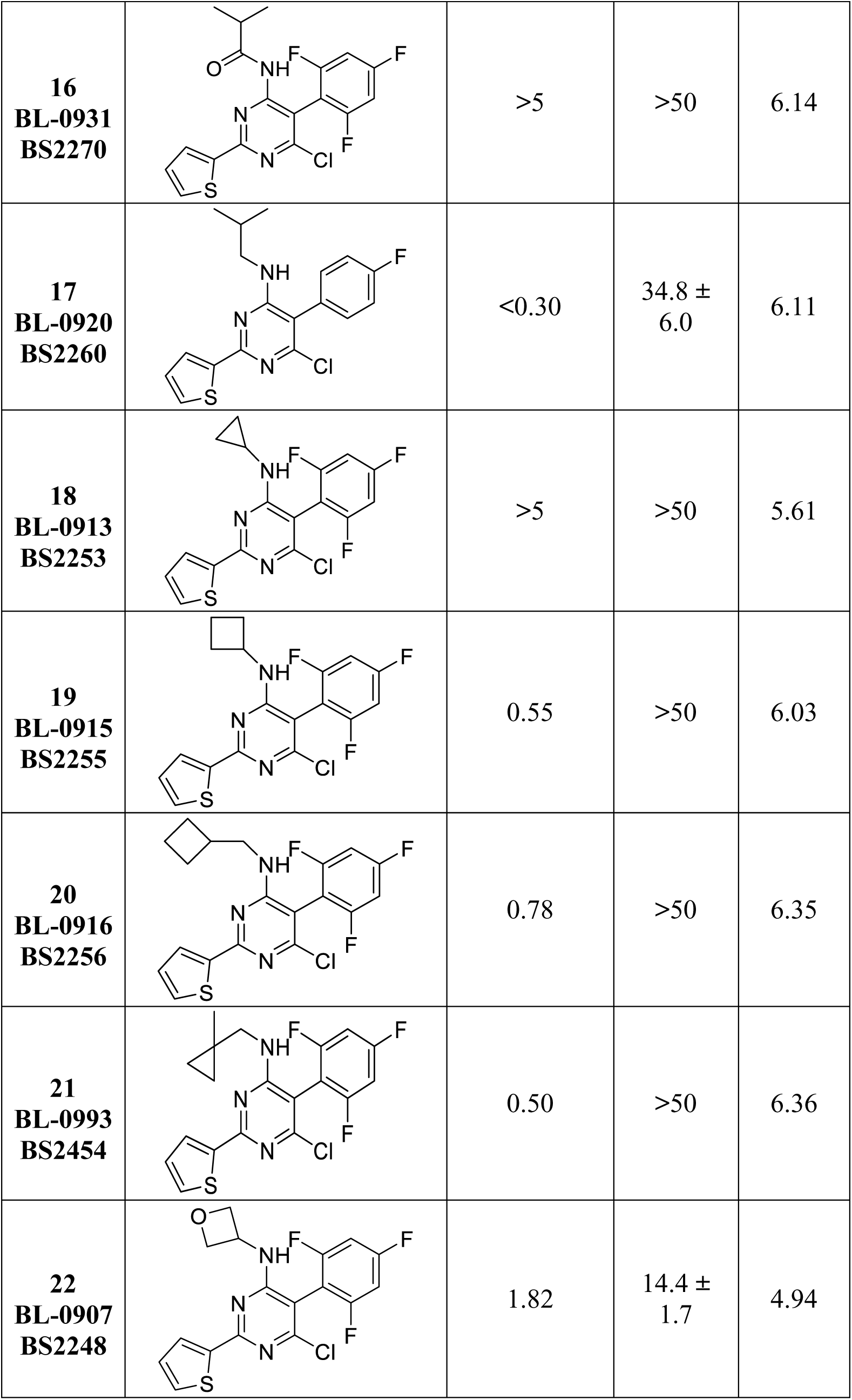
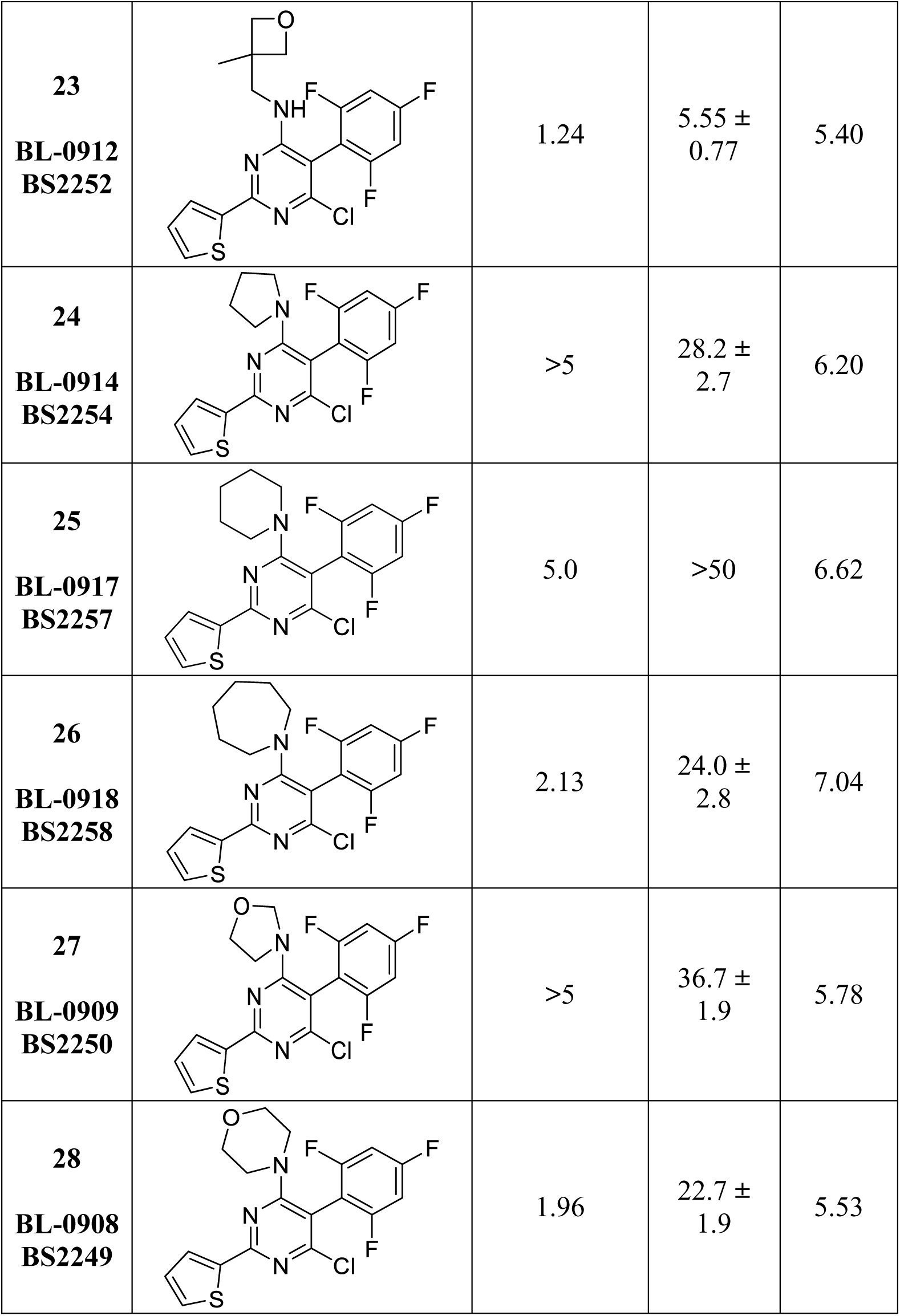
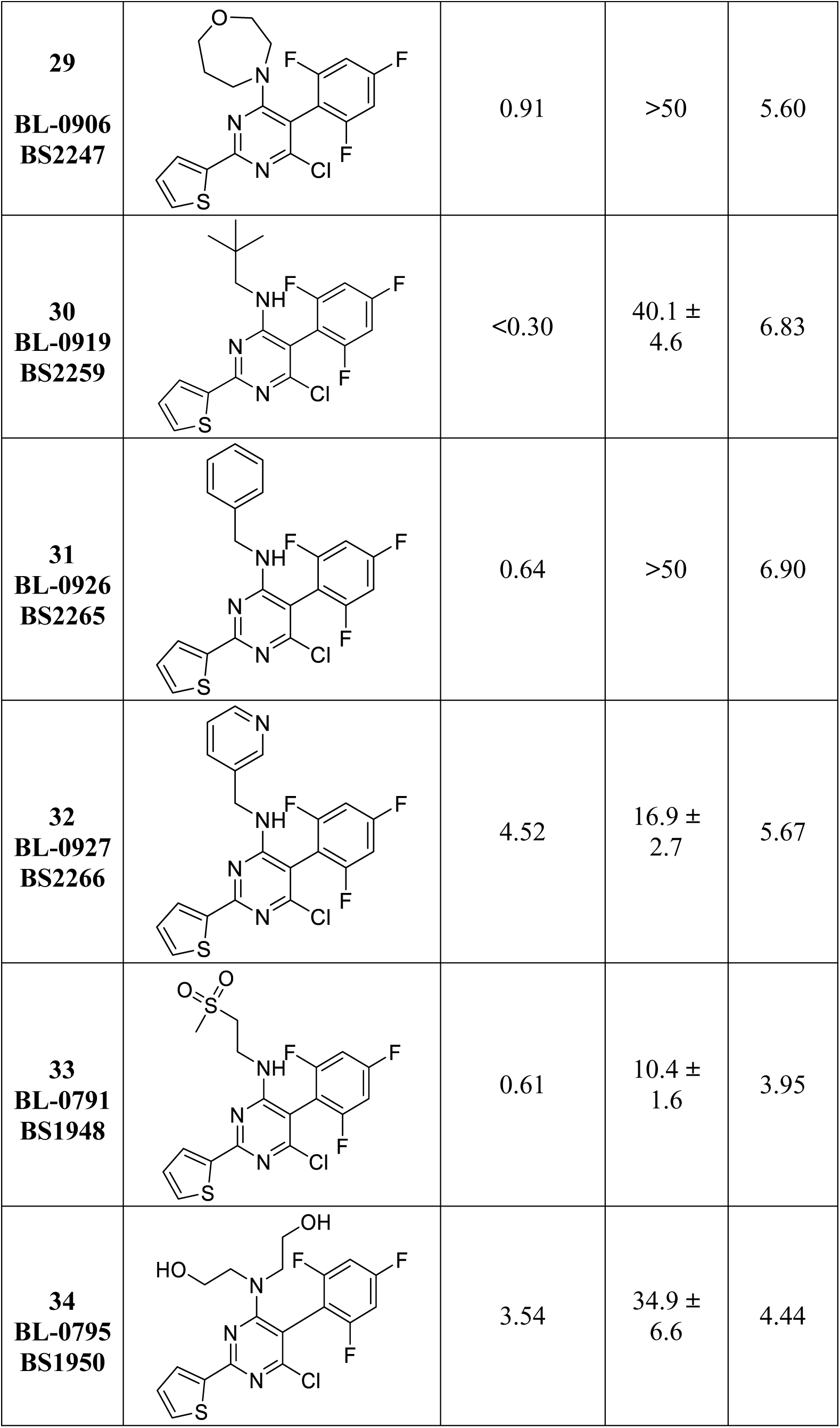
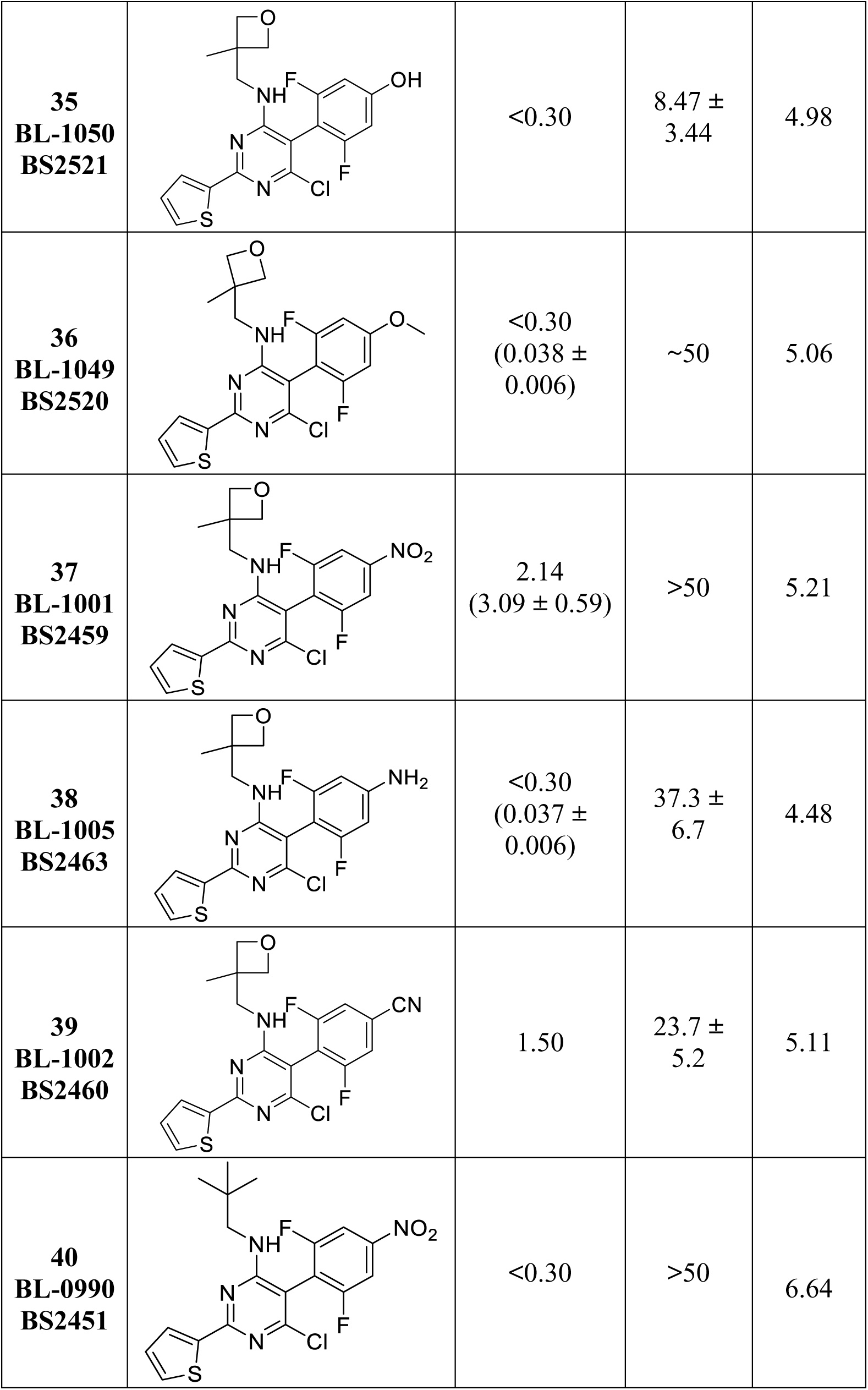
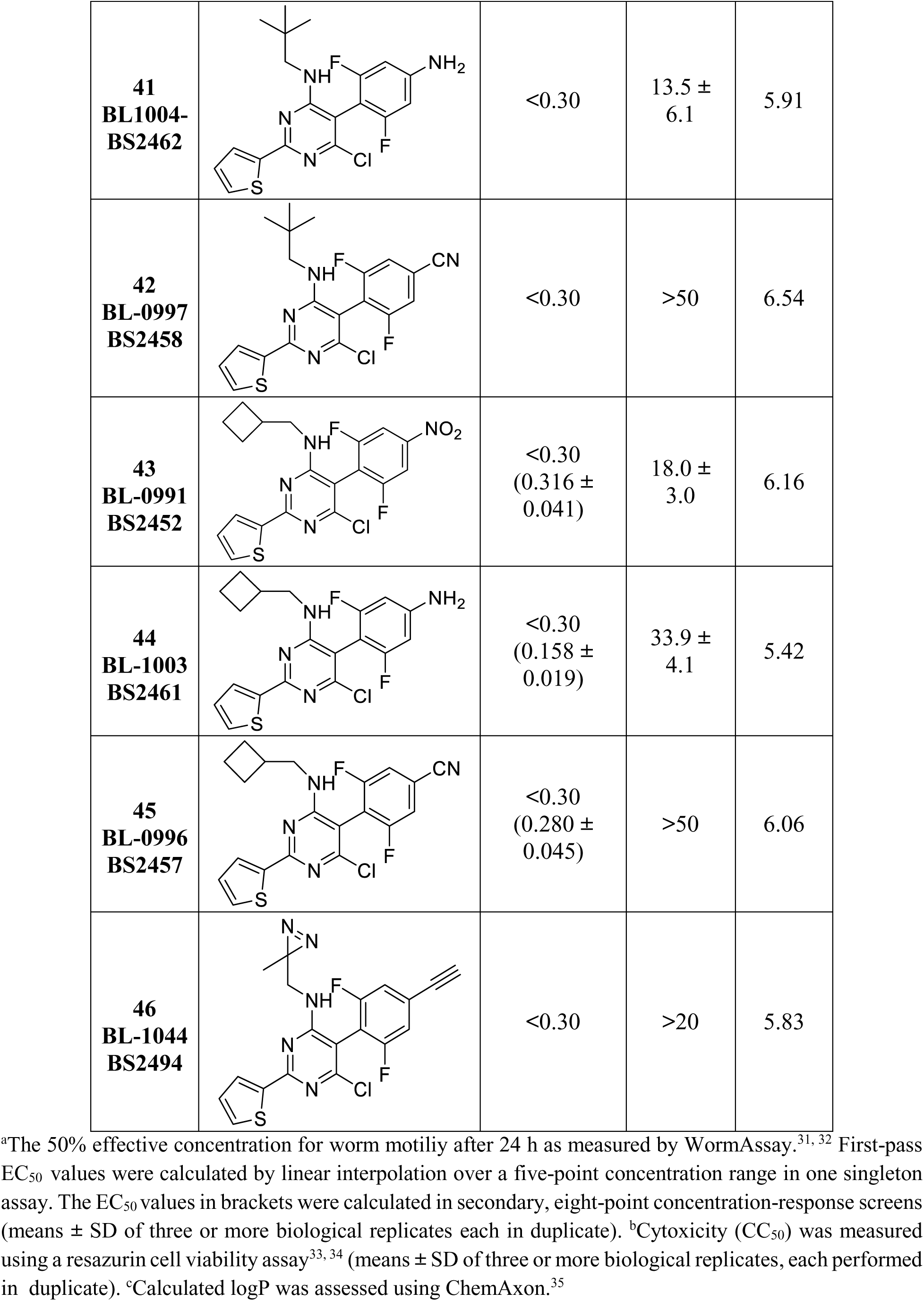
Bioactivities of TPPs against adult *S. mansoni*, cytotoxicity *vs*. HEK293 cells, and calculated logP values.

| Cpd | Structure | WormAssay<br>EC<sub>50</sub> (μM)<sup>a</sup> | HEK293<br>CC<sub>50</sub><br>(μM)<sup>b</sup> | cLogP<sup>c</sup> |
| --- | --- | --- | --- | --- |
| **3**<br>**BL-0674**<br>**BS1398** |  | 0.49<br>(0.538 ± 0.075) | >50 | 6.81 |
| **4**<br>**BL-0932**<br>**BS2271** |  | 5.0 | >50 | 6.66 |
| **5**<br>**BL-0933**<br>**BS2272** |  | 0.59 | >50 | 6.52 |
| **6**<br>**BL-0935**<br>**BS2274** |  | 5.0 | >50 | 6.52 |
| **7**<br>**BL-0936**<br>**BS2275** |  | 0.71 | >50 | 6.66 |
| **8**<br>**BL-0937**<br>**BS2276** |  | 1.86 | >50 | 6.66 |
| **9**<br>**BL-0929**<br>**BS2268** | | 0.65 | $21.0 \pm 3.8$ | 6.38 |
| **10**<br>**BL-0934**<br>**BS2273** | | <0.30 | $34.3 \pm 12.4$ | 6.47 |
| **11**<br>**BL-0947**<br>**BS2280** | | >5 | $1.76 \pm 0.67$ | 6.13 |
| **12**<br>**BL-0946**<br>**BS2279** | | <0.30 | $34.0 \pm 4.3$ | 5.86 |
| **13**<br>**BL-0945**<br>**BS2278** | | <0.30 | $20.2 \pm 1.3$ | 6.62 |
| **14**<br>**BL-0995**<br>**BS2456** |  | 1.02 | >50 | 6.52 |
| **15**<br>**BL-0923**<br>**BS2263** | | >5 | $31.6 \pm 4.9$ | 4.84 |
| **16**<br>**BL-0931**<br>**BS2270** |  | >5 | >50 | 6.14 |
| **17**<br>**BL-0920**<br>**BS2260** |  | <0.30 | 34.8 ± 6.0 | 6.11 |
| **18**<br>**BL-0913**<br>**BS2253** |  | >5 | >50 | 5.61 |
| **19**<br>**BL-0915**<br>**BS2255** |  | 0.55 | >50 | 6.03 |
| **20**<br>**BL-0916**<br>**BS2256** |  | 0.78 | >50 | 6.35 |
| **21**<br>**BL-0993**<br>**BS2454** |  | 0.50 | >50 | 6.36 |
| **22**<br>**BL-0907**<br>**BS2248** |  | 1.82 | 14.4 ± 1.7 | 4.94 |
| **23**<br>**BL-0912**<br>**BS2252** | | 1.24 | $5.55 \pm 0.77$ | 5.40 |
| **24**<br>**BL-0914**<br>**BS2254** | | >5 | $28.2 \pm 2.7$ | 6.20 |
| **25**<br>**BL-0917**<br>**BS2257** |  | 5.0 | >50 | 6.62 |
| **26**<br>**BL-0918**<br>**BS2258** | | 2.13 | $24.0 \pm 2.8$ | 7.04 |
| **27**<br>**BL-0909**<br>**BS2250** | | >5 | $36.7 \pm 1.9$ | 5.78 |
| **28**<br>**BL-0908**<br>**BS2249** | | 1.96 | $22.7 \pm 1.9$ | 5.53 |
| **29**<br>**BL-0906**<br>**BS2247** |  | 0.91 | >50 | 5.60 |
| **30**<br>**BL-0919**<br>**BS2259** |  | <0.30 | 40.1 ± 4.6 | 6.83 |
| **31**<br>**BL-0926**<br>**BS2265** |  | 0.64 | >50 | 6.90 |
| **32**<br>**BL-0927**<br>**BS2266** |  | 4.52 | 16.9 ± 2.7 | 5.67 |
| **33**<br>**BL-0791**<br>**BS1948** |  | 0.61 | 10.4 ± 1.6 | 3.95 |
| **34**<br>**BL-0795**<br>**BS1950** |  | 3.54 | 34.9 ± 6.6 | 4.44 |
| **35**<br>**BL-1050**<br>**BS2521** |  | <0.30 | 8.47 ± 3.44 | 4.98 |
| **36**<br>**BL-1049**<br>**BS2520** |  | <0.30<br>(0.038 ± 0.006) | ~50 | 5.06 |
| **37**<br>**BL-1001**<br>**BS2459** |  | 2.14<br>(3.09 ± 0.59) | >50 | 5.21 |
| **38**<br>**BL-1005**<br>**BS2463** |  | <0.30<br>(0.037 ± 0.006) | 37.3 ± 6.7 | 4.48 |
| **39**<br>**BL-1002**<br>**BS2460** |  | 1.50 | 23.7 ± 5.2 | 5.11 |
| **40**<br>**BL-0990**<br>**BS2451** |  | <0.30 | >50 | 6.64 |
| **41**<br>**BL1004-**<br>**BS2462** |  | <0.30 | 13.5 ± 6.1 | 5.91 |
| **42**<br>**BL-0997**<br>**BS2458** |  | <0.30 | >50 | 6.54 |
| **43**<br>**BL-0991**<br>**BS2452** |  | <0.30<br>(0.316 ± 0.041) | 18.0 ± 3.0 | 6.16 |
| **44**<br>**BL-1003**<br>**BS2461** |  | <0.30<br>(0.158 ± 0.019) | 33.9 ± 4.1 | 5.42 |
| **45**<br>**BL-0996**<br>**BS2457** |  | <0.30<br>(0.280 ± 0.045) | >50 | 6.06 |
| **46**<br>**BL-1044**<br>**BS2494** |  | <0.30 | >20 | 5.83 |

<sup>a</sup>The 50% effective concentration for worm motility after 24 h as measured by WormAssay.<sup>31, 32</sup> First-pass EC<sub>50</sub> values were calculated by linear interpolation over a five-point concentration range in one singleton assay. The EC<sub>50</sub> values in brackets were calculated in secondary, eight-point concentration-response screens (means ± SD of three or more biological replicates each in duplicate). <sup>b</sup>Cytotoxicity (CC<sub>50</sub>) was measured
using a resazurin cell viability assay<sup>33, 34</sup> (means ± SD of three or more biological replicates, each performed in duplicate). °Calculated logP was assessed using ChemAxon.<sup>35</sup>

## 2. Results and Discussion

### 2.1. Design & synthesis of TPP analogs

The thiophene moiety at C2 is required for worm paralysis even though it contributes to the lipophilicity of the TPPs.^13^ Nonetheless, SAR studies conducted with MT-active phenylpyrimidines and triazolopyrimidines showed that, in general, the nature of the amine fragment at C6 and the choice of substituents at the phenyl ring at C5 are important determinants of absorption, distribution, metabolism, and excretion, and pharmacokinetics (ADME-PK).^15, 23, 24^ Thus, the primary focus of the present SAR studies was to evaluate structural modifications at the fragments linked at C5 and C6 of the TPP scaffold with the goal of identifying potent derivatives of **3**. In parallel, we assessed the SPR (*i.e.,* cLogP) to guide the selection of analogs with lipophilicities that may be conducive to acceptable ADME-PK.

The second objective of our study was to design, synthesize and evaluate a TPP probe that could be used in whole-organism PAL studies to investigate the mechanism of action of this class of compounds. Our previous report regarding TPPs *vs. S. mansoni* showed that a diazirine-containing amine fragment at C6 is tolerated without significant loss of paralytic potency.^13^ Starting from these initial results, we envisaged a TPP probe equipped with both a photoactivatable diazirine moiety, as well as an acetylene group, which could be used as a click handle^25, 26^ after the post-photolabeling step to attach an appropriate fluorescent and/or affinity reporter.

The syntheses of the TPP congeners, **4**–**46**, are summarized in Schemes **1**–**4**. Substituted malonates **47**–**55** (Scheme 1) were prepared *via* Ullmann-type cross coupling with the appropriate brominated fluorobenzenes. For electron-poor phenyl rings, the substituted malonates, **54** and **55**, were synthesized via nucleophilic aromatic substitution of diethyl malonate and various difluoro phenyls. The substituted diethylmalonates were then subjected to a cyclocondensation reaction with the thiophene-2-carboximidamide hydrochloride in the presence of DBU to form the 4,6-dihydroxy TPP intermediates, **56**–**64**. Next, a chlorination reaction of the previously obtained bis-phenols in the presence of phosphoryl oxychloride provided the corresponding TPP dichlorides, **65**–**73** (Scheme 1).

**Scheme 1.**
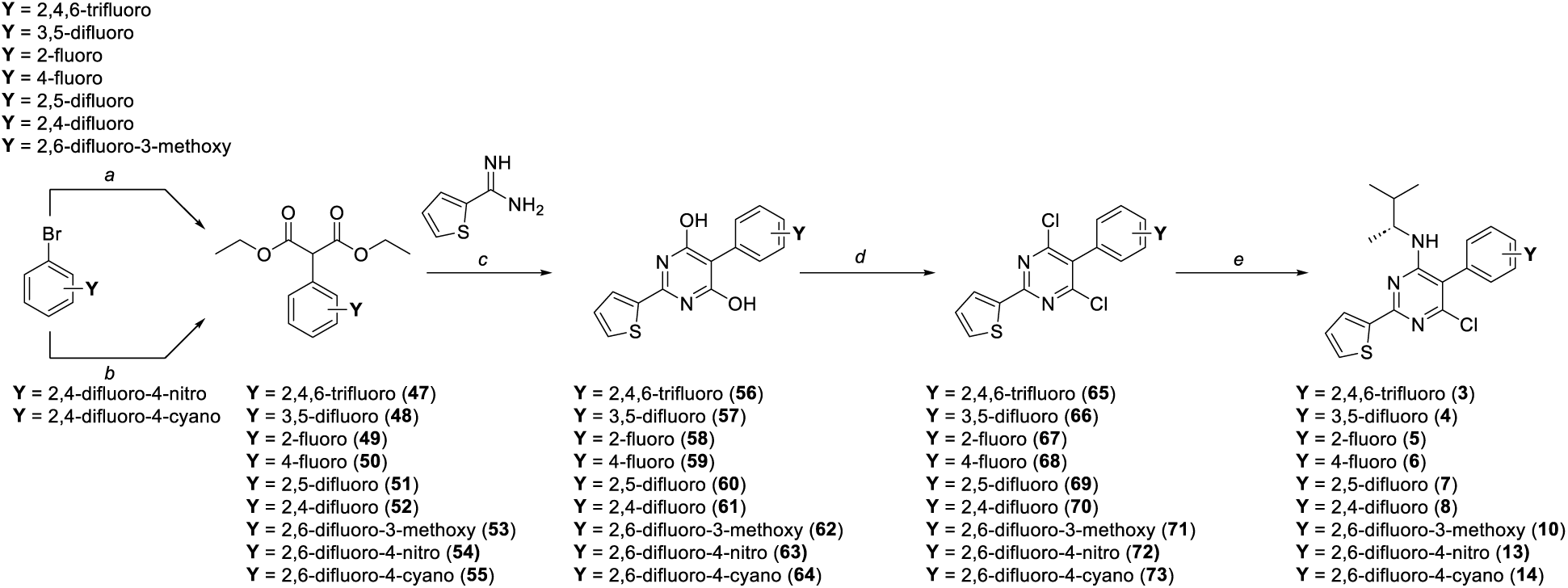
*Reagents and conditions:* (a) diethylmalonate, NaH, appropriately substituted bromobenzene, CuBr, 1,4-dioxane, 60 to 100 °C, 16 h, 58–72%. (b) diethylmalonate, NaH, appropriately substituted fluorobenzene, THF, 0 °C to rt, 2 h, 53–61%. (c) appropriate substituted malonate, thiophene-2-carboximidamide hydrochloride, DBU, NMP, rt to 95 °C, 5 h, 33–94%. (d) appropriate 5-substituted-2-(thiophen-2-yl)pyrimidine-4,6-diol, POCl_3_, 100 °C, 16 h, 40–73%. (e) appropriate 5-substituted-2-(thiophen-2-yl)pyrimidine-4,6-dichloride, (*R*)-3-methylbutan-2-amine, DMF, rt, 16 h, 45–93%.

Displacement of the chloro substituent at C6 with (*R*)-3-methylbutan-2-amine furnished the TPPs **3**, **4**–**8**, **10**, **13** and **14**. Congeners that bear different amines at C6 and *para-*phenyl substitutions at C5 were synthesized via substitution of the chloro substituent in **65**, **71**, **72** or **73** with a variety of amines, which led to the formation of the TPPs **15**–**34**, **36**, **37**, **39**, **40**, **42**, **43** and **45** (Scheme 2).

**Scheme 2.**
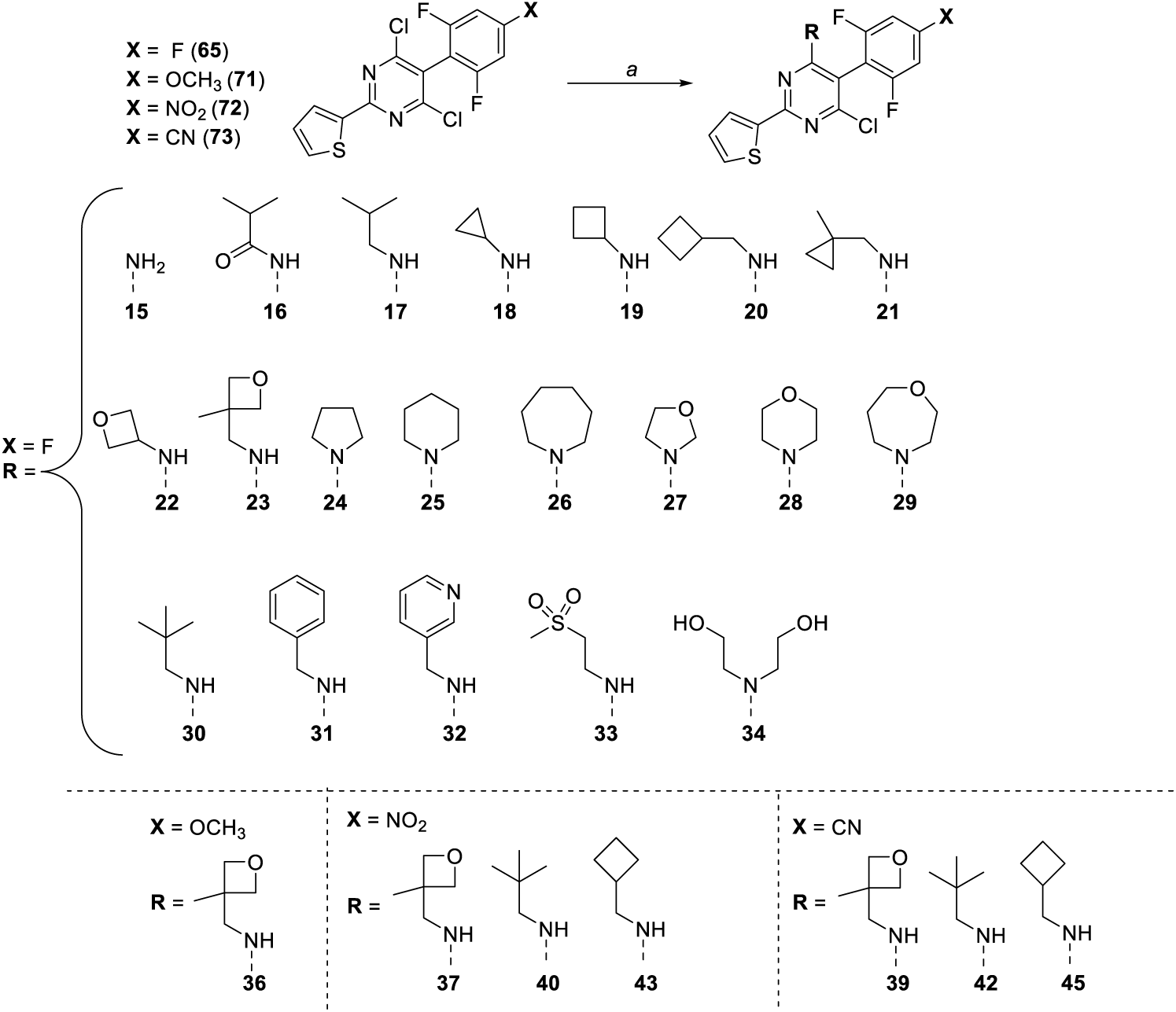
*Reagents and conditions*: (a) appropriate 4,6-dichloro-2-(thiophen-2-yl)-5-(2,6-difluorophenyl, 4-substituted) pyrimidine, appropriate amine, Et_3_N, DMF, rt, 16 h, 36–94%.

Compounds **9** and **35**, which feature a 2,6-difluoro,4-hydroxyphenyl ring at C5, were obtained via boron tribromide-mediated demethylation of the corresponding 4-methoxy analogs, **10** and **36** (Scheme 3). A TPP analog with an alkoxy sidechain at the *para*-phenyl ring at C5 was synthesized via O-alkylation of **9** with the *N*-Boc-protected chloropropanamine, which then led to the formation of **74**. Further Boc-deprotection of the terminal amine generated **11**. Compounds with the 6-difluoro, 4-aniline ring at C5 were synthesized from the 2,6-difluoro, 4-nitro analogs, **13**, **37**, **40** and **43**, via reduction of the nitro group to the corresponding aniline (**12, 38**, **41** and **44**).

**Scheme 3.**
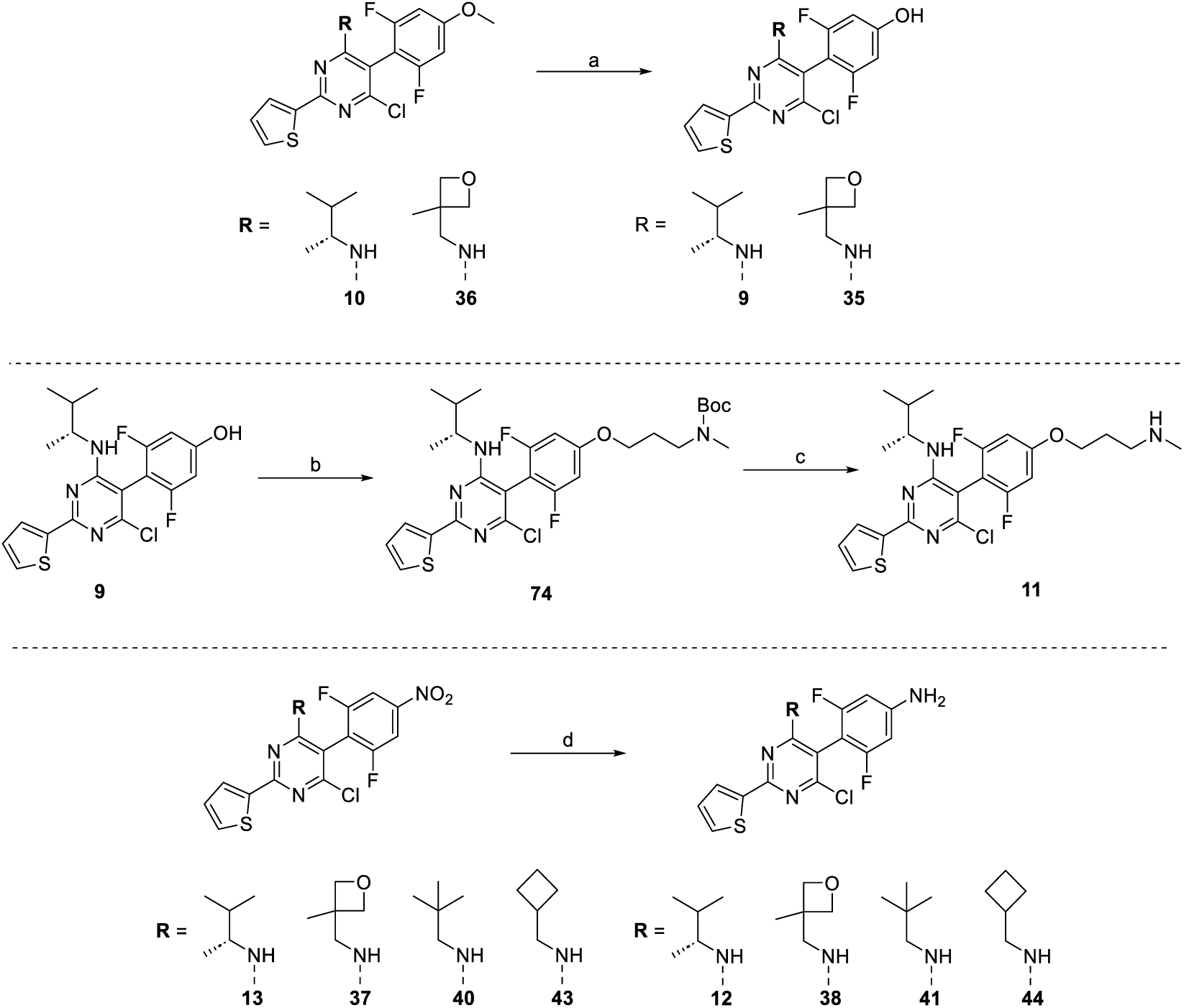
*Reagents and conditions:* (a) BBr_3_, CH_2_Cl_2_, −78 °C to rt, 16 h, 90%. (b) 3-chloro-*N,N-*dimethylpropan-1-amine hydrochloride or tert-butyl (3-chloropropyl)(methyl)carbamate, K_2_CO_3_, DMF, rt, 16-72 h, 83–91%. (c) HCl (4.0 M), 1,4-dioxane, rt, 16 h, 87%. (d) Fe, NH_4_Cl, H_2_O/CH_3_OH (4:5), 80 °C, 2 h, 54–86%.

The TPP-based PAL probe was synthesized as summarized in Scheme 4. Starting with the *para-*iodo malonate, **75**,^24^ the thiophenyl pyrimidine was formed via cyclocondensation with the thiophenyl amidine, followed by chlorination with phosphoryl oxychloride to form **76**. The diazirine amine, **77**, was synthesized as previously described,^27^ and then used to displace the chloro substituent at C6 to give **78**. Finally, Sonogashira coupling^28^ of **78** with TMS-acetylene, followed by deprotection of the TMS group, furnished the PAL probe, **46**.

**Scheme 4.**
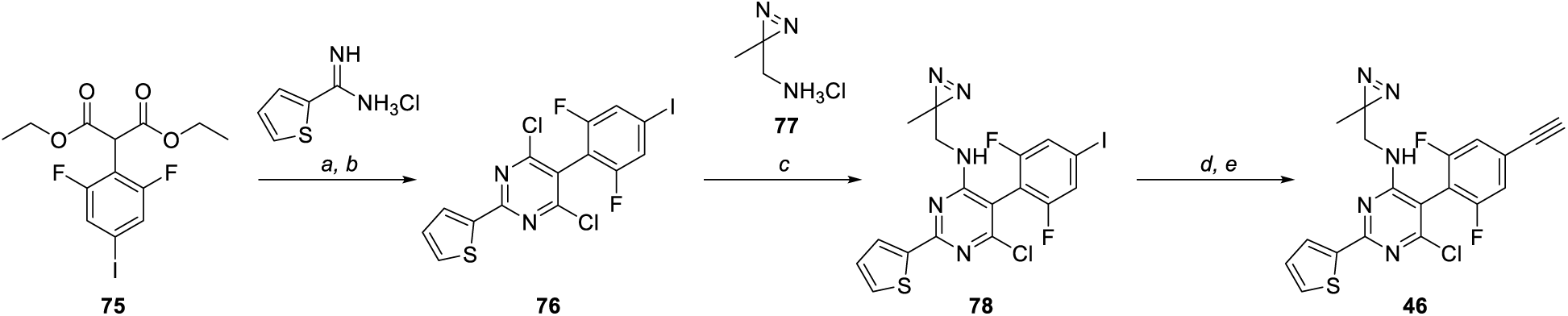
*Reagents and conditions:* (a) **75**, thiophene-2-carboximidamide hydrochloride, DBU, NMP, 95 °C, 5 h, 83%. (b) 5-(2,6-difluoro-4-iodophenyl)-2-(thiophen-2-yl)pyrimidine-4,6-diol, POCl_3_, 100 °C, 16 h, 64%. (c) **76**, **77**, Et_3_N, DMF, rt, 1 h, 74%. (d) **78**, trimethylsilylacetylene, Pd(Ph_3_P)_4_, CuI, Et_3_N, DMF, rt, 2 h, 43%. (e) 6-Chloro-5-(2,6-difluoro-4-((trimethylsilyl)ethynyl)phenyl)-*N*-((3-methyl-3*H*-diazirin-3-yl)methyl)-2-(thiophen-2-yl)pyrimidin-4-amine, K_2_CO_3_, CH_3_OH, rt, 1 h, 79%.

### 2.2. Structure Activity Relationships

Compounds were tested in first-pass screens over five concentrations (0.3 – 5 µM) for 24 h in a previously described phenotypic assay involving *ex vivo* adult (≥ 42-day-old) *S. mansoni* worms.^29, 30^ WormAssay^31, 32^ was used to measure average worm motility which was expressed as an EC_50_ value, *i.e*., the value that corresponds to a decrease of 50% in worm motility relative to DMSO control. Counter-toxicity screens employed HEK293 cells^13^ with up to 50 µM compound, and lipophilicity was assessed using the clogP. Compounds that combined paralytic potency, low HEK293 toxicity and decreased lipophilicity relative to the starting TPP, **3**, were tested in secondary eight-point concentration screens (0.01 – 5 µM) of adult *S. mansoni* to define a final EC_50_ value. The data are summarized in Table 1.

Compared to the parent compound, **3**, analogs that bear different fluorination patterns at the C5 phenyl ring (**4**–**8**) indicate that fluorinations in the *ortho* position are preferred for activity. This trend is evident in that a single fluorination in *ortho* (**5**) is sufficient to retain activity (EC_50_ = 0.59 µM, *i.e*., comparable to that of **3** (EC_50_ = 0.54 µM)), whereas TPPs that are fluorinated only in *meta* (**4**) or *para* (**6**) were less potent (EC_50_ = 5.0 µM). In addition, the difluorinated derivatives, **7** and **8**, which feature a combination of *ortho*- and either *meta*- or *para*-fluorination, respectively, retain paralytic activity (0.71 µM and 1.86 µM, respectively), albeit in the case of **8**, a decrease relative to **3**. Examining the SPR, changing the fluorination patterns at the phenyl ring did not substantially alter lipophilicity with cLogP values being maintained in a narrow range of 6.52 to 6.66. Notably, and regardless of the fluorination pattern, no toxicity of HEK293 cells was measured up to 50 µM, the highest concentration tested.

Next, the data for **9** – **14**, in which the *para* fluoro substituent of the C5 phenyl ring is replaced by substituents of different stereo-electronic characteristics, suggest that electron-withdrawing (*e.g*., OMe (**10**) and NH_2_ (**12**)) or -donating groups (**13**) seem not to be important for activity as steric effects. In either case, paralysis was more potent (EC_50_ < 0.30 µM) compared to **3**. In contrast, **11**, which features a relatively large alkoxy side chain substituent, did not cause paralysis (EC_50_ > 5 µM) and was cytotoxic with a CC_50_ value of 1.76 µM. This suggests that larger groups at this position are undesirable. By varying the electronic properties and polarity of substituents at this position (**9** – **14**, Table 1), we observed a broader range of cLogP values (5.86–6.47), reflecting the impact of these modifications on the SPR. The above *para-*phenyl SAR data provided the basis for the design of a photoactivatable TPP probe (**46**) which incorporates an alkyne click handle in the *para* position without loss of paralytic activity (EC_50_ < 0.30).

Consistent with our earlier study,^13^ the assessment of analogs bearing different amine fragments at the C6 position indicated that the presence of an aliphatic amine fragment is necessary for TPP paralytic activity (Table 1). Indeed, *N*-dealkylation of **3** (**15**) resulted in a loss of paralysis. Similarly, replacement of the (*R*)-3-methylbutan-2-amine of **3** with the isobutyramide (**16**) also generated an inactive compound. Conversely, the presence of the isobutylamine (**17**, EC_50_ < 0.30 µM) at C6 was well-tolerated resulting in a compound with activity better than **3**. Although smaller amine fragments, such as the cyclopropylamine (**18**), appeared to be devoid of activity, homologation of the cyclopropyl group to the corresponding cyclobutyl ring (**19**) recovered activity (EC_50_ = 0.55 µM). Likewise, incrementally larger amine fragments, such as cyclobutylmethanamine (**20,** EC_50_ = 0.78 µM), (1-methylcyclopropyl)methanamine (**21**, EC_50_ = 0.50 µM), 2,2-dimethylpropan-1-amine (**30**, EC_50_ < 0.30 µM) and benzylamine (**31**, EC_50_ = 0.64 µM) exhibited activity comparable to or better than **3**.

The evaluation of a series of homologs bearing cyclic amines at C6 demonstrated similar general trends with the pyrrolidine derivative (**24**, EC_50_ > 5 µM) being inactive compared to the corresponding and modestly active TPPs bearing either a piperidine (**25**, EC_50_ = 5.0 µM) or an azepane (**26**, EC_50_ = 2.13 µM) (Table 1). Interestingly, although these results suggest that the presence of a relatively bulky amine may be generally preferred for activity, derivatives with polar amine fragments exhibited a wide range of potencies, including weak (EC_50_ > 4 µM), moderate (EC_50_ between 1 and 4 µM) and potent (EC_50_ < 1 µM) activity. These include analogs bearing hydrogen bond donor (**34**) or acceptor (**22**, **23**, **28**, **29**, **32**, **33**) groups, which reduce compound lipophilicity.

Additional derivatives (**35**–**45**) were synthesized by combining the most promising amine residues at C6 with the most promising substituted *ortho, ortho-*difluorophenyl fragments at C5 (Table 1). Several of these compounds, **40**-**45**, were more potent than **3**, including in secondary, eight-point concentration response analyses. Some of these, however, *e.g.,* **35**, **41** and **43**, generated modest cytotoxicity to HEK293 cells with CC_50_ values < 20 µM. Importantly, all of the compounds from this set decreased clogP relative to the reference compound **3** (logP = 6.81), with values ranging from 4.48 to 6.64.

Due to their potent paralysis, relatively low cytotoxicity and the decreased lipophilicity compared to **3**, the TPP congeners **36**–**38** and **43**–**45** were selected for secondary, eight-point concentration response analyses (Table 1). Among these, **36** and **38** were especially promising with EC_50_ values of 38 and 37 nM, respectively, and high selectivity indices (SIs) relative to HEK293 cells of 895 and 1,008, respectively. Expanded cytotoxicity tests for **3** and **38** with HeLa and HepG2 cells also demonstrated comparatively low CC_50_ values > 20 µM (Table 4). Also, unlike **3**, both **36** and **38** possess more drug-like physicochemical properties,^21^ with **38** presenting no violations of the Lipinski’s Rule of Five,^20^ a >2 log-unit reduction in clogP and significantly increased solubility (46 µM *vs.* < 0.5 µM, Table 4).

2.3 *PK parameters of **38***.

Compounds **3**, **36** and **38** were administered IP at 2.5 mg/kg to a group of three mice in a cassette dosing format^36^ to determine plasma half-life (t_1/2_) and the maximum plasma concentration (C_max_), two key parameters that are used to gauge the *in vivo* performance of compounds. Compared to the progenitor compound, **3**, compound **38** offered a longer t_1/2_ (1.51 *vs* 0.48 h) for some loss of C_max_ (247.3 *vs* 399.3 ng/mL), whereas **36** was below the limit of quantitation (LQQ; Table 2).

**Table 2:** Plasma half-life and the maximum plasma concentration for selected TPPs.

| COMPOUND | PARAMETER |  |  |  |
| --- | --- | --- | --- | --- |
|  | C<sub>max</sub> (ng/mL) |  | t<sub>1/2</sub> (h) |  |
|  | Mean | CV% | Mean | CV% |
| **3** | 399.33 | 5.72 | 0.48 | 18.0 |
| **38** | 247.3 | 14.2 | 1.51 | 16.8 |
| **36** | LOQ | LOQ | LOQ | LOQ |

### 2.4 Compound **38** induces a sustained paralysis of adult and juvenile S. mansoni

The most promising congener, **38**, was then tested in washout assays to assess the reversibility of the paralytic effect against adult and juvenile (21 – 28-day-old) *S. mansoni* using both WormAssay^31, 32^ and observational analysis.^29, 30^ The choice of concentration (0.6 µM) and washout time (2 h) to perform these assays with **38** was guided by the respective PK parameters, *i.e*., the maximum plasma concentration (C_max_, converting ng/mL to µM) and the plasma half-life (t_1/2_; Table 2). We also performed the washout assays at double the concentration (1.2 µM). For comparison, the current antischistosomal drug, R/S-PZQ, was tested in parallel at the same concentrations. *In vivo,* PZQ is essentially inactive against juvenile (21 – 28-day-old) *S. mansoni*,^8, 37, 38^ and this deficiency is considered to contribute to its variable efficacy in the treatment of human disease.^39, 40^ Thus, a competitive alternative preclinical candidate should be effective against both adult and juvenile stages of the parasite.^12, 38^

Worms were incubated with **38** or PZQ for 2 h, and then washed five times in medium with a subsequent medium exchange at the 72-h time point. At the 2 h time point, and prior to washing, both **38** and PZQ had decreased adult worm motility to between 0 and 10% of the DMSO control (Figure 2A). For juveniles, motility was reduced to 5 – 25% and 0 – 10% of control by **38** and PZQ, respectively (Figure 2B). Observationally, for both developmental stages, paralysis by **38** at the 2 h time point was as described^13^ whereby worm movement and flexing were severely curtailed with the inability of either the oral or ventral sucker to grasp the floor of the well. Otherwise, the worms looked normal. PZQ, in contrast, and consistent with previous reports,^8, 29, 41, 42^ caused a spastic paralysis of both developmental stages involving a severe contraction in body length, an inability of the suckers to adhere to the well floor, and blebbing of the parasite surface (tegument). After washout, worm motility was measured by WormAssay and observed under the microscope, each day or every other day, for up to 7 or 9 days.

**Figure 2.**
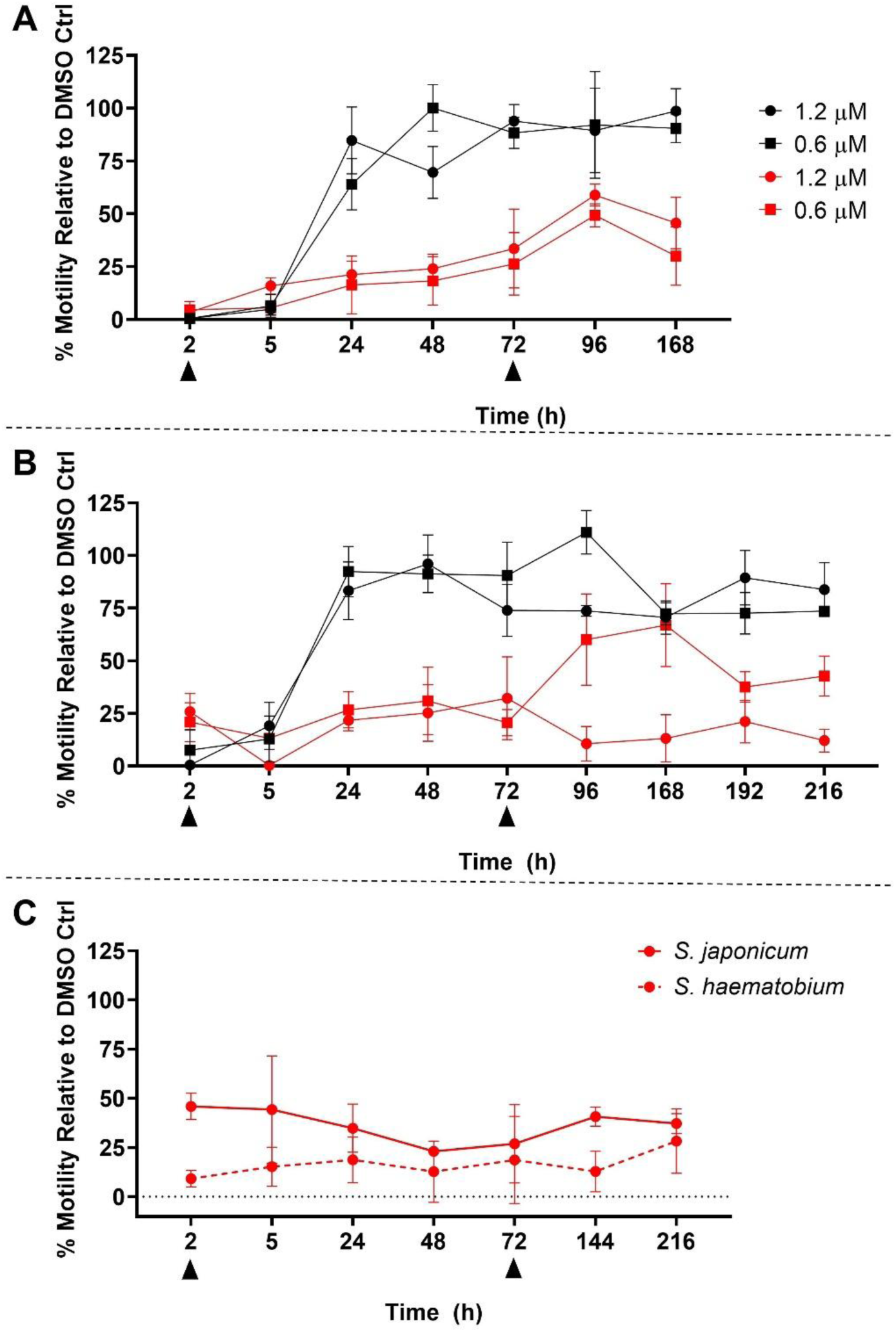
Compound **38** induces a sustained paralysis in schistosomes. *Ex vivo* worms were exposed to either **38** (red) or PZQ (black) for 2 h at 1.2 (circles) or 0.6 µM (squares). After 2 and 72 h, worms were washed five times and once, respectively, as indicated by the arrowheads. Motility as a function of time for (**A**) adult *S. mansoni*, (**B**) juvenile *S. mansoni*, and (**C**) adult *S. japonicum* and *S. haematobium*. For *S. mansoni,* two biological assays each in triplicate were performed; for *S. japonicum* one assay in triplicate was performed, and for *S. haematobium,* three biological assays in at least duplicate were performed (means and SD values shown).

For adult *S. mansoni*, recovery of motility after washout of 1.2 and 0.6 µM **38** ranged between 15 and 60% of the DMSO control over the 9-day incubation period (Figure 2A). Under the microscope, worms at the 9-day time point displayed constrained and repetitive slow movements with both suckers being non-functional. In contrast, 24 h after washout of PZQ, adults had recovered between 65 and 85% of DMSO control motility, with essentially 75 to 100% recovery thereafter up to Day 7 (Figure 2A). Under the microscope on day 7, these worms appeared similar in shape and motility to DMSO controls, including sucker functionality and a seemingly normal surface tegument.

For *S. mansoni* juveniles exposed to 1.2 and 0.6 µM **38** for 2 h, motility after washout was generally < 25% of DMSO control at 1.2 µM, and between 25 and 70% of control at 0.6 µM across the 9-day incubation period (Figure 2B). Observationally, on Day 9, worms presented similar constrained and repetitive slow movements as noted for adults. In contrast, juveniles exposed to PZQ at both concentrations regained between 80 and 95% of DMSO control motility 24 h after washout with 75 to 100% of control motility remaining for the duration of the 9-day assay. Under the microscope, these juveniles appeared similar in shape and motility to DMSO controls, with a normal surface tegument and both suckers being able to grasp the well floor.

Overall, for both adult and juvenile *S. mansoni,* a 2 h pulse with 1.2 and 0.6 µM **38** induced a sustained paralysis as measured quantitatively and assessed observationally. This paralysis contrasts with the seeming full recovery made by both stages of *S. mansoni* after exposure to PZQ under similar conditions.

### 2.5 Compound **38** produces a sustained paralysis of other medically important schistosome species

The desired preclinical candidate profile for a new anti-schistosomal compound includes activity against the two other medically important species of schistosomes, namely *S. haematobium,* which is prevalent in sub-Saharan Africa, and *S. japonicum,* which is located in parts of east and South-East Asia. The availability of both these species is more limited so experiments were performed with one concentration (1.2 µM) of **38**.

At the 2 h time point in the presence of **38**, *S*. *japonicum* motility was 45% of DMSO control (Figure 2C). After washout, motility remained essentially unchanged out to 9 days. Qualitatively on day 9, the worms presented the same phenotypic abnormalities in movement and sucker function as noted for S*. mansoni*.

*S. haematobium* adults seemed particularly susceptible to **38**. Motility was decreased to ∼10% of control before and after washout (Figure 2C). Under microscope on Day 9, the phenotypic abnormalities were as noted for *S. mansoni*. Overall, the data suggest that the TPP series, as represented by **38**, has pan-antischistosomal activity.

### 2.3. Photoaffinity Labeling

To identify possible protein targets of **38**, and guided by the SAR analysis, we designed and synthesized a PAL probe, **46**, and applied a workflow we previously described for *Trypanosoma brucei* ^43^ to *ex vivo* adult *S. mansoni* worms.^44^ The probe features a photoactivatable alkyl diazirine group at C4 and an alkyne click handle at the C5 *para-*phenyl position (Figure 3A). Evaluation of **46** in the worm motility assay confirmed that the compound paralyzes *S. mansoni* with an EC_50_ value of < 0.30 µM (Table 1). In addition to **46**, a triazolopyrimidine PAL probe, **79** (Figure 3A), which binds to tubulin in HEK293 cells, *Trypanosoma brucei* parasites,^43^ and *S. mansoni,* but does not cause paralysis in *S. mansoni* (not shown),^44^ was also tested.

**Figure 3.**
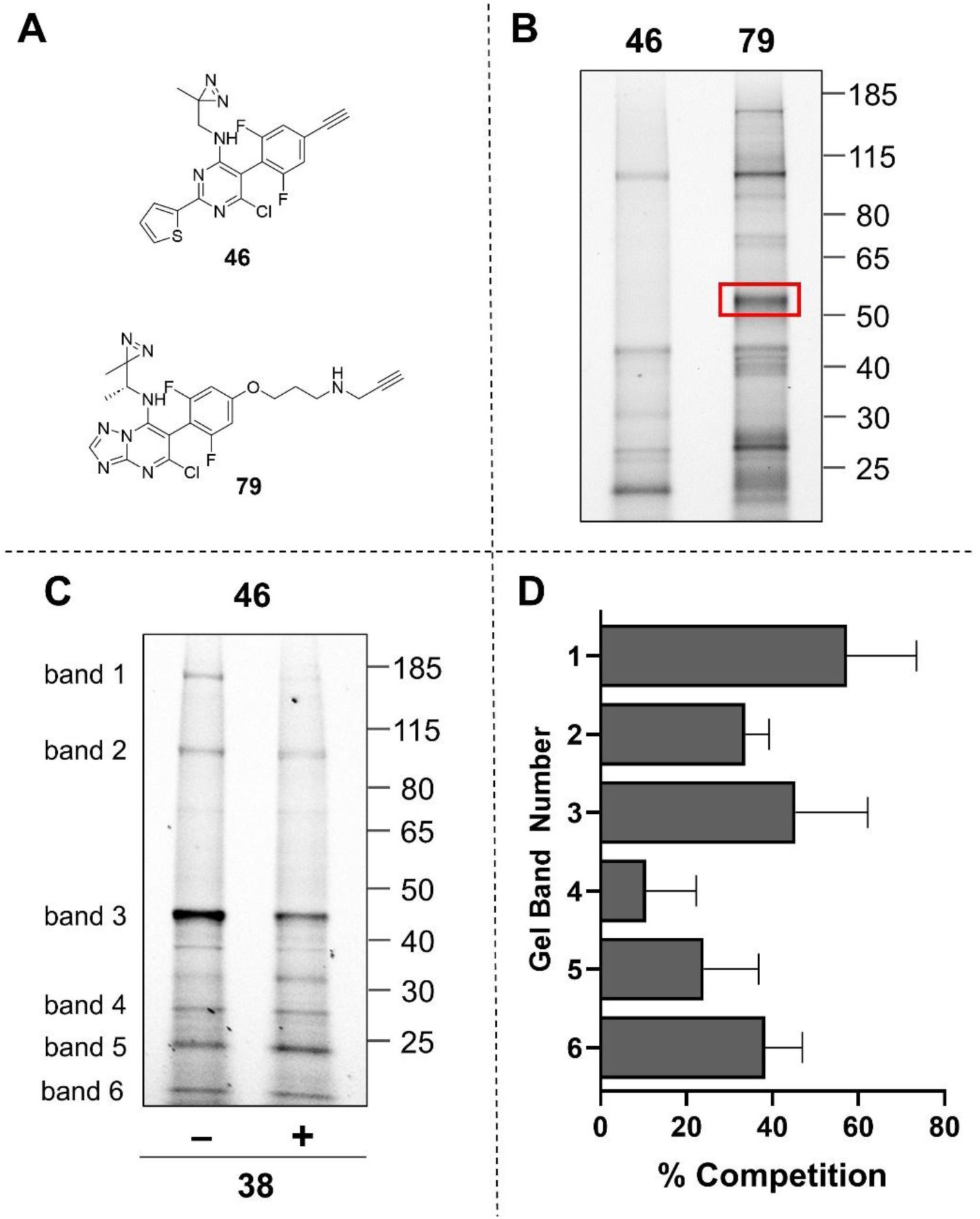
Application of PAL to identify possible protein targets of the TPP series in *S. mansoni*. (**A**) Structures of the TPP and a triazolopyrimidine PAL probes, **46** and **79**, respectively. (**B**) Fluorescence analyses of a SDS-PAGE gel after labelling with **46** or **79**. Proteomics analysis of the protein band labeled specifically by **79** (red box) identifies *S. mansoni* tubulin. (**C**) Fluorescence analyses of SDS-PAGE gel after labeling with **46** in the absence or presence of 25X of the competitor TPP, **38**. (**D**) Decreases in the densitometry signals of each of the six bands labelled by **46** in the presence of the competitor, **38**. For **B** and **C**, sample concentrations were measured using the BCA assay and normalized to 0.3 µg/mL to ensure equal protein loading. For **D**, data shown are the means ± SD from three biological replicates.

After incubation of *S. mansoni* adults with **46** or **79** and subsequent photoactivation, worm lysates were homogenized and subjected to the click reaction^25, 26^ to install a dual fluorescent (TAMRA) and affinity (biotin) reporter. Biotinylated proteins were enriched via neutravidin-affinity pulldown, separated by SDS-PAGE, and visualized by in-gel fluorescence. Fluorescent bands of interest were excised and analyzed by mass spectrometry-based proteomics.

SDS-PAGE fluorescence analysis revealed several bands that appear to be identically labeled by both PAL probes, **46** and **79**, and others that are differentially labeled (Figure 3B). The triazolopyrimidine probe, **79**, labeled a prominent 50 kDa protein band that was previously confirmed by proteomics analysis to be *S. mansoni* tubulin^44^ In contrast, the TPP probe, **46**, did not label tubulin. This result, and the finding that **79** does not paralyze *S. mansoni* (not shown), suggest that worm paralysis by the TPPs is not tubulin-dependent.

Next, to distinguish between specific and non-specific binding of the TPP probe, **46**, a competition experiment was conducted using the photostable TPP, **38**. *Ex vivo* adult worms were pre-incubated for 30 min with the competitor in a 25X excess of the PAL probe concentration, followed by coincubation with the PAL probe and subsequent photoactivation. After worm lysis, protein enrichment by neutravidin-affinity pulldown, the bands of interest were excised and analyzed by proteomics. Six protein bands were diminished in intensity relative to the sample in the absence of competition as measured by fluorescence-based densitometry (Figure 3C and D). The identities of the most abundant proteins found within each band are shown in Table 3.

**Table 3.**
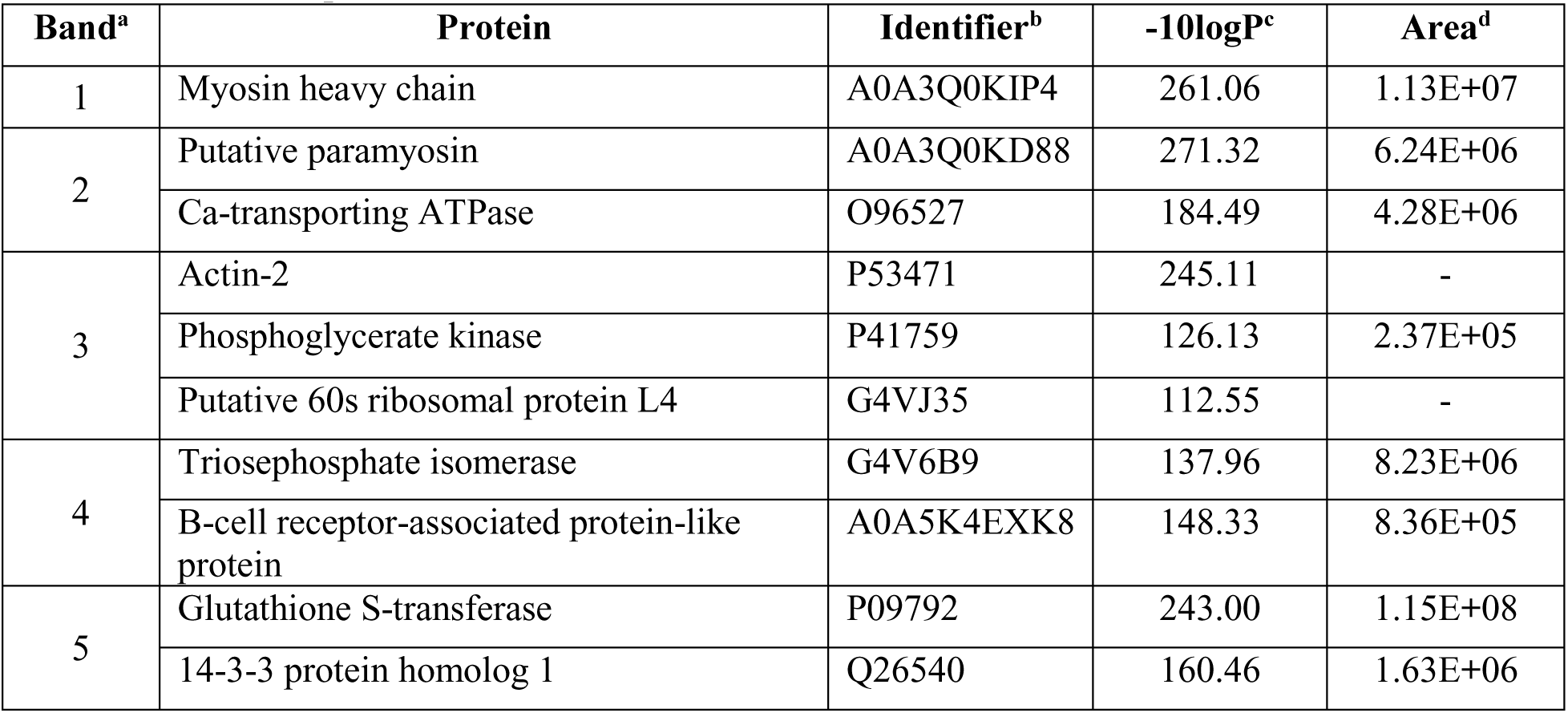
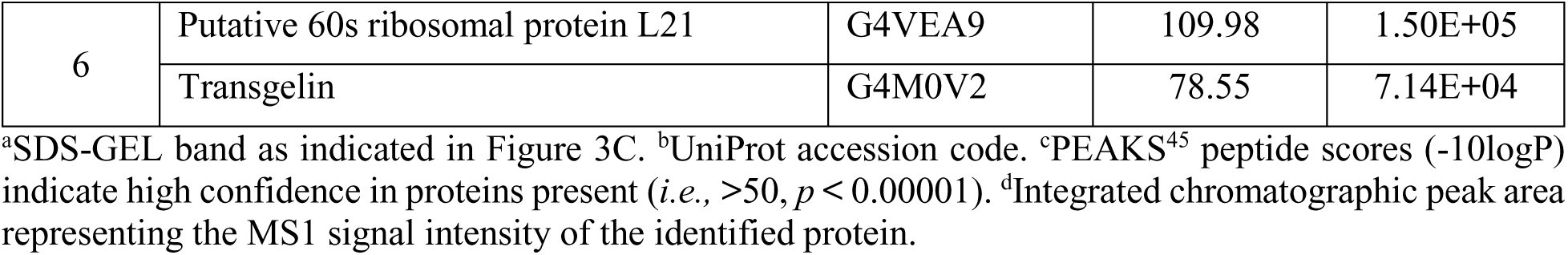
***S. mansoni* proteins enriched by PAL probe 46**

| Band<sup>a</sup> | Protein | Identifier<sup>b</sup> | -10logP<sup>c</sup> | Area<sup>d</sup> |
| --- | --- | --- | --- | --- |
| 1 | Myosin heavy chain | A0A3Q0KIP4 | 261.06 | 1.13E+07 |
| 2 | Putative paramyosin | A0A3Q0KD88 | 271.32 | 6.24E+06 |
|  | Ca-transporting ATPase | O96527 | 184.49 | 4.28E+06 |
| 3 | Actin-2 | P53471 | 245.11 | - |
|  | Phosphoglycerate kinase | P41759 | 126.13 | 2.37E+05 |
|  | Putative 60s ribosomal protein L4 | G4VJ35 | 112.55 | - |
| 4 | Triosephosphate isomerase | G4V6B9 | 137.96 | 8.23E+06 |
|  | B-cell receptor-associated protein-like protein | A0A5K4EXK8 | 148.33 | 8.36E+05 |
| 5 | Glutathione S-transferase | P09792 | 243.00 | 1.15E+08 |
|  | 14-3-3 protein homolog 1 | Q26540 | 160.46 | 1.63E+06 |
| 6 | Putative 60s ribosomal protein L21 | G4VEA9 | 109.98 | 1.50E+05 |
|  | Transgelin | G4M0V2 | 78.55 | 7.14E+04 |

<sup>a</sup>SDS-GEL band as indicated in Figure 3C. <sup>b</sup>UniProt accession code. <sup>c</sup>PEAKS<sup>45</sup> peptide scores (-10logP) indicate high confidence in proteins present (*i.e.*, >50, $p < 0.00001$ ). <sup>d</sup>Integrated chromatographic peak area representing the MS1 signal intensity of the identified protein.

**Table 4:**
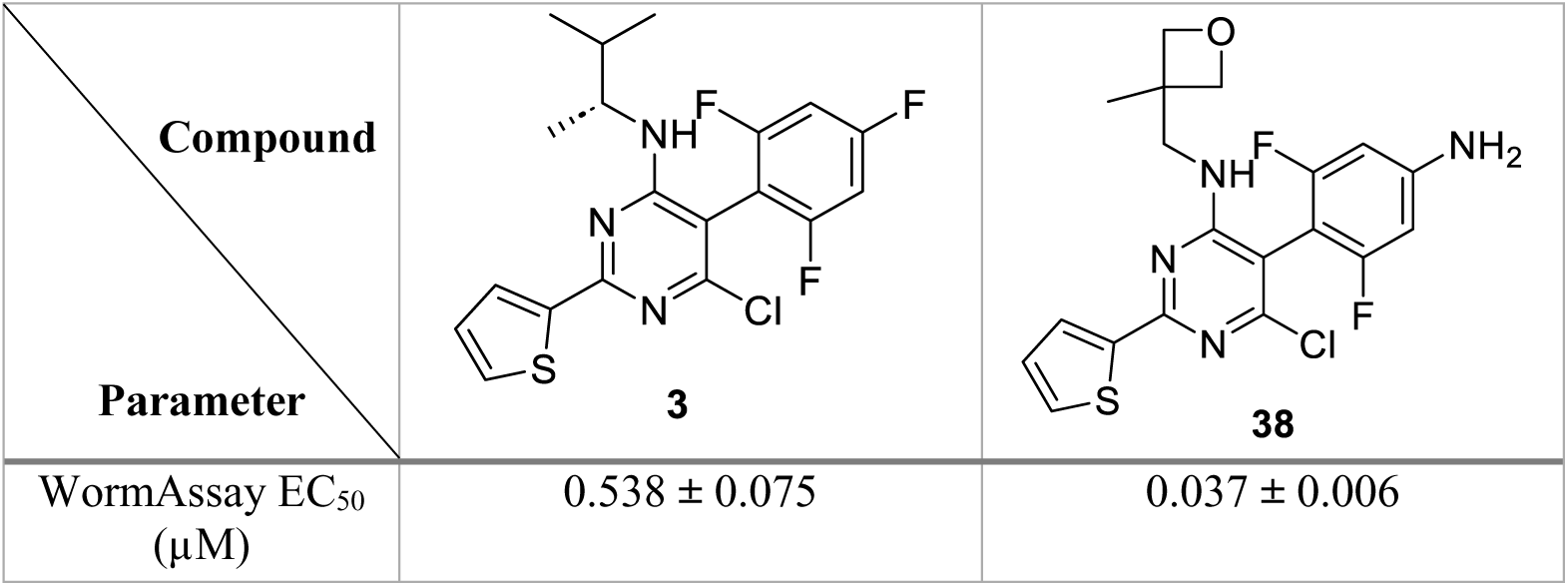
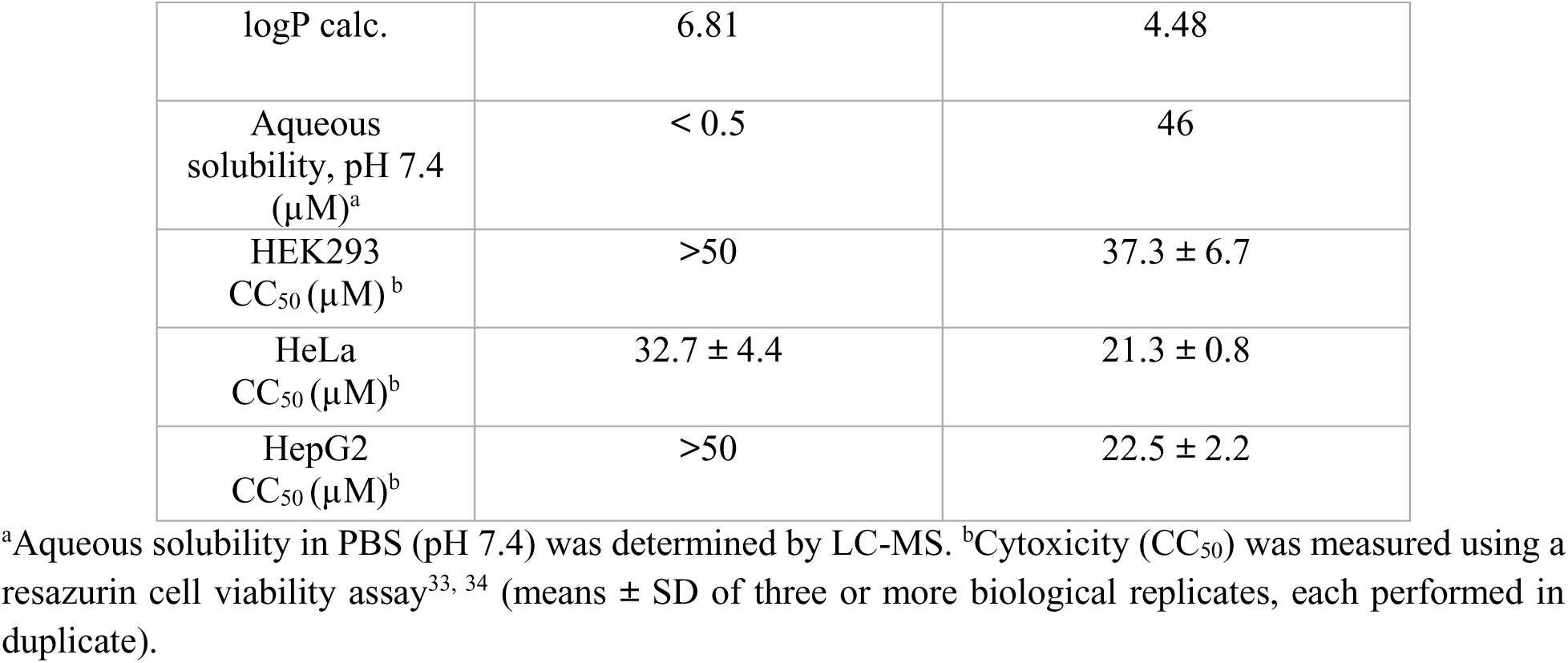
Comparison of the bioactivity and physicochemical properties of the progenitor TPP, 3, and the current compound, 38.

| Compound | **3** | **38** |
| --- | --- | --- |
| Parameter |  |  |
| WormAssay EC<sub>50</sub><br>(μM) | 0.538 ± 0.075 | 0.037 ± 0.006 |
| logP calc. | 6.81 | 4.48 |
| Aqueous solubility, pH 7.4 ($\mu\text{M}$)<sup>a</sup> | < 0.5 | 46 |
| HEK293 CC<sub>50</sub> ($\mu\text{M}$)<sup>b</sup> | >50 | 37.3 ± 6.7 |
| HeLa CC<sub>50</sub> ($\mu\text{M}$)<sup>b</sup> | 32.7 ± 4.4 | 21.3 ± 0.8 |
| HepG2 CC<sub>50</sub> ($\mu\text{M}$)<sup>b</sup> | >50 | 22.5 ± 2.2 |

<sup>a</sup>Aqueous solubility in PBS (pH 7.4) was determined by LC-MS. <sup>b</sup>Cytotoxicity (CC<sub>50</sub>) was measured using a resazurin cell viability assay<sup>33, 34</sup> (means ± SD of three or more biological replicates, each performed in duplicate).

## 3. Discussion

Our initial TPP hit compound, **3**, induced a sustained paralysis of *ex vivo S. mansoni* worms at sub-micromolar concentrations^13^ along with low cytotoxicity in mammalian cells (Table 3). However, its suboptimal physicochemical properties, specifically, low aqueous solubility (<0.5 µM) and high lipophilicity (cLogP = 6.81), made its preclinical development potentially problematic, particularly relating to oral absorption and bioavailability.^20, 21^ To this end, the SAR studies presented here, which examined a series of structural modifications for the fragments linked at C5 and C6 of the pyrimidine core of the TPP, identified selected substituents such as an oxetane-containing amine moiety at C5, and an *ortho, ortho-*difluoroaniline at C6, which mitigated the excessive lipophilic character of **3** and, as represented by **38**, modestly improved the aqueous solubility (<0.5 *vs* 46 µM) and achieved nanomolar paralytic potency (Table 4). Further, **38** demonstrated relatively low cytotoxicity against three mammalian cell lines (> 20 µM, Table 4) as well as a three-fold improvement in its plasma t_1/2_ compared to **3** (Table 2).

Apart from the improved biological and physicochemical properties of **38**, the compound generated a sustained paralysis of *ex vivo* juvenile and adult *S. mansoni* after a short (2 h) pulse at 0.6 and 1.2 µM, *i.e*., assay conditions based on the PK parameters measured for **38** in mice. The activity against juveniles is particularly encouraging in terms of identifying a preclinical candidate with a broader efficacy spectrum than the current drug, PZQ, given the latter’s well-established lack of *in vivo* activity against this developmental stage.^8, 37, 38^ In the same vein, the paralysis measured for the two other medically important schistosome species, *S. haematobium* and *S. japonicum* is also consistent with the TCP for a preclinical candidate, with, perhaps, the indication that a shared target(s) is being engaged. It remains to be determined, however, whether the particular paralysis generated by the TPP series translates to *in vivo* efficacy. Specifically, whereas the *in vitro* paralysis caused by **38** was persistent compared to the rapid recovery of worms after a similar pulse with PZQ, only PZQ visibly damaged the worm’s outer tegument—a feature understood to promote immune-mediated clearance of the parasite *in vivo*.^46–49^

Our SAR studies enabled the design of the TPP-based PAL probe, **46**, which labeled several *S. mansoni* proteins by SDS-PAGE florescence analysis, although, notably, not tubulin. These results suggest that even though the antischistosomal TPPs originated from the structurally related, and MT-active, phenylpyrimidines,^13, 15, 24^ the compounds do not engage tubulin in *S. mansoni* and that the paralysis measured involves other targets. Due to their intensity of fluorescence and the diminished signal after competition with the photostable TPP, **46**, six labelled bands were selected for identification by mass spectrometry-based proteomics. Several of the identified proteins are involved in muscle function, and may be directly or indirectly linked to the TPP-induced paralysis recorded, including the myosin heavy chain and putative paramyosin,^50^ both of which contribute to muscle contraction and cell motility,^51, 52^ as well as a calcium-transporting ATPase, which transports Ca^2+^ ions against their electrochemical gradient across cell membranes after muscle contraction.^53^ Thus, further studies are needed to unambiguously identify the TPP target(s) responsible for paralysis of *S. mansoni*.

## 4. Conclusion

This study expands on the SAR/SPR of TPPs as potent paralytic agents of adult and juvenile *S. mansoni,* and schistosomes in general. Through systematic structural modifications at the C5 and C6 positions of the pyrimidine core, we identified new TPP analogs with improved anti-schistosomal activity, reduced cytotoxicity, and enhanced physicochemical and PK properties. Notably, our application of the PAL technology revealed that the TPP mechanism of action is unlikely to directly involve tubulin, implicating alternative protein targets, one or more of which were potentially identified. The findings reported here encourage the further evaluation of TPPs as novel anti-schistosomal therapeutics. Future work will attempt to decrease further toxicity and enhance PK properties that would better support efficacy testing in a small animal model of schistosome infection.

## 5. **Experimental** Schistosome handling

The National Medical Research Institute (NMRI) isolate of *S. mansoni* was maintained using *Biomphalaria glabrata* snails as intermediate hosts and female Swiss Webster mice as definitive hosts. Mice (4 – 6 weeks old) were infected by subcutaneous injections of 200 cercariae. At 28 or 42 days-post-infection, hamsters were euthanized, and juvenile or adult *S. mansoni* worms, respectively, were harvested by perfusion of the hepatic portal and mesenteric venous system with DMEM.^29, 54^ Use of mice was in accordance with a protocol approved by the Institutional Animal Care and Use Committee (IACUC) at the

University of California, San Diego. Live *S. haematobium* and *S. japonicum* adults were shipped overnight from the Schistosomiasis Resource Center at the Biomedical Research Institute (Rockville, MD) in DMEM supplemented with 1% heat-inactvated FBS, 100 U/mL penicillin and 100 µg/mL streptomycin.

### 5.2 WormAssay to measure worm motility

After perfusion (*S. mansoni*) or overnight shipping (*S. haematobium* and *S. japonicum*), worms were washed five times in Basch medium 169^55^ supplemented with 100 U/mL penicillin and 100 µg/mL streptomycin, including one wash that contained 200 µg/mL amphotericin B. Worms (five adult male and two female worms, or 10 – 12 juvenile worms) were manually added to 24 well-plates and allowed to acclimatize overnight at 37 °C and 5% CO_2_ in 1 mL of the same medium containing 2.5% heat-inactvated FBS. Test compound in DMSO was then added and the volume brought up to 2 mL (0.1% final DMSO concentration). Incubations were maintained at 37 °C and 5% CO_2_ for the time periods indicated.

Average worm motility per well was measured using WormAssay^31, 32^ and percent motility was determined relative to DMSO controls. EC_50_ values in first pass singleton assays were calculated from five concentration points (0.3 – 5 µM), whereas for compounds of special interest, EC_50_ values were determined from eight concentration points (0.005 – 5 µM) in three or more biological replicates, each in triplicate. EC_50_ values were generated with Prism GraphPad Version 8.0 (San Diego, CA) using a sigmoidal four parameter logistic curve.

### 5.2. Cytotoxicity assay

HEK293, HepG2 and HeLa cell lines were cultured in DMEM supplemented with 10% heat-inactivated FBS and penicillin-streptomycin (1%) at 37 °C and 5% CO_2_. Cytotoxicity against each cell line was determined using the redox indicator, resazurin, ^33, 34^ in a 96-well microplate format.

Test compounds were serially diluted in DMSO to generated eight concentration points (0.02 – 50 µM). Compound (1 µL in DMSO) was added to the assay plate followed by addition of 49 mL fresh medium. Cells were diluted to 4 x 10^5^ cells/mL, and 50 µL of this suspension was added for a total density of 2 x 10^5^ cells/well. Assay plates were incubated at 37 °C and 5% CO_2._ for 48 h, followed by addition of 20 µL 0.5 mM resazurin solution in PBS. Assay plates were incubated for a further 2 h at 37 °C in the dark. The fluorescence signal of each well was measured using a 2104 EnVision multilabel plate reader at ex/em 531/595 nm. Assays were performed in three or more biological replicates, each in technical duplicate. CC_50_ values were calculated using Prism GraphPad Version 8.0 and a sigmoidal four parameter logistic curve.

### 5.3. Photoaffinity labelling

*Incubation and photoactivation*. PAL experiments were performed as previously described.^44^ Experiments were conducted under sterile conditions in a biosafety cabinet. In a 24-well microplate, five male and two female worms were incubated with 2 µM probe (0.1% final DMSO) in 1 mL Basch medium supplemented with 100 U/mL penicillin, 100 µg/mL streptomycin and 2.5% heat-inactivated FBS. Incubations were carried out at 37 °C and 5% CO₂ for 2 h. For competition experiments, worms were preincubated for 30 min with 50 µM of the photostable competitor (compound **38**) prior to the addition of 2 µM PAL probe. The microplate lids were then removed, and the worms were exposed for 30 min to a hand-held 365 nm UV light source positioned 1–2 cm above the surface of the medium. After photoactivation, the parasites were washed three times with PBS before freezing at −80 °C.

*Homogenization and click reaction*. Worms were lysed in 200 µL PBS containing 1X protease inhibitor cocktail (without EDTA; Thermo Cat#1862209) using a Fisherbrand Model 120 Sonic Dismembrator probe sonicator. Sonication was performed with 2-sec pulses and 5-sec pauses for a total duration of 10 min. The protein concentration was determined using the Pierce™ BCA Protein Assay Kit (ThermoFisher Cat#23227), and samples were normalized by dilution in PBS to 0.3 mg/mL. The click reaction was performed as previously described,^26, 44^ using 200 µL of the protein sample, 5 µL of a 5 mM Biotin-TAMRA azide solution (Click Chemistry Tools Cat#1048), and 25 µL of a click catalyst mixture prepared in a 3:1:1 ratio of 1.7 mM Tris(benzyltriazolylmethyl)amine (TBTA; in 80% t-butanol and 20% DMSO), 50 mM CuSO₄ (in water), and 50 mM Tris(2-carboxyethyl)phosphine (TCEP; adjusted to pH 7.0 in water). The reaction mixture was incubated at 37 °C for 45 min.

*SDS-PAGE and fluorescence visualization*. Clicked protein samples (30 µL) were mixed with 10 µL of 4X LDS sample loading buffer (Fisher Cat# NP0007) supplemented with 10% 2-mercaptoethanol (TCI Cat# M1948). Samples were denatured at 95 °C for 2 min and then loaded onto a precast Bolt™ 4–12% Bis-Tris Plus mini gel (1.00 mm). Electrophoresis was carried out at a constant voltage of 120 V for 1.5 h. Protein bands were visualized by fluorescence imaging using a Bio-Rad ChemiDoc MP imaging system, followed by staining with Coomassie Brilliant Blue G-250. Fluorescence signal intensities for the probe-competition experiments were densitometrically quantified using a Bio-Rad Image Lab 6.1 software. *Affinity purification*. Following the click reaction, proteins were precipitated by adding three volumes of cold acetone and incubating the mixture at −20 °C for 30 min. The precipitated proteins were collected by centrifugation at 2,000 × *g*, and the supernatant was carefully aspirated. The resulting pellet was washed with cold acetone, resuspended, and centrifuged twice more. Proteins were then subjected to affinity purification as previously described, using 50% NeutrAvidin agarose beads (25 µL; Fisher Cat#29200).^43^ The purified proteins were resolved by SDS–PAGE, visualized by fluorescence imaging, and stained with Coomassie Brilliant Blue G-250 as described above. Gel bands of interest were subsequently excised for proteomic analysis.

*In-gel digest and proteomics analysis.* In-gel digest of gel excisions was performed as previously reported.^43^ After digestion and peptide extraction, samples were desalted using C18 LTS tips (Rainin) and dried by vacuum centrifugation. Peptides were resuspended in 12 μL of 0.1% TFA and 5 μL of each sample were analyzed on a Q Exactive Mass Spectrometer (Thermo Scientific). Separation was performed by reverse-phase chromatography on an UltiMate 3000 HPLC system (Thermo Scientific) with an in-house packed C18 column (1.7 μm ACQUITY UPLC BEH, 75 μm × 25 cm) maintained at 65 °C. The flow rate was 300 nL/min over a 76-minute linear gradient from 5% Solvent A to 25% Solvent B, where Solvent A consisted of 0.1% formic acid in water and Solvent B comprised 0.1% formic acid in acetonitrile. Full MS survey scans were acquired at a resolution of 35,000 at 200 m/z across 350–1500 m/z, with automatic gain control (AGC) set to 1 × 10⁶ and a maximum injection time (maxIT) of 300 ms. Data-dependent MS/MS acquisition was performed using higher-energy collision dissociation (HCD) at 28 normalized collision energy on the 20 most intense precursor ions. MS/MS scans were collected at a resolution of 17,500 at 200 m/z with an AGC of 5 × 10⁶ and a maxIT of 50 ms. The PEAKS 8.5 software (Bioinformatics Solutions Inc.) was used to analyse the data.

### 5.4. Chemistry

*Materials and Methods*: All solvents and reagents were reagent grade. All reagents were purchased from reputable vendors and used as received. Thin layer chromatography (TLC) was performed with 200 μM MilliporeSigma precoated silica gel aluminum sheets. TLC spots were visualized under UV light or using KMnO_4_ stain. Flash chromatography was performed with SiliaFlash P60 (particle size 40-63 μM) supplied by Silicycle. Melting points were taken from Mel-Temp II by Barnstead Thermolyne, using an Omega digital thermometer. Infrared (IR) spectra were recorded on a PerkinElmer FT-IR spectrometer. Proton and carbon NMR spectra were recorded on a 600 MHz Bruker AVANCE III spectrometer. Chemical shifts were reported relative to residual solvent’s peak. High-resolution mass spectra were measured at the University of California San Diego Molecular Mass Spectrometry Facility. Analytical reverse phase high-performance liquid chromatography (HPLC) was performed using a SunFire C18 (4.6 x 50 mm, 5 mL) analytical column, while preparative reverse phase HPLC purifications were performed using a SunFire preparative C18 OBD column (5 μm 19 × 50 mm) on a Gilson HPLC equipped with a UV and mass detector. Samples were analyzed with analytical HPLC and employed 10% to 90% of CH_3_CN in H_2_O over 6–12 min and flow rate of 2 mL/min. Samples were purified by preparative HPLC employed 10% to 90% of CH_3_CN in H_2_O over 6–20 min and flow rate of 20 mL/min. All final compounds were >95% pure by LCMS (Agilent 1260 Infinity II LC with Infinity Lab LC/MSD).

**General procedure A.** To a solution of NaH (3.00 eq) in degassed anh. 1,4-dioxane (0.5-1 M), diethyl malonate (3.00 eq) was added dropwise at 60 °C. The suspension was stirred for 10 min at this temperature, followed by addition of CuBr (1.20 eq) and substituted bromobenzene (1.00 eq). The reaction mixture was stirred for 16 h at 100 °C. The resulting mixture was cooled to rt and 37% HCl (4.75 eq) was slowly added. The reaction mixture was then filtered through Celite^®^ to filter off excess CuBr and the resulting filtrate was extracted with EtOAc (3x) and washed with brine. The combined organic extracts were collected, dried over Na_2_SO_4_, and concentrated under reduced pressure. Purification via silica gel column chromatography (gradient of up to 10% EtOAc in hexanes) furnished a mixture of desired product and diethyl malonate that could be further purified via distillation under reduced pressure to obtain the desired product.

**General procedure B**. To a solution of NaH (1.30 eq) in anh. THF (0.5 M), diethyl malonate (1.10 eq) was added dropwise at 0 °C under N_2_. The suspension was stirred at rt for 1 h, followed by dropwise addition of substituted fluorobenzene (1.00 eq) at 0 °C. The resulting mixture was stirred at rt for 2 h. The reaction was quenched with satd. NH_4_Cl, extracted with EtOAc (3x), and washed with brine. The combined organic extracts were collected, dried over Na_2_SO_4_, and concentrated under reduced pressure. Purification via silica gel column chromatography (gradient of up to 10% EtOAc in hexanes) furnished a mixture of desired product and diethyl malonate that could be further purified via distillation under reduced pressure to obtain the desired product.

**General procedure C**. To a solution of substituted malonate (1.20 eq) in anh. NMP (0.6 M), thiophene-2-carboximidamide hydrochloride (1.20 eq) was added followed by DBU (2.00 eq) at rt under N_2_ atmosphere. The reaction was stirred at 95 °C for 5 h. The resulting mixture was cooled to 40 °C and 37% HCl (1.42 eq) was added at this temperature. The mixture was cooled to rt, and water was added to precipitate the desired product, which was vacuum filtered, washed with water, dried under vacuum for 2 h, and used directly in subsequent steps without further purification.

**General procedure D.** A solution of 5-substituted-2-(thiophen-2-yl)pyrimidine-4,6-diol (1.00 eq) in neat POCl_3_ (15.0 eq) was stirred in a pressure flask at 100 °C for 16 h. The resulting mixture was then cooled to rt, quenched with ice-water, and stirred for 30 min. Saturated aq. NaHCO_3_ was slowly added until > pH 5. The mixture was extracted with EtOAc (3x), washed with water (3x) and with brine. The combined organic extracts were collected, dried over Na_2_SO_4_, and concentrated under reduced pressure. Purification via silica gel column chromatography (gradient of up to 20% EtOAc in hexanes) furnished the desired product.

**General procedure E.** To a solution of 5-substituted-2-(thiophen-2-yl)pyrimidine-4,6-dichloride (1.00 eq) in anh. DMF (0.20–0.10 M), appropriate amine (1.00–2.20 eq) was added at rt. For hydrochloride amines, equimolar Et_3_N was added. The reaction mixture was stirred at rt for 16 h and then quenched with water, extracted with EtOAc (3x), and washed with water (5x). The combined organic extracts were collected, dried over Na_2_SO_4_, and concentrated under reduced pressure. Purification via silica gel column chromatography (gradient of up to 20% EtOAc in hexanes) furnished the desired product.

**General procedure F.** To a solution of *N-*substituted 6-chloro-5-(2,6-difluoro-4-methoxyphenyl)-2-(thiophen-2-yl)pyrimidin-4-amine (1.00 eq) in anh. CH_2_Cl_2_ (0.1 M), BBr_3_ was added at −78 °C (5.00 eq). The mixture was slowly warmed to rt and stirred for 16 h. Water was then added, followed by extraction with EtOAc (3x), and the combined organic extracts were collected, dried over Na_2_SO_4_, and concentrated under reduced pressure. Purification via silica gel column chromatography (gradient of up to 20% EtOAc in hexanes) furnished the desired product.

**General procedure G.** To a solution of *N-*substituted 6-chloro-5-(2,6-difluoro-4-nitrophenyl)-2-(thiophen-2-yl)pyrimidin-4-amine (1.00 eq) in water/MeOH (4:5, 0.1 M), iron (4.00 eq) and NH_4_Cl (5.00 eq) were added at rt. The reaction mixture was stirred at 80 °C for 2 h. The mixture was cooled to rt, filtered through a pad of Celite^®^, and washed with water. The filtrate was extracted with EtOAc (3x) and washed with brine. The combined organic extracts were dried over Na_2_SO_4_ and concentrated under reduced pressure. Purification via reverse phase HPLC furnished the desired product.

**(*R*)-6-Chloro-5-(3,5-difluorophenyl)-*N*-(3-methylbutan-2-yl)-2-(thiophen-2-yl)pyrimidin-4-amine (4, BL-0932)**. Synthesized according to General procedure E, using 4,6-dichloro-5-(3,5-difluorophenyl)-2-(thiophen-2-yl)pyrimidine (0.030 g, 0.087 mmol, 1.00 eq) and (R)-3-methylbutan-2-amine (0.017 g, 0.19 mmol, 2.20 eq). Purification via silica gel column chromatography furnished the desired product (0.022 g, 0.056 mmol, 64%). ^1^H NMR (600 MHz, CDCl_3_) δ 7.97 (d, *J* = 3.5 Hz, 1H), 7.46 (d, *J* = 5.0 Hz, 1H), 7.12 (t, *J* = 4.3 Hz, 1H), 6.89 (d, *J* = 30.7 Hz, 3H), 4.43 (d, *J* = 8.8 Hz, 1H), 4.22 (q, *J* = 6.8 Hz, 1H), 1.82 (h, *J* = 6.8 Hz, 1H), 1.11 (d, *J* = 6.8 Hz, 3H), 0.91 – 0.86 (m, 6H) ppm. ^13^C NMR (150 MHz, CDCl_3_) δ 164.62 (dd, *J* = 6.0, 3.7 Hz), 163.89 – 162.72 (m), 160.52, 160.14, 156.89, 142.99, 137.77 – 135.63 (m), 130.23, 129.52, 128.18, 113.37 (dd, *J* = 63.5, 22.6 Hz), 111.68, 104.76 (t, *J* = 25.0 Hz), 52.09, 32.80, 18.72, 18.40, 17.14 ppm. MS(ESI) m/z for C_19_H_18_ClF_2_N_3_S (393.1) observed [M+H]^+^: 394.1.

**6-Chloro-5-(2-fluorophenyl)-*N*-((*R*)-3-methylbutan-2-yl)-2-(thiophen-2-yl)pyrimidin-4-amine (5, BL-0933)**. Synthesized according to General procedure E, using 4,6-dichloro-5-(2-fluorophenyl)-2-(thiophen-2-yl)pyrimidine (0.030 g, 0.092 mmol, 1.00 eq) and (R)-3-methylbutan-2-amine (0.018 g, 0.20 mmol, 2.20 eq). Purification via silica gel column chromatography furnished the desired product (0.023 g, 0.061 mmol, 66%). ^1^H NMR (600 MHz, CDCl_3_) δ 7.98 (d, *J* = 3.7 Hz, 1H), 7.51 – 7.43 (m, 2H), 7.34 – 7.29 (m, 2H), 7.24 (d, *J* = 9.3 Hz, 1H), 7.14 – 7.09 (m, 1H), 4.38 (t, *J* = 7.1 Hz, 1H), 4.23 (ddd, *J* = 15.1, 9.1, 5.8 Hz, 1H), 1.80 (dtd, *J* = 13.8, 6.8, 2.6 Hz, 1H), 1.09 (t, *J* = 6.1 Hz, 3H), 0.86 (dd, *J* = 6.9, 2.5 Hz, 6H) ppm. ^13^C NMR (150 MHz, CDCl_3_) δ 161.33 – 160.81 (m), 160.16, 159.37, 159.25, 157.87, 157.81, 143.19, 132.32, 131.83, 131.37 (d, *J* = 8.0 Hz), 129.93, 129.27, 125.23 (d, *J* = 9.2 Hz), 120.31 (dd, *J* = 16.3, 10.3 Hz), 116.71 (dd, *J* = 26.3, 21.5 Hz), 107.83 (d, *J* = 1.9 Hz), 51.92, 51.83, 32.81, 32.78, 18.59, 18.41, 18.40, 18.24, 17.17, 17.08, 128.88 – 127.74 (m) ppm. MS(ESI) m/z for C_19_H_19_ClFN_3_S (375.1) observed [M+H]^+^: 376.1.

**(*R*)-6-Chloro-5-(4-fluorophenyl)-*N*-(3-methylbutan-2-yl)-2-(thiophen-2-yl)pyrimidin-4-amine (6, BL-0935)**. Synthesized according to General procedure E, using 4,6-dichloro-5-(4-fluorophenyl)-2-(thiophen-2-yl)pyrimidine (0.027 g, 0.083 mmol, 1.00 eq) and (R)-3-methylbutan-2-amine (0.016 g, 0.18 mmol, 2.20 eq). Purification via silica gel column chromatography furnished the desired product (0.017 g, 0.045 mmol, 54%). ^1^H NMR (600 MHz, CDCl_3_) δ 7.97 (d, *J* = 3.7 Hz, 1H), 7.44 (d, *J* = 5.0 Hz, 1H), 7.30 (dq, *J* = 10.1, 4.1 Hz, 2H), 7.22 (t, *J* = 8.6 Hz, 2H), 7.12 (t, *J* = 4.4 Hz, 1H), 4.43 (d, *J* = 8.3 Hz, 1H), 4.20 (h, *J* = 6.8 Hz, 1H), 1.80 (dq, *J* = 12.9, 6.8 Hz, 1H), 1.09 (d, *J* = 6.8 Hz, 3H), 0.86 (dd, *J* = 6.9, 4.5 Hz, 6H) ppm. ^13^C NMR (150 MHz, CDCl_3_) δ 163.77, 162.12, 161.12, 159.74, 157.08, 143.25, 132.05 (dd, *J* = 63.9, 8.0 Hz), 129.88, 129.18, 128.85 (d, *J* = 3.5 Hz), 128.10, 116.89 (dd, *J* = 21.6, 14.2 Hz), 112.87, 51.90, 32.77, 18.62, 18.38, 17.11 ppm. MS(ESI) m/z for C_19_H_19_ClFN_3_S (375.1) observed [M+H]^+^: 376.1.

**(*R*)-6-Chloro-5-(3,5-difluorophenyl)-*N*-(3-methylbutan-2-yl)-2-(thiophen-2-yl)pyrimidin-4-amine (7, BL-0936).** Synthesized according to General procedure E, using 4,6-dichloro-5-(3,5-difluorophenyl)-2-(thiophen-2-yl)pyrimidine (0.030 g, 0.087 mmol, 1.00 eq) and (R)-3-methylbutan-2-amine (0.017 g, 0.19 mmol, 2.20 eq). Purification via silica gel column chromatography furnished the desired product (0.022 g, 0.056 mmol, 64%). ^1^H NMR (600 MHz, CDCl_3_) δ 7.97 (d, *J* = 3.5 Hz, 1H), 7.46 (d, *J* = 5.0 Hz, 1H), 7.12 (t, *J* = 4.3 Hz, 1H), 6.89 (d, *J* = 30.7 Hz, 3H), 4.43 (d, *J* = 8.8 Hz, 1H), 4.22 (q, *J* = 6.8 Hz, 1H), 1.82 (h, *J* = 6.8 Hz, 1H), 1.11 (d, *J* = 6.8 Hz, 3H), 0.91 – 0.86 (m, 6H) ppm. ^13^C NMR (150 MHz, CDCl_3_) δ 164.62 (dd, *J* = 6.0, 3.7 Hz), 163.89 – 162.72 (m), 160.52, 160.14, 156.89, 142.99, 137.77 – 135.63 (m), 130.23, 129.52, 128.18, 113.37 (dd, *J* = 63.5, 22.6 Hz), 111.68, 104.76 (t, *J* = 25.0 Hz), 52.09, 32.80, 18.72, 18.40, 17.14 ppm. MS(ESI) m/z for C_19_H_18_ClF_2_N_3_S (393.1) observed [M+H]^+^: 394.1.

**6-Chloro-5-(2,5-difluorophenyl)-*N*-((*R*)-3-methylbutan-2-yl)-2-(thiophen-2-yl)pyrimidin-4-amine (8, BL-0937)**. Synthesized according to General procedure E, using 4,6-dichloro-5-(2,5-difluorophenyl)-2-(thiophen-2-yl)pyrimidine (0.025 g, 0.073 mmol, 1.00 eq) and (R)-3-methylbutan-2-amine (0.014 g, 0.16 mmol, 2.20 eq). Purification via silica gel column chromatography furnished the desired product (0.013 g, 0.033 mmol, 45%). ^1^H NMR (600 MHz, CDCl_3_) δ 7.99 (d, *J* = 3.9 Hz, 1H), 7.46 (d, *J* = 5.0 Hz, 1H), 7.24 – 7.14 (m, 2H), 7.12 (t, *J* = 4.4 Hz, 1H), 7.03 (dt, *J* = 9.2, 4.5 Hz, 1H), 4.40 – 4.35 (m, 1H), 4.25 (dh, *J* = 13.4, 6.6 Hz, 1H), 1.82 (dq, *J* = 13.2, 6.7 Hz, 1H), 1.11 (dd, *J* = 6.8, 3.3 Hz, 3H), 0.88 (t, *J* = 7.6 Hz, 6H) ppm. ^13^C NMR (150 MHz, CDCl_3_) δ 160.68 (d, *J* = 10.0 Hz), 160.47, 159.81 (dd, *J* = 4.3, 2.5 Hz), 158.67 – 158.11 (m), 157.92 (d, *J* = 9.4 Hz), 157.04 (dd, *J* = 17.0, 2.5 Hz), 155.42 (dd, *J* = 17.1, 2.4 Hz), 143.01 (d, *J* = 1.9 Hz), 132.32 (d, *J* = 111.4 Hz), 130.45 – 130.13 (m), 129.54 (d, *J* = 1.8 Hz), 128.15, 121.75 (ddd, *J* = 19.2, 8.5, 6.2 Hz), 118.56 (ddd, *J* = 73.7, 23.8, 2.8 Hz), 118.21 – 117.34 (m), 106.78, 52.10, 51.96, 32.86, 32.82, 18.71, 18.47, 18.41, 18.27, 17.25, 17.09 ppm. MS(ESI) m/z for C_19_H_18_ClF_2_N_3_S (393.1) observed [M+H]^+^: 394.1.

**(*R*)-4-(4-chloro-6-((3-methylbutan-2-yl)amino)-2-(thiophen-2-yl)pyrimidin-5-yl)-3,5-difluorophenol (9, BL-0929)**. To a solution of (R)-6-chloro-5-(2,6-difluoro-4-methoxyphenyl)-N-(3-methylbutan-2-yl)-2-(thiophen-2-yl)pyrimidin-4-amine (0.030 g, 0.071 mmol, 1.00 eq) in anh. CH_2_Cl_2_ (0.40 mL), BBr_3_ was added at −78 °C (0.089 g, 0.35 mmol, 5.00 eq). The mixture was slowly warmed to rt and stirred overnight. After 16 h, water was added, followed by extraction with EtOAc (3x). The combined organic extracts were collected, dried over Na_2_SO_4_, and concentrated in vacuo. Purification via silica gel column chromatography (gradient of up to 50% EtOAc in hexanes) furnished the desired product (0.026 g, 0.063 mmol, 90%). ^1^H NMR (600 MHz, CDCl_3_) δ 7.98 (d, *J* = 3.9 Hz, 1H), 7.45 (dd, *J* = 5.0, 1.3 Hz, 1H), 7.13 – 7.09 (m, 1H), 6.58 (d, *J* = 9.0 Hz, 2H), 4.46 (d, *J* = 8.3 Hz, 1H), 4.25 (q, *J* = 6.5 Hz, 1H), 1.12 (d, *J* = 6.8 Hz, 3H), 0.88 (dd, *J* = 6.9, 3.2 Hz, 6H) ppm. ^13^C NMR (150 MHz, DMSO) δ 161.80, 161.33, 161.07, 160.23 (d, *J* = 90.7 Hz), 159.34, 158.42, 142.35, 131.09, 129.20, 128.52, 101.88, 99.93 (dd, *J* = 23.8, 8.1 Hz), 52.14, 32.40, 19.35, 18.93, 16.99 ppm. MS(ESI) m/z for C_19_H_18_ClF_2_N_3_OS (409.1) observed [M+H]+: 410.1.

**(*R*)-6-Chloro-5-(2,6-difluoro-4-methoxyphenyl)-*N*-(3-methylbutan-2-yl)-2-(thiophen-2-yl)pyrimidin-4-amine (10, BL-0934)**. Synthesized according to General procedure E, using 4,6-dichloro-5-(2,6-difluoro-4-methoxyphenyl)-2-(thiophen-2-yl)pyrimidine (0.100 g, 0.27 mmol, 1.00 eq) and (R)-3-methylbutan-2-amine (0.051 g, 0.59 mmol, 2.20 eq). Purification via silica gel column chromatography furnished the desired product (0.088 g, 0.21 mmol, 77%). ^1^H NMR (600 MHz, CDCl_3_) δ 7.99 (d, *J* = 3.7 Hz, 1H), 7.45 (d, *J* = 5.0 Hz, 1H), 7.12 (t, *J* = 4.4 Hz, 1H), 6.62 (d, *J* = 9.2 Hz, 2H), 4.41 (d, *J* = 8.6 Hz, 1H), 4.26 (h, *J* = 6.6 Hz, 1H), 3.86 (s, 3H), 1.83 (h, *J* = 6.6 Hz, 1H), 1.12 (d, *J* = 6.8 Hz, 3H), 0.88 (dd, *J* = 7.1, 4.1 Hz, 6H) ppm. ^13^C NMR (150 MHz, CDCl_3_) δ 162.34 (t, *J* = 13.8 Hz), 162.11 (dd, *J* = 25.5, 9.9 Hz), 161.26, 160.58, 160.58 – 160.28 (m), 159.25, 143.16, 130.07, 129.46, 128.11, 101.88, 101.56 (t, *J* = 21.4 Hz), 98.86 (ddd, *J* = 25.6, 17.0, 3.3 Hz), 56.08, 51.92, 32.91, 18.54, 18.32, 17.23 ppm. MS(ESI) m/z for C_20_H_20_ClF_2_N_3_OS (423.1) observed [M+H]+: 424.1.

(*R*)-6-Chloro-5-(2,6-difluoro-4-(3-(methylamino)propoxy)phenyl)-*N*-(3-methylbutan-2-yl)-2-(thiophen-2-yl)pyrimidin-4-amine (11, BL-0947). To a solution of *tert*-butyl (R)-(3-(4-(4-chloro-6-((3-methylbutan-2-yl)amino)-2-(thiophen-2-yl)pyrimidin-5-yl)-3,5-difluorophenoxy)propyl)(methyl)carbamate (0.033 g, 0.057 mmol, 1.00 eq) in anh. dioxane, HCl (4.0 M in dioxane, 0.071 mL, 5.00 eq) was added at rt. The reaction mixture was stirred at rt for 16 h. The resulting mixture was concentrated under reduced pressure, basified with aq. 1M NaOH and extracted with EtOAc (3x). The combined organic extracts were dried over Na_2_SO_4_ and concentrated under reduced pressure. The product was purified via reverse phase HPLC to yield the desired product as a formate salt (0.026 g, 0.049 mmol, 87%). ^1^H NMR (600 MHz, DMSO) δ 8.31 (s, 1H), 7.82 (d, *J* = 3.9 Hz, 1H), 7.72 (d, *J* = 5.0 Hz, 1H), 7.15 (t, *J* = 4.4 Hz, 1H), 6.86 (d, *J* = 10.1 Hz, 2H), 6.78 (d, *J* = 8.6 Hz, 1H), 4.11 (t, *J* = 7.1 Hz, 3H), 2.89 (t, *J* = 7.5 Hz, 2H), 2.46 (s, 3H), 2.01 (p, *J* = 6.7 Hz, 2H), 1.77 (hept, *J* = 6.8 Hz, 1H), 1.05 (d, *J* = 6.8 Hz, 3H), 0.81 (dd, *J* = 17.0, 6.7 Hz, 6H) ppm. ^13^C NMR (150 MHz, DMSO) δ 162.69, 161.92 – 161.09 (m), 160.68, 160.12, 159.71 (dd, *J* = 15.8, 9.2 Hz), 158.77, 141.15, 132.33, 130.59, 128.73, 117.49, 105.42 (d, *J* = 18.3 Hz), 104.17 – 95.75 (m), 29.53 ppm. MS(ESI) m/z for C_23_H_27_ClF_2_N_4_OS (480.2) observed [M+H]+: 481.2.

**(*R*)-5-(4-Amino-2,6-difluorophenyl)-6-chloro-*N*-(3-methylbutan-2-yl)-2-(thiophen-2-yl)pyrimidin-4-amine (12, BL-0946)**. To a solution of (R)-6-chloro-5-(2,6-difluoro-4-nitrophenyl)-N-(3-methylbutan-2-yl)-2-(thiophen-2-yl)pyrimidin-4-amine (0.030 g, 0.068 mmol, 1.00 eq) in water/MeOH (4:5, 0.135 mL), iron (0.015 g, 0.27 mmol, 4.00 eq) and NH_4_Cl (0.018 g, 0.34 mmol, 5.00 eq) were added at rt. The reaction mixture was stirred at 80 °C for 2 h. The mixture was cooled to rt, filtered through a pad of Celite^®^, and washed with water. The filtrate was extracted with EtOAc (3x) and washed with brine. The combined organic extracts were dried over Na_2_SO_4_ and concentrated under reduced pressure. Purification via reverse phase HPLC furnished the desired product (0.015 g, 0.036 mmol, 54%). ^1^H NMR (600 MHz, CDCl_3_) δ 7.97 (d, *J* = 3.9 Hz, 1H), 7.44 (d, *J* = 5.0 Hz, 1H), 7.11 (t, *J* = 4.4 Hz, 1H), 6.32 (d, *J* = 9.4 Hz, 2H), 4.50 (d, *J* = 8.6 Hz, 1H), 4.25 (h, *J* = 6.6 Hz, 1H), 4.13 (s, 2H), 1.82 (h, *J* = 6.7 Hz, 1H), 1.11 (d, *J* = 6.6 Hz, 3H), 0.88 (dd, *J* = 7.0, 4.6 Hz, 6H) ppm. ^13^C NMR (150 MHz, CDCl_3_) δ 162.28 (dd, *J* = 24.2, 9.8 Hz), 161.47, 160.64 (dd, *J* = 23.6, 9.7 Hz), 160.31, 159.23, 149.90 (t, *J* = 13.7 Hz), 143.24, 129.93, 129.30, 128.09, 102.49, 99.07 – 97.60 (m), 51.87, 32.91, 18.53, 18.32, 17.22 ppm. MS(ESI) m/z for C_19_H_19_ClF_2_N_4_S (408.1) observed [M+H]+: 409.1.

**(*R*)-6-Chloro-5-(2,6-difluoro-4-nitrophenyl)-*N*-(3-methylbutan-2-yl)-2-(thiophen-2-yl)pyrimidin-4-amine (13, BL-0945)**. Synthesized according to General procedure E, using 4,6-dichloro-5-(2,6-difluoro-4-nitrophenyl)-2-(thiophen-2-yl)pyrimidine (0.100 g, 0.258 mmol, 1.00 eq) and (R)-3-methylbutan-2-amine (0.050 g, 0.567 mmol, 2.20 eq). Purification via silica gel column chromatography furnished the desired product (0.105 g, 0.239 mmol, 93%). ^1^H NMR (600 MHz, CDCl_3_) δ 8.02 (d, *J* = 3.7 Hz, 1H), 8.00 – 7.96 (m, 2H), 7.50 (d, *J* = 5.0 Hz, 1H), 7.14 (dd, *J* = 5.0, 3.7 Hz, 1H), 4.30 (dt, *J* = 8.3, 6.2 Hz, 1H), 4.21 (d, *J* = 8.3 Hz, 1H), 1.84 (dq, *J* = 13.4, 6.8 Hz, 1H), 1.13 (d, *J* = 6.8 Hz, 3H), 0.89 (dd, *J* = 6.8, 3.7 Hz, 6H) ppm. ^13^C NMR (150 MHz, CDCl_3_) δ 161.48, 161.44, 160.31, 159.70 (dd, *J* = 31.9, 6.7 Hz), 158.84, 149.49 (t, *J* = 10.8 Hz), 142.56, 130.85, 130.16, 128.29, 117.23 (t, *J* = 21.0 Hz), 108.54 (ddd, *J* = 26.4, 21.9, 3.9 Hz), 99.63, 52.34, 32.89, 18.68, 18.31, 17.20 ppm. MS(ESI) m/z for C_19_H_17_ClF_2_N_4_O_2_S (438.1) observed [M+H]+: 439.1.

**(*R*)-4-(4-Chloro-6-((3-methylbutan-2-yl)amino)-2-(thiophen-2-yl)pyrimidin-5-yl)-3,5-difluorobenzonitrile (14, BL-0995)**. Synthesized according to General procedure E, using 4-(4,6-dichloro-2-(thiophen-2-yl)pyrimidin-5-yl)-3,5-difluorobenzonitrile (0.030 g, 0.081 mmol, 1.00 eq) and (*R*)-3-methylbutan-2-amine (0.015 g, 0.17 mmol, 2.10 eq). Purification via silica gel column chromatography furnished the desired product (0.022 g, 0.052 mmol, 65%). ^1^H NMR (600 MHz, CDCl_3_) δ 8.01 (dd, *J* = 3.7, 1.3 Hz, 1H), 7.49 (dd, *J* = 5.0, 1.2 Hz, 1H), 7.43 – 7.38 (m, 2H), 7.13 (dd, *J* = 5.0, 3.7 Hz, 1H), 4.29 (dt, *J* = 13.3, 6.7 Hz, 1H), 4.20 (d, *J* = 8.4 Hz, 1H), 1.83 (dq, *J* = 13.3, 6.7 Hz, 1H), 1.13 (d, *J* = 6.7 Hz, 3H), 0.89 (dd, *J* = 6.8, 3.3 Hz, 6H) ppm. ^13^C NMR (150 MHz, CDCl_3_) δ 161.62 (d, *J* = 23.1 Hz), 161.42, 160.41, 160.14 – 159.77 (m), 158.91, 142.63, 130.81, 130.14, 128.30, 117.18 – 115.81 (m), 115.32 (d, *J* = 11.7 Hz), 99.74, 52.29, 32.91, 18.64, 18.33, 17.26 ppm. MS(ESI) m/z for C_20_H_17_ClF_2_N_4_S (411.8) observed [M+H]+: 412.1.

**6-Chloro-2-(thiophen-2-yl)-5-(2,4,6-trifluorophenyl)pyrimidin-4-amine (15, BL-0923)**. Synthesized according to General procedure E, using 4,6-dichloro-2-(thiophen-2-yl)-5-(2,4,6-trifluorophenyl) pyrimidine (0.100 g, 0.277 mmol, 1.00 eq) and ammonia (7.0 M in MeOH, 0.079 mL, 2.00 eq), using MeOH as solvent (1.8 mL) instead of DMF. Purification via silica gel column chromatography furnished the desired product (0.028 g, 0.082 mmol, 30%). ^1^H NMR (600 MHz, CDCl_3_) δ 8.01 (d, *J* = 3.7 Hz, 1H), 7.50 (d, *J* = 5.0 Hz, 1H), 7.13 (t, *J* = 4.4 Hz, 1H), 6.85 (dd, *J* = 8.6, 7.0 Hz, 2H), 4.95 (s, 2H) ppm. ^13^C NMR (150 MHz, CDCl_3_) δ 165.78 – 164.10 (m), 163.98 – 162.78 (m), 162.89, 162.46, 161.87 (dd, *J* = 12.4, 7.8 Hz), 161.24, 160.50, 160.11 (dd, *J* = 14.6, 9.3 Hz), 141.94, 130.73, 130.22, 128.35 (d, *J* = 1.7 Hz), 107.04 – 105.51 (m), 102.48 – 99.73 (m) ppm. MS(ESI) m/z for C_14_H_7_ClF_3_N_3_S (341.0) observed [M+H]+: 342.0.

***N*-(6-Chloro-2-(thiophen-2-yl)-5-(2,4,6-trifluorophenyl)pyrimidin-4-yl)isobutyramide (16, BL-0931)**. To a solution of 6-chloro-2-(thiophen-2-yl)-5-(2,4,6-trifluorophenyl)pyrimidin-4-amine (0.020 g, 0.059 mmol, 1.00 eq) in anh. CH_2_Cl_2_ (0.50 mL), DMAP (0.0014 g, 0.012 mmol, 0.20 eq), and Et_3_N (0.0095 g, 0.094 mmol, 1.60 eq) were added. The mixture was cooled to 0 C, and isobutyryl chloride (0.0094 g, 0.088 mmol, 1.50 eq) was added. The reaction mixture was warmed to rt and stirred overnight. The resulting mixture was diluted with water and extracted with EtOAc (3x). The combined organic extracts were collected, dried over Na_2_SO_4_, and concentrated in vacuo. Purification via silica gel column chromatography furnished the desired product (0.014 g, 0.034 mmol, 58%). ^1^H NMR (600 MHz, CDCl_3_) δ 8.05 (dd, *J* = 3.7, 1.3 Hz, 1H), 7.56 (dd, *J* = 5.0, 1.3 Hz, 1H), 7.41 (s, 1H), 7.17 (dd, *J* = 5.2, 3.6 Hz, 1H), 6.83 (dd, *J* = 8.8, 7.0 Hz, 2H), 3.32 (p, *J* = 6.8 Hz, 1H), 1.19 (d, *J* = 6.8 Hz, 6H) ppm. ^13^C NMR (150 MHz, CDCl_3_) δ 177.23, 165.30 – 164.25 (m), 163.20 (t, *J* = 16.0 Hz), 162.40, 161.56 – 161.27 (m), 161.16, 159.74 (dd, *J* = 14.9, 9.1 Hz), 157.63, 141.27, 131.85, 131.12, 128.74, 108.40 – 107.62 (m), 103.05 – 100.23 (m), 35.51, 19.19 ppm. MS(ESI) m/z for C_18_H_13_ClF_3_N_3_OS (411.0) observed [M+H]+: 412.0.

**6-Chloro-*N*-isobutyl-2-(thiophen-2-yl)-5-(2,4,6-trifluorophenyl)pyrimidin-4-amine (17, BL-0920)**. Synthesized according to General procedure E, using 4,6-dichloro-2-(thiophen-2-yl)-5-(2,4,6-trifluorophenyl) pyrimidine (0.030 g, 0.083 mmol, 1.00 eq) and 2-methylpropan-1-amine (0.013 g, 0.18 mmol, 2.20 eq). Purification via silica gel column chromatography furnished the desired product (0.031 g, 0.078 mmol, 94%). ^1^H NMR (600 MHz, CDCl_3_) δ 8.00 (dd, *J* = 3.7, 1.5 Hz, 1H), 7.47 (dd, *J* = 5.0, 1.5 Hz, 1H), 7.12 (dd, *J* = 5.3, 3.5 Hz, 1H), 6.88 – 6.80 (m, 2H), 4.72 (dq, *J* = 12.1, 6.1 Hz, 1H), 3.38 (t, *J* = 6.5 Hz, 2H), 1.96 (dpd, *J* = 13.4, 6.7, 2.3 Hz, 1H), 0.92 (d, *J* = 6.8 Hz, 6H) ppm. ^13^C NMR (150 MHz, CDCl_3_) δ 164.59 (t, *J* = 15.3 Hz), 162.92 (t, *J* = 15.2 Hz), 161.88 (dd, *J* = 14.9, 9.1 Hz), 161.46, 160.88, 160.21 (dd, *J* = 14.9, 9.2 Hz), 159.17, 142.84, 130.38, 129.72, 128.17, 106.13 (td, *J* = 21.2, 4.7 Hz), 101.83 – 100.97 (m), 100.88, 48.94, 28.44, 20.10 ppm. MS(ESI) m/z for C_18_H_15_ClF_3_N_3_S (397.1) observed [M+H]+: 398.1.

**6-Chloro-*N*-cyclopropyl-2-(thiophen-2-yl)-5-(2,4,6-trifluorophenyl)pyrimidin-4-amine (18, BL-0913)**. Synthesized according to General procedure E, using 4,6-dichloro-2-(thiophen-2-yl)-5-(2,4,6-trifluorophenyl) pyrimidine (0.030 g, 0.083 mmol, 1.00 eq) and cyclopropanamine (0.010 g, 0.17 mmol, 2.00 eq). Purification via silica gel column chromatography furnished the desired product (0.026 g, 0.068 mmol, 82%). ^1^H NMR (600 MHz, CDCl_3_) δ 8.04 (d, *J* = 3.9 Hz, 1H), 7.48 (d, *J* = 5.0 Hz, 1H), 7.13 (t, *J* = 4.4 Hz, 1H), 6.84 – 6.79 (m, 2H), 4.79 (s, 1H), 2.89 (dq, *J* = 7.0, 3.4 Hz, 1H), 0.85 (d, *J* = 5.7 Hz, 2H), 0.56 – 0.48 (m, 2H) ppm. ^13^C NMR (150 MHz, CDCl_3_) δ 164.60 (t, *J* = 15.6 Hz), 162.93 (t, *J* = 15.3 Hz), 162.52, 161.81 (dd, *J* = 14.8, 9.2 Hz), 161.05, 160.14 (dd, *J* = 15.2, 9.0 Hz), 159.32, 142.78, 130.57, 129.96, 128.22, 106.46 – 105.82 (m), 102.85 – 100.79 (m), 24.68, 7.58, 1.18 ppm. HRMS (ES^+^) calculated for [C_17_H_12_ClF_3_N_3_S]^+^ 382.0387, found 382.0388.

**6-Chloro-*N*-cyclobutyl-2-(thiophen-2-yl)-5-(2,4,6-trifluorophenyl)pyrimidin-4-amine (19, BL-0915)**. Synthesized according to General procedure E, using 4,6-dichloro-2-(thiophen-2-yl)-5-(2,4,6-trifluorophenyl) pyrimidine (0.030 g, 0.083 mmol, 1.00 eq) and cyclobutanamine (0.012 g, 0.17 mmol, 2.10 eq). Purification via silica gel column chromatography furnished the desired product (0.027 g, 0.068 mmol, 82%). ^1^H NMR (600 MHz, CDCl_3_) δ 8.00 (d, *J* = 3.7 Hz, 1H), 7.47 (d, *J* = 5.0 Hz, 1H), 7.12 (t, *J* = 4.4 Hz, 1H), 6.83 (dd, *J* = 8.7, 6.9 Hz, 2H), 4.69 (d, *J* = 6.6 Hz, 1H), 4.62 (h, *J* = 7.8 Hz, 1H), 2.46 (tdd, *J* = 9.4, 7.1, 2.9 Hz, 2H), 1.88 (pd, *J* = 9.8, 3.0 Hz, 2H), 1.79 (dt, *J* = 16.9, 10.1 Hz, 2H) ppm. ^13^C NMR (150 MHz, CDCl_3_) δ 164.61 (t, *J* = 15.2 Hz), 162.93 (t, *J* = 15.4 Hz), 161.85 (dd, *J* = 15.2, 9.4 Hz), 160.96, 160.40, 160.18 (dd, *J* = 15.2, 9.4 Hz), 159.30, 142.80, 130.43, 129.83, 128.19, 107.40 – 105.39 (m), 102.49 – 99.59 (m), 47.12, 31.22, 15.34 ppm. MS(ESI) m/z for C_18_H_13_ClF_3_N_3_S (395.1) observed [M+H]+: 396.1.

**6-Chloro-*N*-(cyclobutylmethyl)-2-(thiophen-2-yl)-5-(2,4,6-trifluorophenyl)pyrimidin-4-amine (20, BL-0916)**. Synthesized according to General procedure E, using 4,6-dichloro-2-(thiophen-2-yl)-5-(2,4,6-trifluorophenyl) pyrimidine (0.030 g, 0.083 mmol, 1.00 eq), cyclobutylmethanamine hydrochloride (0.012 g, 0.10 mmol, 1.20 eq), and Et_3_N (0.010 g, 0.10 mmol, 1.20 eq). Purification via silica gel column chromatography furnished the desired product (0.031 g, 0.076 mmol, 91%). ^1^H NMR (600 MHz, CDCl_3_) δ 8.01 (dd, *J* = 3.8, 1.4 Hz, 1H), 7.48 (dd, *J* = 5.0, 1.3 Hz, 1H), 7.13 (dd, *J* = 5.2, 3.6 Hz, 1H), 6.84 (dd, *J* = 8.8, 6.8 Hz, 2H), 4.53 (t, *J* = 5.8 Hz, 1H), 3.57 (dd, *J* = 7.3, 5.6 Hz, 2H), 2.56 (h, *J* = 7.7 Hz, 1H), 2.03 (ddt, *J* = 12.5, 10.1, 6.0 Hz, 2H), 1.93 – 1.83 (m, 2H), 1.76 – 1.66 (m, 2H) ppm. ^13^C NMR (150 MHz, CDCl_3_) δ 165.22 – 164.45 (m), 163.44 – 162.80 (m), 161.92 (dd, *J* = 14.8, 9.5 Hz), 161.51, 160.99, 160.55 – 159.93 (m), 159.23, 106.20 (t, *J* = 19.8 Hz), 103.64 – 100.02 (m), 46.68, 35.22, 25.81, 18.51 ppm. MS(ESI) m/z for C_19_H_15_ClF_3_N_3_S (409.1) observed [M+H]+: 410.1.

**6-Chloro-*N*-((1-methylcyclopropyl)methyl)-2-(thiophen-2-yl)-5-(2,4,6-trifluorophenyl)pyrimidin-4-amine (21, BL-0993)**. Synthesized according to General procedure E, using 4,6-dichloro-2-(thiophen-2-yl)-5-(2,4,6-trifluorophenyl)pyrimidine (0.050 g, 0.138 mmol, 1.00 eq), (1-Methylcyclopropyl)methanamine hydrochloride (0.020 g, 0.166 mmol, 1.20 eq), and Et_3_N (0.028 g, 0.277 mmol, 2.00 eq). Purification via silica gel column chromatography furnished the desired product (0.040 g, 0.098 mmol, 70%). ^1^H NMR (600 MHz, CDCl_3_) δ 8.05 (d, *J* = 3.9 Hz, 1H), 7.54 (d, *J* = 5.0 Hz, 1H), 7.15 (t, *J* = 4.4 Hz, 1H), 6.92 – 6.81 (m, 2H), 4.94 (s, 1H), 3.47 (s, 2H), 1.05 (s, 3H), 0.48 (s, 2H), 0.34 (s, 1H) ppm. ^13^C NMR (150 MHz, CDCl_3_) δ 164.95, 163.27, 162.08 – 161.72 (m), 161.50, 160.41 – 160.11 (m), 159.97, 157.37, 140.99, 131.60, 130.73, 128.64, 105.20, 102.06 – 101.01 (m), 50.19, 21.29, 16.26, 11.78 ppm. HRMS (ES^+^): calcd for C_19_H_16_ClF_3_N_3_S [M + H]^+^, 410.0700; found, 410.0696.

**6-Chloro-*N*-(oxetan-3-yl)-2-(thiophen-2-yl)-5-(2,4,6-trifluorophenyl)pyrimidin-4-amine (22, BL-0907)**. Synthesized according to General procedure E, using 4,6-dichloro-2-(thiophen-2-yl)-5-(2,4,6-trifluorophenyl) pyrimidine (0.030 g, 0.083 mmol, 1.00 eq) and oxetan-3-amine (0.012 g, 0.17 mmol, 2.00 eq). Purification via silica gel column chromatography furnished the desired product (0.012 g, 0.030 mmol, 36%). ^1^H NMR (600 MHz, CDCl_3_) δ 7.98 (dd, *J* = 3.7, 1.3 Hz, 1H), 7.49 (dd, *J* = 4.9, 1.2 Hz, 1H), 7.13 (dd, *J* = 5.0, 3.7 Hz, 1H), 6.88 – 6.83 (m, 2H), 5.25 (h, *J* = 6.1 Hz, 1H), 5.05 (d, *J* = 5.9 Hz, 1H), 5.01 (t, *J* = 7.1 Hz, 2H), 4.55 (t, *J* = 6.6 Hz, 2H) ppm. ^13^C NMR (151 MHz, CDCl_3_) δ 164.81 (t, *J* = 15.2 Hz), 163.13 (t, *J* = 15.0 Hz), 161.83 (dd, *J* = 15.1, 9.3 Hz), 160.92, 160.35, 160.16 (dd, *J* = 14.8, 9.0 Hz), 159.92, 142.20, 130.90, 130.18, 128.36, 105.70, 101.49, 78.37, 47.13 ppm. HRMS (ES^+^) calcd for [C_17_H_11_ClF_3_N_3_OS]^+^ 398.0336, found 398.0339.

**6-Chloro-*N*-((3-methyloxetan-3-yl)methyl)-2-(thiophen-2-yl)-5-(2,4,6-trifluorophenyl)pyrimidin-4-amine (23, BL-0912)**. Synthesized according to General procedure E, using 4,6-dichloro-2-(thiophen-2-yl)-5-(2,4,6-trifluorophenyl) pyrimidine (0.030 g, 0.083 mmol, 1.00 eq) and (3-methyloxetan-3-yl)methanamine (0.017 g, 0.17 mmol, 2.00 eq). Purification via silica gel column chromatography furnished the desired product (0.029 g, 0.068 mmol, 82%). ^1^H NMR (600 MHz, CDCl_3_) δ 8.01 (dd, *J* = 3.8, 1.2 Hz, 1H), 7.48 (dd, *J* = 5.0, 1.1 Hz, 1H), 7.13 (dd, *J* = 5.0, 3.7 Hz, 1H), 6.85 (dd, *J* = 8.8, 6.8 Hz, 2H), 4.97 (t, *J* = 6.1 Hz, 1H), 4.51 (d, *J* = 6.1 Hz, 2H), 4.38 (d, *J* = 5.9 Hz, 2H), 3.78 (d, *J* = 6.1 Hz, 2H), 1.30 (s, 3H) ppm. ^13^C NMR (150 MHz, CDCl_3_) δ 164.73 (t, *J* = 14.9 Hz), 163.27 – 162.85 (m), 161.86 (dd, *J* = 14.9, 9.1 Hz), 161.71, 160.94, 160.18 (dd, *J* = 14.9, 9.2 Hz), 159.54, 105.84 (q, *J* = 16.5 Hz), 101.42 (dd, *J* = 53.0, 27.5 Hz), 80.55, 47.79, 40.68, 21.91 ppm. HRMS (ES+) calculated for [C19H16ClF3N3OS]+ 426.0649, found 426.0651.

**4-Chloro-6-(pyrrolidin-1-yl)-2-(thiophen-2-yl)-5-(2,4,6-trifluorophenyl)pyrimidine (24, BL-0914).** Synthesized according to general procedure E, using 4,6-dichloro-2-(thiophen-2-yl)-5-(2,4,6-trifluorophenyl) pyrimidine (0.030 g, 0.083 mmol, 1.00 eq) and pyrrolidine (0.012 mg, 0.17 mmol, 2.10 eq). The product was purified via column chromatography to yield the desired product (0.018 g, 0.043 mmol, 52%). ^1^H NMR (600 MHz, CDCl_3_) δ 7.98 (d, *J* = 3.7 Hz, 1H), 7.45 (d, *J* = 5.0 Hz, 1H), 7.11 (t, *J* = 4.3 Hz, 1H), 6.75 (t, *J* = 7.7 Hz, 2H), 3.31 (s, 4H), 1.86 – 1.83 (m, 4H) ppm. ^13^C NMR (150 MHz, CDCl_3_) δ 164.14 (t, *J* = 15.6 Hz), 162.47 (t, *J* = 15.8 Hz), 162.05 (dd, *J* = 15.3, 9.3 Hz), 161.07, 160.39 (dd, *J* = 15.2, 9.3 Hz), 159.86, 159.83, 142.91, 110.42 (td, *J* = 21.6, 5.6 Hz), 100.72 – 99.99 (m), 99.76, 48.86, 25.43 ppm. MS(ESI) m/z for C_18_H_13_ClF_3_N_3_S (395.1) observed [M+H]+: 396.1.

**4-Chloro-6-(piperidin-1-yl)-2-(thiophen-2-yl)-5-(2,4,6-trifluorophenyl)pyrimidine (25, BL-0917)**. Synthesized according to General procedure E, using 4,6-dichloro-2-(thiophen-2-yl)-5-(2,4,6-trifluorophenyl) pyrimidine (0.030 g, 0.083 mmol, 1.00 eq) and piperidine (0.016 g, 0.18 mmol, 2.20 mmol). Purification via silica gel column chromatography furnished the desired product (0.018 g, 0.044 mmol, 53%). ^1^H NMR (600 MHz, CDCl_3_) δ 7.99 (d, *J* = 4.0 Hz, 1H), 7.46 (d, *J* = 5.0 Hz, 1H), 7.12 (t, *J* = 4.4 Hz, 1H), 6.79 (t, *J* = 7.8 Hz, 2H), 3.40 – 3.37 (m, 4H), 1.59 (p, *J* = 5.9 Hz, 2H), 1.47 (p, *J* = 5.7 Hz, 4H) ppm. ^13^C NMR (150 MHz, CDCl_3_) δ 165.40 – 163.76 (m), 163.56, 162.69 – 162.22 (m), 161.80, 161.24 (dd, *J* = 15.4, 9.3 Hz), 159.90, 159.57 (dd, *J* = 15.3, 9.1 Hz), 142.78, 130.31, 129.71, 128.21, 102.48, 101.57 – 99.79 (m), 48.54, 25.76, 24.56 ppm. MS(ESI) m/z for C_19_H_15_ClF_3_N_3_S (409.1) observed [M+H]+: 410.1.

**1-(6-Chloro-2-(thiophen-2-yl)-5-(2,4,6-trifluorophenyl)pyrimidin-4-yl)azepane (26, BL-0918)**. Synthesized according to General procedure E, using 4,6-dichloro-2-(thiophen-2-yl)-5-(2,4,6-trifluorophenyl) pyrimidine (0.030 g, 0.083 mmol, 1.00 eq) and azepane (0.018 g, 0.18 mmol, 2.20 eq). Purification via silica gel column chromatography furnished the desired product (0.019 g, 0.045 mmol, 54%). ^1^H NMR (600 MHz, CDCl_3_) δ 7.98 (d, *J* = 3.9 Hz, 1H), 7.45 (d, *J* = 5.3 Hz, 1H), 7.11 (t, *J* = 4.5 Hz, 1H), 6.76 (t, *J* = 7.8 Hz, 2H), 3.41 (t, *J* = 6.2 Hz, 4H), 1.67 (q, *J* = 5.5 Hz, 4H), 1.50 (dt, *J* = 6.4, 3.1 Hz, 4H) ppm. ^13^C NMR (150 MHz, CDCl_3_) δ 164.02 (d, *J* = 15.9 Hz), 162.36 (d, *J* = 14.8 Hz), 162.22, 161.82, 161.50 (dd, *J* = 15.5, 9.3 Hz), 159.85 (dd, *J* = 15.3, 9.2 Hz), 159.42, 143.00, 130.12, 129.45, 128.13, 111.08 (dd, *J* = 42.4, 5.2 Hz), 101.55 – 100.25 (m), 99.98, 50.57, 27.96, 26.82 ppm. MS(ESI) m/z for C_20_H_17_ClF_3_N_3_S (423.1) observed [M+H]+: 424.1.

**3-(6-Chloro-2-(thiophen-2-yl)-5-(2,4,6-trifluorophenyl)pyrimidin-4-yl)oxazolidine (27, BL-0909)**. Synthesized according to General procedure E, using 4,6-dichloro-2-(thiophen-2-yl)-5-(2,4,6-trifluorophenyl) pyrimidine (0.030 g, 0.083 mmol, 1.00 eq) and oxazolidine (0.012 g, 0.17 mmol, 2.00 eq). Purification via silica gel column chromatography furnished the desired product (0.019 g, 0.048 mmol, 57%). ^1^H NMR (600 MHz, CDCl_3_) δ 8.02 – 7.96 (m, 1H), 7.50 – 7.46 (m, 1H), 7.13 (dd, *J* = 4.9, 3.8 Hz, 1H), 6.79 (dd, *J* = 8.6, 6.6 Hz, 2H), 4.96 (s, 2H), 4.01 (t, *J* = 6.4 Hz, 2H), 3.23 (t, *J* = 6.4 Hz, 2H) ppm. ^13^C NMR (151 MHz, CDCl_3_) δ 164.52 (t, *J* = 15.0 Hz), 162.84 (t, *J* = 15.0 Hz), 162.18 (dd, *J* = 14.7, 8.8 Hz), 161.50, 160.52 (dd, *J* = 14.8, 8.9 Hz), 160.24, 158.62, 142.28, 130.66, 130.03, 128.28, 109.05 (d, *J* = 5.2 Hz), 100.16, 80.82, 67.65, 45.62 ppm. HRMS (ES^+^) calcd for [C_17_H_12_ClF_3_N_3_OS]^+^ 398.0336, found 398.0337.

**4-(6-Chloro-2-(thiophen-2-yl)-5-(2,4,6-trifluorophenyl)pyrimidin-4-yl)morpholine (28, BL-0908)**. Synthesized according to General procedure E, using 4,6-dichloro-2-(thiophen-2-yl)-5-(2,4,6-trifluorophenyl) pyrimidine (0.030 g, 0.083 mmol, 1.00 eq) and morpholine (0.014 g, 0.17 mmol, 2.00 eq). Purification via silica gel column chromatography furnished the desired product (0.026 g, 0.063 mmol, 76%). ^1^H NMR (600 MHz, CDCl_3_) δ 7.99 (d, *J* = 3.9 Hz, 1H), 7.48 (d, *J* = 5.0 Hz, 1H), 7.12 (t, *J* = 4.4 Hz, 1H), 6.81 (dd, *J* = 8.7, 6.9 Hz, 2H), 3.64 – 3.60 (m, 4H), 3.44 – 3.41 (m, 4H) ppm. ^13^C NMR (151 MHz, CDCl_3_) δ 164.25 (t, *J* = 15.1 Hz), 163.45, 162.58 (t, *J* = 15.1 Hz), 162.17, 161.16 (dd, *J* = 14.7, 9.3 Hz), 160.03, 159.50 (dd, *J* = 14.8, 9.3 Hz), 142.32, 130.68, 130.04, 128.31, 109.91 (td, *J* = 20.6, 4.6 Hz), 102.90, 101.06 (t, *J* = 27.7 Hz), 66.58, 47.55 ppm. HRMS (ES^+^) calcd for [C_18_H_13_ClF_3_N_3_OS]^+^ 412.0493, found 412.0493.

**4-(6-Chloro-2-(thiophen-2-yl)-5-(2,4,6-trifluorophenyl)pyrimidin-4-yl)-1,4-oxazepane (29, BL-0906)**. Synthesized according to General procedure E, using 4,6-dichloro-2-(thiophen-2-yl)-5-(2,4,6-trifluorophenyl) pyrimidine (0.030 g, 0.083 mmol, 1.00 eq), 1,4-oxazepane hydrochloride (0.017 g, 0.120 mmol, 1.50 eq), and Et_3_N (0.025 g, 0.250 mmol, 3.00 eq). Purification via silica gel column chromatography furnished the desired product (0.024 g, 0.056 mmol, 68%). ^1^H NMR (600 MHz, CDCl_3_) δ 7.72 (dd, *J* = 302.7, 4.5 Hz, 1H), 7.22 – 6.52 (m, 1H), 4.45 – 3.03 (m, 4H), 1.86 (p, *J* = 6.0 Hz, 1H). ^13^C NMR (150 MHz, CDCl_3_) δ 163.28 – 162.82 (m), 161.65, 161.50 – 161.20 (m), 161.04, 160.32 (dd, *J* = 14.6, 9.1 Hz), 158.66 (dd, *J* = 14.7, 9.1 Hz), 158.53, 141.52, 129.31, 128.64, 127.15, 109.46 (d, *J* = 4.9 Hz), 100.03 – 99.42 (m), 69.15, 68.84, 51.95, 47.90, 28.48 ppm. HRMS (ES^+^) calcd for [C_19_H_16_ClF_3_N_3_OS]^+^ 426.0649, found 426.0652.

**6-Chloro-*N*-neopentyl-2-(thiophen-2-yl)-5-(2,4,6-trifluorophenyl)pyrimidin-4-amine (30, BL-0919)**. Synthesized according to General procedure E, using 4,6-dichloro-2-(thiophen-2-yl)-5-(2,4,6-trifluorophenyl) pyrimidine (0.030 g, 0.083 mmol, 1.00 eq) and 2,2-dimethylpropan-1-amine (0.016 g, 0.18 mmol, 2.20 eq). Purification via silica gel column chromatography furnished the desired product (0.032 g, 0.078 mmol, 94%). ^1^H NMR (600 MHz, CDCl_3_) δ 8.00 (d, *J* = 3.7 Hz, 1H), 7.47 (d, *J* = 5.0 Hz, 1H), 7.14 – 7.11 (m, 1H), 6.86 (dd, *J* = 8.8, 6.8 Hz, 2H), 4.62 (t, *J* = 6.8 Hz, 1H), 3.41 (d, *J* = 6.6 Hz, 2H), 0.90 (s, 9H) ppm. ^13^C NMR (150 MHz, CDCl_3_) δ 165.19 – 164.28 (m), 163.00 (t, *J* = 14.9 Hz), 161.92 (dd, *J* = 15.0, 9.3 Hz), 161.76, 160.90, 160.25 (dd, *J* = 14.7, 9.0 Hz), 159.29, 142.90, 130.39, 129.76, 128.20, 106.58 – 105.83 (m), 102.75 – 101.10 (m), 100.77, 52.18, 32.83, 27.34 ppm. MS(ESI) m/z for C_19_H_17_ClF_3_N_3_S (411.1) observed [M+H]+: 412.1.

***N*-Benzyl-6-chloro-2-(thiophen-2-yl)-5-(2,4,6-trifluorophenyl)pyrimidin-4-amine (31, BL-0926)**. Synthesized according to General procedure E, using 4,6-dichloro-2-(thiophen-2-yl)-5-(2,4,6-trifluorophenyl) pyrimidine (0.100 g, 0.277 mmol, 1.00 eq) and benzylamine (0.020 g, 0.18 mmol, 2.20 eq). Purification via silica gel column chromatography furnished the desired product (0.032 g, 0.074 mmol, 89%). ^1^H NMR (600 MHz, CDCl_3_) δ 8.00 (d, *J* = 3.7 Hz, 1H), 7.47 (d, *J* = 4.8 Hz, 1H), 7.36 – 7.30 (m, 4H), 7.28 (dd, *J* = 6.5, 2.3 Hz, 1H), 7.12 (dd, *J* = 5.0, 3.7 Hz, 1H), 6.85 – 6.79 (m, 2H), 4.89 (t, *J* = 5.8 Hz, 1H), 4.77 (d, *J* = 5.7 Hz, 2H) ppm. ^13^C NMR (150 MHz, CDCl_3_) δ 164.68 (t, *J* = 14.8 Hz), 163.00 (t, *J* = 15.0 Hz), 161.90 (dd, *J* = 15.1, 9.2 Hz), 161.20, 161.06, 160.23 (dd, *J* = 15.2, 9.1 Hz), 159.53, 142.63, 138.38, 130.60, 130.03, 128.86, 128.23, 127.69, 106.67 – 105.67 (m), 103.53 – 99.95 (m), 45.48 ppm. MS(ESI) m/z for C_21_H_13_ClF_3_N_3_S (431.0) observed [M+H]+: 432.0.

**6-Chloro-*N*-(pyridin-3-ylmethyl)-2-(thiophen-2-yl)-5-(2,4,6-trifluorophenyl)pyrimidin-4-amine (32, BL-0927)**. Synthesized according to General procedure E, using 4,6-dichloro-2-(thiophen-2-yl)-5-(2,4,6-trifluorophenyl) pyrimidine (0.100 g, 0.277 mmol, 1.00 eq) and pyridine-3-ylmethanamine (0.020 g, 0.18 mmol, 2.20 eq). Purification via silica gel column chromatography furnished the desired product (0.033 g, 0.076 mmol, 92%). ^1^H NMR (600 MHz, DMSO) δ 8.60 (s, 1H), 8.48 – 8.39 (m, 1H), 7.96 (t, *J* = 6.0 Hz, 1H), 7.88 (d, *J* = 3.9 Hz, 1H), 7.79 (d, *J* = 5.0 Hz, 1H), 7.71 (dd, *J* = 7.9, 2.0 Hz, 1H), 7.48 – 7.41 (m, 2H), 7.34 (dd, *J* = 7.9, 4.8 Hz, 1H), 7.22 – 7.15 (m, 1H), 4.59 (d, *J* = 5.9 Hz, 2H) ppm. ^13^C NMR (150 MHz, DMSO) δ 161.44 (dd, *J* = 15.8, 10.0 Hz), 160.96, 159.97, 159.79 (dd, *J* = 15.9, 9.4 Hz), 158.15, 148.91, 148.12, 141.65, 134.97, 134.88, 131.69, 129.81, 128.55, 123.50, 107.24 – 105.24 (m), 101.64 (td, *J* = 26.6, 4.7 Hz), 100.88, 41.92 ppm. MS(ESI) m/z for C_20_H_12_ClF_3_N_4_S (432.0) observed [M+H]+: 433.0.

**6-Chloro-*N*-(2-(methylsulfonyl)ethyl)-2-(thiophen-2-yl)-5-(2,4,6-trifluorophenyl)pyrimidin-4-amine (33, BL-0791).** Synthesized according to General procedure E, using 4,6-dichloro-2-(thiophen-2-yl)-5-(2,4,6-trifluorophenyl)pyrimidine (0.100 g, 0.277 mmol, 1.00 eq), 2-(methylsulfonyl)ethan-1-aminium chloride (0.044 g, 0.277 mmol, 1.00 eq), and Et_3_N (0.056 g, 0.554 mmol, 2.00 eq). Purification via silica gel column chromatography furnished the desired product (0.105 g, 0.234 mmol, 84%). ^1^H NMR (600 MHz, CDCl3) δ 8.04 – 7.99 (m, 1H), 7.50 (d, J = 4.9 Hz, 1H), 7.17 – 7.12 (m, 1H), 6.84 (t, J = 8.2 Hz, 2H), 5.36 (t, J = 5.9 Hz, 1H), 4.06 (q, J = 6.0 Hz, 2H), 3.41 (t, J = 6.0 Hz, 2H), 2.94 (s, 3H) ppm. ^13^C NMR (150 MHz, CDCl3) δ 164.01 (dt, J = 253.2, 15.2 Hz), 160.96 (ddd, J = 251.7, 15.1, 9.1 Hz), 160.93 (d, J = 6.9 Hz), 159.94, 142.24, 130.82, 130.24, 128.47, 105.40 (d, J = 4.9 Hz), 102.00, 101.79 – 101.19 (m), 53.49, 42.01, 35.67 ppm. HRMS (ES^+^): calcd for C_17_H_14_ClF_3_N_3_O_2_S_2_ [M + H]^+^, 448.0163; found, 448.0158.

**2,2’-((6-Chloro-2-(thiophen-2-yl)-5-(2,4,6-trifluorophenyl)pyrimidin-4-yl)azanediyl)bis(ethan-1-ol) (34, BL-0795)**. Synthesized according to General procedure E, using 4,6-dichloro-2-(thiophen-2-yl)-5-(2,4,6-trifluorophenyl)pyrimidine (0.050 g, 0.138 mmol, 1.00 eq) and diethanolamine (0.029 g, 0.277 mmol, 2.00 eq). Purification via silica gel column chromatography furnished the desired product(0.054 g, 0.126 mmol, 90%). ^1^H NMR (600 MHz, CDCl_3_) δ 7.95 (d, *J* = 3.7 Hz, 1H), 7.47 (d, *J* = 5.0 Hz, 1H), 7.12 (t, *J* = 4.4 Hz, 1H), 6.79 (t, *J* = 8.1 Hz, 2H), 3.66 (t, *J* = 5.0 Hz, 4H), 3.55 (t, *J* = 4.9 Hz, 6H) ppm. ^13^C NMR (150 MHz, CDCl_3_) δ 163.52 (dt, *J* = 252.8, 15.2 Hz), 163.19, 162.46, 160.56 (ddd, *J* = 249.8, 14.8, 9.1 Hz), 159.55, 142.04, 130.75, 130.02, 128.48, 110.02 (td, *J* = 20.8, 4.9 Hz), 102.42, 101.31 – 100.73 (m), 60.99, 54.29 ppm. HRMS (ES^+^): calcd for C_18_H_16_ClF_3_N_3_O_2_S [M + H]^+^, 430.0598; found, 430.0592.

**4-(4-Chloro-6-(((3-methyloxetan-3-yl)methyl)amino)-2-(thiophen-2-yl)pyrimidin-5-yl)-3,5-difluorophenol (35, BL-1050).** Synthesized according to General procedure G, using 6-chloro-5-(2,6-difluoro-4-methoxyphenyl)-N-((3-methyloxetan-3-yl)methyl)-2-(thiophen-2-yl)pyrimidin-4-amine (0.030 g, 0.069 mmol, 1.00 eq) and BBr_3_ (0.086 g, 0.340 mmol, 5.00 eq). Purification via silica gel column chromatography furnished the desired product as a hydrobromide salt (0.012 g, 0.028 mmol, 41%) ^1^H NMR (600 MHz, DMSO) δ 10.71 (s, 1H), 7.92 (d, *J* = 3.7 Hz, 1H), 7.77 (d, *J* = 5.0 Hz, 1H), 7.20 (q, *J* = 4.2 Hz, 1H), 7.02 (t, *J* = 6.2 Hz, 1H), 6.64 (d, *J* = 10.3 Hz, 2H), 4.84 (s, 1H), 3.64 – 3.57 (m, 1H), 3.57 – 3.51 (m, 2H), 3.46 (d, *J* = 10.0 Hz, 1H), 3.39 (s, 2H), 3.37 – 3.24 (m, 1H), 0.87 (s, 3H) ppm. ^13^C NMR (150 MHz, DMSO) δ 162.03, 161.65, 160.71 (t, *J* = 14.8 Hz), 160.21 – 159.88 (m), 159.41, 158.36, 156.49, 142.06, 131.36, 129.52, 128.55, 102.04, 99.94 (d, *J* = 24.9 Hz), 99.20, 65.24, 45.16, 41.71, 41.32, 18.79 ppm. MS(ESI) m/z for C_19_H_16_ClF_2_N_3_O_2_S (423.1) observed [M+H]+: 424.1.

**6- Chloro-5-(2,6-difluoro-4-methoxyphenyl)-*N*-((3-methyloxetan-3-yl)methyl)-2-(thiophen-2-yl)pyrimidin-4-amine (36, BL-1049)**. Synthesized according to General procedure E, using 4,6-Dichloro-5-(2,6-difluoro-4-methoxyphenyl)-2-(thiophen-2-yl)pyrimidine (0.200 g, 0.536 mmol, 1.00 eq), and (3-methyloxetan-3-yl)methanamine (0.114 g, 1.13 mmol, 2.10 eq). Purification via silica gel column chromatography furnished the desired product (0.205 g, 0.468 mmol, 87%). ^1^H NMR (600 MHz, CDCl_3_) δ 7.99 (dd, *J* = 3.7, 1.2 Hz, 1H), 7.45 (dd, *J* = 5.0, 1.3 Hz, 1H), 7.11 (dd, *J* = 5.0, 3.7 Hz, 1H), 6.64 – 6.57 (m, 2H), 5.00 (t, *J* = 6.1 Hz, 1H), 4.51 (d, *J* = 6.1 Hz, 2H), 4.35 (d, *J* = 6.0 Hz, 2H), 3.84 (s, 3H), 3.77 (d, *J* = Hz, 2H), 1.29 (s, 3H) ppm. MS(ESI) m/z for C_20_H_18_ClF_2_N_3_O_2_S (437.1) observed [M+H]+: 438.1.

**6-Chloro-5-(2,6-difluoro-4-nitrophenyl)-*N*-((3-methyloxetan-3-yl)methyl)-2-(thiophen-2-yl)pyrimidin-4-amine (37, BL-1001)**. Synthesized according to General procedure E, using 4,6-Dichloro-5-(2,6-difluoro-4-nitrophenyl)-2-(thiophen-2-yl)pyrimidine (0.050 g, 0.129 mmol, 1.00 eq), and (3-methyloxetan-3-yl)methanamine (0.027 g, 0.271 mmol, 2.10 eq). Purification via silica gel column chromatography furnished the desired product (0.052 g, 0.115 mmol, 89%). ^1^H NMR (600 MHz, CDCl_3_) δ 8.00 (dd, *J* = 3.7, 1.3 Hz, 1H), 7.49 (dd, *J* = 5.1, 1.3 Hz, 1H), 7.40 – 7.33 (m, 2H), 7.13 (dd, *J* = 5.0, 3.7 Hz, 1H), 5.19 (t, *J* = 6.0 Hz, 1H), 4.50 (d, *J* = 6.1 Hz, 2H), 4.37 (d, *J* = 6.1 Hz, 2H), 3.78 (d, *J* = 6.0 Hz, 2H), 2.59 (s, 2H), 2.02 (d, *J* = 3.9 Hz, 1H), 1.30 (s, 3H) ppm. ^13^C NMR (150 MHz, CDCl_3_) δ 161.46, 161.38 (d, *J* = 6.4 Hz), 161.15, 159.68 (d, *J* = 6.8 Hz), 159.11, 149.56 (t, *J* = 10.8 Hz), 142.13, 131.14, 130.46, 128.42, 116.84 (t, *J* = 20.8 Hz), 108.78 – 108.34 (m), 100.02, 80.53, 47.92, 40.66, 21.86 ppm. HRMS (ES^+^): calcd for C_19_H_16_ClF_2_N_4_O_3_S [M + H]^+^, 453.0594; found, 453.0599.

**5-(4-Amino-2,6-difluorophenyl)-6-chloro-N-((3-methyloxetan-3-yl)methyl)-2-(thiophen-2-yl)pyrimidin-4-amine (38, BL-1005)**

Synthesized according to General procedure G, using 6-chloro-5-(2,6-difluoro-4-nitrophenyl)-N-((3-methyloxetan-3-yl)methyl)-2-(thiophen-2-yl)pyrimidin-4-amine (0.020 g, 0.044 mmol, 1.00 eq). Purification via silica gel column chromatography furnished the desired product(0.016 g, 0.038 mmol, 86%). ^1^H NMR (600 MHz, CDCl_3_) δ 7.99 (dt, *J* = 3.9, 1.4 Hz, 1H), 7.47 – 7.41 (m, 1H), 7.11 (ddd, *J* = 5.1, 3.7, 1.4 Hz, 1H), 6.34 – 6.28 (m, 2H), 5.04 (t, *J* = 6.1 Hz, 1H), 4.53 (dd, *J* = 6.1, 1.5 Hz, 2H), 4.36 (dd, *J* = 6.1, 1.5 Hz, 2H), 3.79 – 3.75 (m, 2H), 1.30 (s, 3H) ppm. ^13^C NMR (150 MHz, CDCl_3_) δ 162.52 – 162.08 (m), 160.64 (d, *J* = 9.7 Hz), 160.29, 159.63, 150.12 (t, *J* = 13.8 Hz), 142.82, 130.19, 129.60, 128.21, 102.80, 98.53 – 98.18 (m), 97.72 (t, *J* = 21.5 Hz), 80.62, 47.68, 40.70, 21.94 ppm. HRMS (ES^+^): calcd for C_19_H_20_ClF_2_N_4_S [M + H]^+^, 409.1060; found, 409.1063.

**4-(4-Chloro-6-(((3-methyloxetan-3-yl)methyl)amino)-2-(thiophen-2-yl)pyrimidin-5-yl)-3,5-difluorobenzonitrile (39, BL-1002)**

Synthesized according to General procedure E, using 4-(4,6-dichloro-2-(thiophen-2-yl)pyrimidin-5-yl)-3,5-difluorobenzonitrile (0.020 g, 0.054 mmol, 1.00 eq), and (3-methyloxetan-3-yl)methanamine (0.012 g, 0.110 mmol, 2.10 eq). Purification via silica gel column chromatography furnished the desired product (0.022 g, 0.051 mmol, 94%). ^1^H NMR (600 MHz, CDCl_3_) δ 8.05 – 8.01 (m, 1H), 7.94 (d, *J* = 6.5 Hz, 2H), 7.55 – 7.49 (m, 1H), 7.18 – 7.12 (m, 1H), 5.05 (t, *J* = 5.8 Hz, 1H), 4.49 (d, *J* = 6.1 Hz, 2H), 4.40 (d, *J* = 6.1 Hz, 2H), 3.80 (d, *J* = 5.8 Hz, 2H), 1.31 (s, 3H) ppm. ^13^C NMR (150 MHz, CDCl_3_) δ 161.62 (d, *J* = 6.8 Hz), 161.41, 161.22, 159.92 (d, *J* = 6.9 Hz), 159.15, 142.20, 131.08, 130.41, 128.41, 116.76 – 116.32 (m), 116.16, 115.79 (t, *J* = 20.5 Hz), 115.28 (t, *J* = 11.8 Hz), 100.07, 80.53, 47.86, 40.68, 21.89 ppm. HRMS (ES^+^): calcd for C_20_H_16_ClF_2_N_4_OS [M + H]^+^, 433.0696; found, 433.0700.

**6-Chloro-5-(2,6-difluoro-4-nitrophenyl)-N-neopentyl-2-(thiophen-2-yl)pyrimidin-4-amine (40, BL-0990)**

Synthesized according to General procedure E, using 4,6-Dichloro-5-(2,6-difluoro-4-nitrophenyl)-2-(thiophen-2-yl)pyrimidine (0.120 g, 0.309 mmol, 1.00 eq) and 2,2-dimethylpropan-1-amine (0.059 g, 0.680 mmol, 2.20 eq). Purification via silica gel column chromatography furnished the desired product(0.094 g, 0.214 mmol, 69%). ^1^H NMR (600 MHz, CDCl_3_) δ 8.01 (d, *J* = 3.7 Hz, 1H), 7.96 (d, *J* = 6.4 Hz, 2H), 7.49 (d, *J* = 5.0 Hz, 1H), 7.13 (t, *J* = 4.3 Hz, 1H), 4.60 (t, *J* = 6.5 Hz, 1H), 3.44 (d, *J* = 6.6 Hz, 2H), 0.91 (s, 9H) ppm. ^13^C NMR (150 MHz, CDCl_3_) δ 161.42, 161.37, 161.13, 159.70 (d, *J* = 6.7 Hz), 158.87, 149.52 (t, *J* = 10.7 Hz), 142.50, 130.86, 130.16, 128.30, 117.16 (t, *J* = 21.0 Hz), 109.33 – 107.70 (m), 99.59, 52.25, 32.88, 27.36 ppm. HRMS (ES^+^): calcd for C_18_H_14_ClF_5_N_3_OS [M + H]^+^, 439.0808; found, 439.0803.

**5-(4-Amino-2,6-difluorophenyl)-6-chloro-N-neopentyl-2-(thiophen-2-yl)pyrimidin-4-amine (41, BL-1004)**

Synthesized according to General procedure G, using 6-chloro-5-(2,6-difluoro-4-nitrophenyl)-N-neopentyl-2-(thiophen-2-yl)pyrimidin-4-amine (0.020 g, 0.046 mmol, 1.00 eq). Purification via silica gel column chromatography furnished the desired product (0.012 g, 0.029 mmol, 64%). ^1^H NMR (600 MHz, CDCl_3_) δ 7.98 (d, *J* = 3.6 Hz, 1H), 7.44 (d, *J* = 5.1 Hz, 1H), 7.13 – 7.09 (m, 1H), 6.32 (d, *J* = 9.2 Hz, 2H), 4.80 (t, *J* = 6.3 Hz, 1H), 3.63 (s, 2H), 3.39 (d, *J* = 6.4 Hz, 2H), 0.90 (s, 9H) ppm. ^13^C NMR (150 MHz, CDCl_3_) δ 162.33, 162.26, 160.66 (d, *J* = 9.1 Hz), 160.24, 159.28, 149.93, 143.16, 129.99, 129.36, 128.12, 102.43, 98.42 (d, *J* = 26.8 Hz), 98.05 (d, *J* = 22.5 Hz), 52.14, 32.83, 27.35 ppm. HRMS (ES^+^): calcd for C_19_H_18_ClF_2_N_4_OS [M + H]^+^, 423.0852; found, 423.0849.

**4-(4-Chloro-6-(neopentylamino)-2-(thiophen-2-yl)pyrimidin-5-yl)-3,5-difluorobenzonitrile (42, BL-0997)**

Synthesized according to General procedure E, using 4-(4,6-dichloro-2-(thiophen-2-yl)pyrimidin-5-yl)-3,5-difluorobenzonitrile (0.030 g, 0.082 mmol, 1.00 eq) and 2,2-dimethylpropan-1-amine (0.015 g, 0.171 mmol, 2.10 eq). Purification via silica gel column chromatography furnished the desired product (0.030 g, 0.082 mmol, 88%). ^1^H NMR (600 MHz, CDCl_3_) δ 8.03 (d, *J* = 3.7 Hz, 1H), 7.50 (d, *J* = 5.2 Hz, 1H), 7.41 (d, *J* = 5.7 Hz, 2H), 7.16 – 7.11 (m, 1H), 4.55 (s, 1H), 3.42 (d, *J* = 6.3 Hz, 2H), 2.00 (s, 1H), 0.92 – 0.89 (m, 9H) ppm. ^13^C NMR (150 MHz, CDCl_3_) δ 161.65 (d, *J* = 6.9 Hz), 161.25 (d, *J* = 15.2 Hz), 159.95 (d, *J* = 6.8 Hz), 158.93, 142.51, 130.88, 130.22, 128.33, 116.85 – 116.33 (m), 116.22, 116.04, 115.90, 115.34 (t, *J* = 11.5 Hz), 99.70, 52.28, 32.87, 27.38 ppm. HRMS (ES^+^): calcd for C_20_H_18_ClF_2_N_4_S [M + H]^+^, 419.0903; found, 419.0901.

**6-Chloro-N-(cyclobutylmethyl)-5-(2,6-difluoro-4-nitrophenyl)-2-(thiophen-2-yl)pyrimidin-4-amine (43, BL-0991)**.

Synthesized according to General procedure E, using 4,6-Dichloro-5-(2,6-difluoro-4-nitrophenyl)-2-(thiophen-2-yl)pyrimidine (0.062 g, 0.160 mmol, 1.00 eq), cyclobutylmethanamine hydrochloride (0.023 g, 0.192 mmol, 1.20 eq) and Et_3_N (0.032 g, 0.319 mmol, 2.00 eq). Purification via silica gel column chromatography furnished the desired product (0.094 g, 0.214 mmol, 69%). ^1^H NMR (600 MHz, CDCl_3_) δ 8.01 (dd, *J* = 3.6, 1.4 Hz, 1H), 7.91 (d, *J* = 6.4 Hz, 2H), 7.50 (dd, *J* = 5.0, 1.5 Hz, 1H), 7.14 (dd, *J* = 5.2, 3.6 Hz, 1H), 4.62 (t, *J* = 5.8 Hz, 1H), 3.60 (dd, *J* = 7.5, 5.5 Hz, 2H), 2.65 – 2.54 (m, 1H), 2.08 – 2.00 (m, 2H), 1.94 – 1.85 (m, 2H), 1.78 – 1.68 (m, 2H) ppm. ^13^C NMR (150 MHz, CDCl_3_) δ 161.44, 161.37, 160.85, 159.70 (d, *J* = 6.8 Hz), 158.76, 149.44 (t, *J* = 10.8 Hz), 142.44, 130.91, 130.23, 128.32, 117.17 (t, *J* = 21.0 Hz), 109.38 – 108.11 (m), 99.78, 46.85, 35.11, 25.87, 18.48 ppm. HRMS (ES^+^): calcd for C_19_H_16_ClF_2_N_4_O_2_S [M + H]^+^, 437.0645; found, 437.0641.

**5-(4-Amino-2,6-difluorophenyl)-6-chloro-N-(cyclobutylmethyl)-2-(thiophen-2-yl)pyrimidin-4-amine (44, BL-1003)**

Synthesized according to General procedure F, using 6-chloro-N-(cyclobutylmethyl)-5-(2,6-difluoro-4-nitrophenyl)-2-(thiophen-2-yl)pyrimidin-4-amine (0.020 g, 0.048 mmol, 1.00 eq). Purification via silica gel column chromatography furnished the desired product (0.011 g, 0.028 mmol, 61%). ^1^H NMR (600 MHz, CDCl_3_) δ 7.99 (d, *J* = 3.7 Hz, 1H), 7.45 (dd, *J* = 5.1, 1.3 Hz, 1H), 7.11 (dd, *J* = 5.0, 3.6 Hz, 1H), 6.37 – 6.27 (m, 2H), 4.71 (t, *J* = 5.7 Hz, 1H), 4.09 (s, 2H), 3.59 – 3.52 (m, 2H), 2.56 (hept, *J* = 7.6 Hz, 1H), 2.08 – 1.98 (m, 2H), 1.92 – 1.83 (m, 2H), 1.75 – 1.66 (m, 2H) ppm.^13^C NMR (150 MHz, CDCl_3_) δ 162.31 (d, *J* = 9.7 Hz), 162.00, 160.66 (d, *J* = 9.8 Hz), 160.35, 159.25, 149.88 (t, *J* = 13.8 Hz), 143.13, 129.98, 129.41, 128.11, 102.59, 98.51 – 98.23 (m), 98.17, 46.57, 35.25, 25.78, 18.50 ppm. HRMS (ES^+^): calcd for C_19_H_18_ClF_2_N_4_S [M + H]^+^, 407.0903; found, 407.0902.

**4-(4-Chloro-6-((cyclobutylmethyl)amino)-2-(thiophen-2-yl)pyrimidin-5-yl)-3,5-difluorobenzonitrile (45, BL-0996)**

Synthesized according to General procedure E, using 4-(4,6-dichloro-2-(thiophen-2-yl)pyrimidin-5-yl)-3,5-difluorobenzonitrile (0.030 g, 0.082 mmol, 1.00 eq), cyclobutylmethanamine hydrochloride (0.012 g, 0.098 mmol, 1.20 eq), and Et_3_N (0.016 g, 0.163 mmol, 2.00 eq). Purification via silica gel column chromatography furnished the desired product (0.028 g, 0.067 mmol, 82%). ^1^H NMR (600 MHz, CDCl_3_) δ 8.05 – 8.01 (m, 1H), 7.50 (d, *J* = 5.0 Hz, 1H), 7.39 (d, *J* = 5.8 Hz, 2H), 7.14 (dd, *J* = 5.0, 3.7 Hz, 1H), 4.42 (s, 1H), 3.58 (dd, *J* = 7.3, 5.7 Hz, 2H), 2.62 – 2.51 (m, 1H), 2.08 – 2.03 (m, 2H), 1.96 – 1.83 (m, 2H), 1.75 – 1.66 (m, 2H) ppm. ^13^C NMR (150 MHz, CDCl_3_) δ 161.66, 161.46, 160.95, 159.92, 158.89, 142.55, 130.87, 130.24, 128.32, 116.57, 116.53, 116.42, 116.38, 116.27, 116.06, 115.26, 99.87, 46.81, 35.17, 25.84, 18.51 ppm. HRMS (ES^+^): calcd for C_20_H_16_ClF_2_N_4_S [M + H]^+^, 417.0747; found, 417.0746.

**6-Chloro-5-(4-ethynyl-2,6-difluorophenyl)-N-((3-methyl-3H-diazirin-3-yl)methyl)-2-(thiophen-2-yl)pyrimidin-4-amine (46, BL-1044)**

To a solution of **78** (0. 120, 0.232 mmol, 1.00 eq), CuI (0.007 mg, 0.0348 mmol, 0.15 eq), and Pd(Ph_3_P)_4_ (0.0272 g, 0 0.023 mmol, 0.10 eq) in degassed DMF (1.00 mL 0.2 M), Et_3_N was added (0.070 mg, 0.696 mmol, 3.00 eq) followed by trimethylsilylacetylene (0.030 mg, 0.302 mmol, 1.30 eq). The resulting mixture was diluted with water and extracted with EtOAc (3x) and washed with water (5x). The combined organic extracts were collected, dried over Na_2_SO_4_, filtered, and evaporated under reduced pressure, and was used without further purification. ^1^H NMR (600 MHz, CDCl_3_) δ 8.08 (dd, *J* = 3.7, 1.2 Hz, 1H), 7.51 (dd, *J* = 5.0, 1.2 Hz, 1H), 7.15 (dd, *J* = 8.0, 3.9 Hz, 3H), 4.49 (s, 1H), 3.47 (d, *J* = 6.2 Hz, 2H), 1.11 (s, 3H), 0.32 – 0.22 (m, 12H). The resulting crude (0.066 g, 0.14 mmol, 1.00 eq) was immediately used in the subsequent reaction by reacting with K_2_CO_3_ (0.056 g, 0.41 mmol, 3.00 eq) in MeOH (1.0 mL, 0.14 M). The resulting mixture was stirred at rt for 1 h. The reaction was then filtered and purified by reverse-phase HPLC (10 to 90 % ACN in H_2_O, 0.1 % formic acid) obtaining the desired compound **46** (0.035 g, 0.084 mmol, 62%). ^1^H NMR (600 MHz, CDCl_3_) δ 7.99 – 7.95 (m, 1H), 7.43 – 7.39 (m, 1H), 7.05 (dq, *J* = 9.0, 4.0 Hz, 3H), 4.65 (t, *J* = 6.2 Hz, 1H), 3.40 (d, *J* = 6.3 Hz, 2H), 3.17 (s, 1H), 1.03 (s, 3H) ppm. ^13^C NMR (150 MHz, CDCl_3_) δ 161.00 (d, J = 7.5 Hz), 160.84 (d, J = 2.7 Hz), 159.32 (d, J = 7.4 Hz), 159.23 , 142.30 , 130.76 , 130.22 , 128.29 , 125.97 (t, J = 11.9 Hz), 117.46 – 114.91 (m), 110.43 (t, J = 20.8 Hz), 101.65 , 80.99 , 80.81 (t, J = 3.7 Hz), 44.96 , 25.73 , 18.08 ppm. MS(ESI) m/z for C_19_H_12_ClF_2_N_5_S (415) observed [M+H]+: 416. Malonates **47-52** were synthesized as previously published.

**Diethyl 2-(2,6-difluoro-4-methoxyphenyl)malonate (53)**

Synthesized according to General procedure A, using 2-bromo-1,3-difluoro-5-methoxybenzene (5.00 g, 22.4 mmol, 1.00 eq), diethyl malonate (10.8 g, 67.3 mmol, 3.00 eq), copper(I) bromide (3.86 g, 26.9 mmol, 1.20 eq), and sodium hydride (2.69 g, 67.3 mmol, 3.00 eq). Purification via silica gel column chromatography furnished the title compound (4.90 g, 16.2 mmol, 72%). ^1^H NMR (600 MHz, CDCl_3_) δ 6.48 (d, *J* = 9.6 Hz, 2H), 4.86 (s, 1H), 4.26 – 4.21 (m, 4H), 3.78 (s, 3H), 1.30 – 1.26 (m, 6H) ppm.

**Diethyl 2-(2,6-difluoro-4-nitrophenyl)malonate (54)**

Synthesized according to General procedure B, using 1,2,3-trifluoro-5-nitrobenzene (5.00 g, 28.2 mmol, 1.00 eq), diethyl malonate (1.50 g, 36.7 mmol, 1.30 eq) and sodium hydride (4.97 g, 31.1 mmol, 1.10 eq). Purification via silica gel column chromatography furnished the title compound (5.50 g, 17.3 mmol, 61%). ^1^H NMR (600 MHz, CDCl_3_) δ 7.84 (d, *J* = 7.0 Hz, 2H), 5.01 (s, 1H), 4.27 (q, *J* = 7.1 Hz, 4H), 1.29 (t, *J* = 7.1 Hz, 6H) ppm.

**Diethyl 2-(4-cyano-2,6-difluorophenyl)malonate (55**)

Synthesized according to General procedure B, using 3,4,5-trifluorobenzonitrile (3.00 g, 19.1 mmol, 1.00 eq), diethyl malonate (6.12 g, 38.2 mmol, 2.00 eq) and sodium hydride (1.60 g, 40.1 mmol, 2.10 eq). Purification via silica gel column chromatography furnished the title compound (3.01 g, 10.1 mmol, 53%). ^1^H NMR (600 MHz, CDCl_3_): δ

**7.30 (d, J = 7.0 Hz, 2H), 5.01 (s, 1H), 4.31–4.24 (m, 4H), 1.29 (t, J = 7.4 Hz, 6H) ppm.**

**2-(Thiophen-2-yl)-5-(2,4,6-trifluorophenyl)pyrimidine-4,6-diol (56)**

Synthesized according to General procedure C, using diethyl 2-(2,4,6-trifluorophenyl)malonate (2.00 g, 6.89 mmol, 1.00 eq), thiophene-2-carboximidamide hydrochloride (1.34 g, 8.27 mmol, 1.20 eq), and DBU (2.10 g, 13.8 mmol, 2.00 eq). The resulting precipitate was isolated by vacuum filtration and used in the subsequent step without further purification (1.93 g, 5.94 mmol, 86%). ^1^H NMR (600 MHz, DMSO) δ 8.26 (s, 1H), 7.94 (d, *J* = 5.3 Hz, 1H), 7.28 – 7.26 (m, 1H), 7.24 – 7.17 (m, 2H) ppm.

**4,6-Dichloro-5-(3,5-difluorophenyl)-2-(thiophen-2-yl)pyrimidine (57)**

Synthesized according to General procedure C, using diethyl 2-(3,5-difluorophenyl)malonate (0.200 g, 0.735 mmol, 1.00 eq), thiophene-2-carboximidamide hydrochloride (0.143 g, 0.882 mmol, 1.20 eq), and DBU (0.224 g, 1.47 mmol, 2.00 eq). The resulting precipitate was isolated by vacuum filtration and used in the subsequent step without further purification (0.115 g, 0.375 mmol, 51%). ^1^H NMR (600 MHz, DMSO) δ 7.46 – 7.39 (m, 1H), 7.10 (d, *J* = 5.0 Hz, 1H), 6.50 (d, *J* = 7.0 Hz, 2H), 6.44 (t, *J* = 4.5 Hz, 1H), 6.24 (tt, *J* = 9.4, 2.6 Hz, 1H) ppm.

**5-(2-Fluorophenyl)-2-(thiophen-2-yl)pyrimidine-4,6-diol (58)**

Synthesized according to General procedure C, using diethyl 2-(2-fluorophenyl)malonate (0.200 g, 0.787 mmol, 1.00 eq), thiophene-2-carboximidamide hydrochloride (0.154 g, 0.944 mmol, 1.20 eq), and DBU (0.240 g, 1.57 mmol, 2.00 eq). The resulting precipitate was isolated by vacuum filtration and used in the subsequent step without further purification (0.182 g, 0.631 mmol, 80%). ^1^H NMR (600 MHz, DMSO) δ 8.24 (s, 1H), 7.91 (d, *J* = 5.3 Hz, 1H), 7.35 (td, *J* = 7.7, 5.9 Hz, 2H), 7.26 (dd, *J* = 5.0, 3.7 Hz, 1H), 7.21 – 7.16 (m, 2H) ppm.

**5-(4-Fluorophenyl)-2-(thiophen-2-yl)pyrimidine-4,6-diol (59)**

Synthesized according to General procedure C, using diethyl 2-(4-fluorophenyl)malonate (0.090 g, 0.35 mmol, 1.00 eq), thiophene-2-carboximidamide hydrochloride (0.069 g, 0.42 mmol, 1.20 eq), and DBU (0.110 g, 0.71 mmol, 2.00 eq). The resulting precipitate was isolated by vacuum filtration and used in the subsequent step without further purification (0.040 g, 0.14 mmol, 39%). ^1^H NMR (600 MHz, DMSO) δ 8.19 (s, 1H), 7.86 (d, *J* = 5.0 Hz, 1H), 7.67 – 7.47 (m, 2H), 7.22 (dd, *J* = 5.0, 3.7 Hz, 1H), 7.13 (t, *J* = 9.0 Hz, 2H) ppm.

**5-(2,5-Difluorophenyl)-2-(thiophen-2-yl)pyrimidine-4,6-diol (60)**

Synthesized according to General procedure C, using diethyl 2-(2,5-difluorophenyl)malonate (0.100 g, 0.367 mmol, 1.00 eq), thiophene-2-carboximidamide hydrochloride (0.071 g, 0.441 mmol, 1.20 eq), and DBU (0.112 g, 0.735 mmol, 2.00 eq). The resulting precipitate was isolated by vacuum filtration and used in the subsequent step without further purification (0.072 g, 0.24 mmol, 64%). ^1^H NMR (600 MHz, DMSO) δ 8.23 (s, 1H), 7.91 (q, *J* = 7.1 Hz, 1H), 7.48 – 7.01 (m, 4H) ppm.

**5-(2,4-Difluorophenyl)-2-(thiophen-2-yl)pyrimidine-4,6-diol (61)**

Synthesized according to General procedure C, using diethyl 2-(2,5-difluorophenyl)malonate (0.100 g, 0.367 mmol, 1.00 eq), thiophene-2-carboximidamide hydrochloride (0.071 g, 0.441 mmol, 1.20 eq), and DBU (0.112 g, 0.735 mmol, 2.00 eq). The resulting precipitate was isolated by vacuum filtration and used in the subsequent step without further purification (0.037 g, 0.12 mmol, 33%). ^1^H NMR (600 MHz, DMSO) δ 8.26 – 8.19 (m, 1H), 7.90 (d, *J* = 5.0 Hz, 1H), 7.39 (td, *J* = 8.6, 6.7 Hz, 1H), 7.25 (dd, *J* = 5.2, 3.8 Hz, 1H), 7.21 (td, *J* = 9.8, Hz, 1H), 7.09 – 7.05 (m, 1H) ppm.

**5-(2,6-Difluoro-4-methoxyphenyl)-2-(thiophen-2-yl)pyrimidine-4,6-diol (62)**

Synthesized according to General procedure C, using diethyl 2-(2,6-difluoro-4-methoxyphenyl)malonate (0.500 g, 1.65 mmol, 1.00 eq), thiophene-2-carboximidamide hydrochloride (0.323 g, 1.98 mmol, 1.20 eq), and DBU (0.504 g, 3.31 mmol, 2.00 eq). The resulting precipitate was isolated by vacuum filtration and used in the subsequent step without further purification (0.520 g, 1.55 mmol, 94%). ^1^H NMR (600 MHz, DMSO) δ 8.24 (s, 1H), 7.92 (t, *J* = 7.0 Hz, 1H), 7.30 – 7.25 (m, 1H), 6.75 (dt, *J* = 9.4, 4.8 Hz, 2H), 3.81 (s, 3H) ppm.

**5-(2,6-Difluoro-4-nitrophenyl)-2-(thiophen-2-yl)pyrimidine-4,6-diol (63)**

Synthesized according to General procedure C, using diethyl 2-(2,6-difluoro-4-nitrophenyl)malonate (1.00 g, 3.65 mmol, 1.00 eq), thiophene-2-carboximidamide hydrochloride (0.615 g, 3.78 mmol, 1.20 eq), and DBU (0.960 g, 6.30 mmol, 2.00 eq). The resulting precipitate was isolated by vacuum filtration and used in the subsequent step without further purification (0.96 g, 2.70 mmol, 87%). ^1^H NMR (600 MHz, DMSO) δ 8.11 – 8.06 (m, 2H), 7.97 (d, *J* = 5.1 Hz, 2H), 7.33 (t, *J* = 4.5 Hz, 1H) ppm.

**4-(4,6-Dihydroxy-2-(thiophen-2-yl)pyrimidin-5-yl)-3,5-difluorobenzonitrile (64)**

Synthesized according to General procedure C, using diethyl 2-(4-cyano-2,6-difluorophenyl)malonate (2.00 g, 6.73 mmol, 1.00 eq), thiophene-2-carboximidamide hydrochloride (1.31 g, 8.07 mmol, 1.20 eq), and DBU (2.05 g, 13.5 mmol, 2.00 eq). The resulting precipitate was isolated by vacuum filtration and used in the subsequent step without further purification (1.85 g, 5.28 mmol, 79%). ^1^H NMR (600 MHz, DMSO) δ 7.91 (d, *J* = 7.6 Hz, 2H), 7.81 (d, *J* = 6.4 Hz, 2H), 7.31 – 7.25 (m, 1H) ppm.

**4,6-Dichloro-2-(thiophen-2-yl)-5-(2,4,6-trifluorophenyl)pyrimidine (65)**

Synthesized according to General procedure D, using 2-(thiophen-2-yl)-5-(2,4,6-trifluorophenyl)pyrimidine-4,6-diol (1.900 g, 5.86 mmol, 1.00 eq) and POCl_3_ (13.5 g, 87.9 mmol, 15.0 eq). Purification via silica gel column chromatography furnished the desired product (1.28 g, 3.56 mmol, 61%). ^1^H NMR (600 MHz, CDCl_3_) δ 8.13 (dd, *J* = 3.7, 1.3 Hz, 1H), 7.61 (dd, *J* = 5.0, 1.3 Hz, 1H), 7.18 (dd, *J* = 5.0, 3.8 Hz, 1H), 6.84 (dd, *J* = 8.7, 7.0 Hz, 2H) ppm.

**4,6-Dichloro-5-(3,5-difluorophenyl)-2-(thiophen-2-yl)pyrimidine (66)**

Synthesized according to General procedure D, using 5-(3,5-difluorophenyl)-2-(thiophen-2-yl)pyrimidine-4,6-diol (0.100 g, 0.326 mmol, 1.00 eq) and POCl_3_ (0.751 g, 4.90 mmol, 15.0 eq). Purification via silica gel column chromatography furnished the desired product (0.057 g, 0.17 mmol, 51%). ^1^H NMR (600 MHz, CDCl_3_) δ 8.11 (dd, *J* = 3.7, 1.3 Hz, 1H), 7.60 (dd, *J* = 5.0, 1.3 Hz, 1H), 7.18 (dd, *J* = 5.0, 3.7 Hz, 1H), 6.94 (tt, *J* = 9.0, 2.4 Hz, 1H), 6.91 – 6.85 (m, 2H) ppm.

**4,6-Dichloro-5-(2-fluorophenyl)-2-(thiophen-2-yl)pyrimidine (67)**

Synthesized according to General procedure D, using 5-(2-fluorophenyl)-2-(thiophen-2-yl)pyrimidine-4,6-diol (0.150 g, 0.520 mmol, 1.00 eq) and POCl_3_ (1.20 g, 7.80 mmol, 15.0 eq). Purification via silica gel column chromatography furnished the desired product (0.122 g, 0.375 mmol, 72%). ^1^H NMR (600 MHz, CDCl_3_) δ 8.11 (dd, *J* = 3.9, 1.1 Hz, 1H), 7.61 – 7.57 (m, 1H), 7.50 (tdd, *J* = 7.0, 5.2, 2.4 Hz, 1H), 7.33 – 7.27 (m, 2H), 7.22 (t, *J* = 8.9 Hz, 1H), 7.18 (dd, *J* = 4.9, 3.8 Hz, 1H) ppm.

**4,6-Dichloro-5-(4-fluorophenyl)-2-(thiophen-2-yl)pyrimidine (68)**

Synthesized according to General procedure D, using 5-(4-fluorophenyl)-2-(thiophen-2-yl)pyrimidine-4,6-diol (0.035 g, 0.120 mmol, 1.00 eq) and POCl_3_ (0.28 g, 1.80 mmol, 15.0 eq). The resulting crude residue was purified via column chromatography to furnish the desired product (0.029 g, 0.089 mmol, 73%). ^1^H NMR (600 MHz, CDCl_3_) δ 8.10 (d, *J* = 3.7 Hz, 1H), 7.58 (d, *J* = 4.8 Hz, 1H), 7.32 (dd, *J* = 8.2, 5.8 Hz, 2H), 7.24 – 7.16 (m, 3H) ppm.

**4,6-Dichloro-5-(2,5-difluorophenyl)-2-(thiophen-2-yl)pyrimidine (69)**

Synthesized according to General procedure D, using 5-(2,5-difluorophenyl)-2-(thiophen-2-yl)pyrimidine-4,6-diol (0.060 g, 0.200 mmol, 1.00 eq) and POCl_3_ (0.45 g, 2.90 mmol, 15.0 eq). Purification via silica gel column chromatography furnished the desired product (0.046 g, 0.13 mmol, 68%). ^1^H NMR (600 MHz, CDCl_3_) δ 8.12 (d, *J* = 3.7 Hz, 1H), 7.60 (d, *J* = 5.0 Hz, 1H), 7.19 (q, *J* = 5.2 Hz, 3H), 7.06 – 7.01 (m, 1H) ppm.

**4,6-Dichloro-5-(2,4-difluorophenyl)-2-(thiophen-2-yl)pyrimidine (70)**

Synthesized according to General procedure D, using 5-(2,4-difluorophenyl)-2-(thiophen-2-yl)pyrimidine-4,6-diol (0.030 g, 0.098 mmol, 1.00 eq) and POCl_3_ (0.23 g, 1.50 mmol, 15.0 eq). Purification via silica gel column chromatography furnished the desired product (0.017 g, 0.050 mmol, 51%). ^1^H NMR (600 MHz, CDCl_3_) δ 8.11 (dd, *J* = 3.9, 1.1 Hz, 1H), 7.60 (dd, *J* = 5.0, 1.1 Hz, 1H), 7.29 (td, *J* = 8.3, 6.0 Hz, 1H), 7.18 (dd, *J* = 4.9, 3.8 Hz, 1H), 7.04 (td, *J* = 8.2, 3.0 Hz, 1H), 6.98 (td, *J* = 9.0, 2.5 Hz, 1H) ppm.

**4,6-Dichloro-5-(2,6-difluoro-4-methoxyphenyl)-2-(thiophen-2-yl)pyrimidine (71)**

Synthesized according to General procedure D, using 5-(2,6-difluoro-4-methoxyphenyl)-2-(thiophen-2-yl)pyrimidine-4,6-diol (0.500 g, 1.49 mmol, 1.00 eq) and POCl_3_ (3.42 g, 22.3 mmol, 15.0 eq). Purification via silica gel column chromatography furnished the desired product (0.358 g, 0.959 mmol, 65%). ^1^H NMR (600 MHz, CDCl_3_) δ 8.11 (d, *J* = 3.1 Hz, 1H), 7.59 (d, *J* = 4.4 Hz, 1H), 7.20 – 7.16 (m, 1H), 6.63 – 6.57 (m, 2H), 3.87 (s, 3H) ppm.

**4,6-Dichloro-5-(2,6-difluoro-4-nitrophenyl)-2-(thiophen-2-yl)pyrimidine (72)**

Synthesized according to General procedure D, using 5-(2,6-difluoro-4-nitrophenyl)-2-(thiophen-2-yl)pyrimidine-4,6-diol (0.500 g, 1.42 mmol, 1.00 eq) and POCl_3_ (3.27 g, 21.4 mmol, 15.0 eq). Purification via silica gel column chromatography furnished the desired product (0.220 g, 0.567, 40%). ^1^H NMR (600 MHz, CDCl_3_) δ 8.16 (dd, *J* = 3.7, 1.3 Hz, 1H), 7.97 (d, *J* = 6.4 Hz, 2H), 7.65 (dd, *J* = 5.0, 1.3 Hz, 1H), 7.20 (dd, *J* = 5.0, 3.7 Hz, 1H) ppm.

**4-(4,6-Dichloro-2-(thiophen-2-yl)pyrimidin-5-yl)-3,5-difluorobenzonitrile (73)**. Synthesized according to General procedure D, using 4-(4,6-dihydroxy-2-(thiophen-2-yl)pyrimidin-5-yl)-3,5-difluorobenzonitrile (1.00 g, 3.02 mmol, 1.00 eq) and POCl_3_ (4.63 g, 30.2 mmol, 15.0 eq). Purification via silica gel column chromatography furnished the desired product (0.79 g, 2.15 mmol, 71%). ^1^H NMR (600 MHz, CDCl_3_) δ 8.15 – 8.13 (m, 1H), 7.64 – 7.62 (m, 1H), 7.39 (d, *J* = 6.1 Hz, 2H), 7.20 – 7.18 (m, 1H) ppm.

***tert*-Butyl (*R*)-(3-(4-(4-chloro-6-((3-methylbutan-2-yl)amino)-2-(thiophen-2-yl)pyrimidin-5-yl)-3,5-difluorophenoxy)propyl)(methyl)carbamate (74)**

To a solution of (R)-4-(4-chloro-6-((3-methylbutan-2-yl)amino)-2-(thiophen-2-yl)pyrimidin-5-yl)-3,5-difluorophenol (0.028 g, 0.068 mmol, 1.00) in anh. DMF (0.70 mL), K_2_CO_3_ (0.047 mmol, 0.34 mmol, 5.00 eq) was added, followed by *tert*-butyl (3-chloropropyl)(methyl)carbamate (0.021 g, 0.10 mmol, 1.50 eq). The reaction mixture was stirred 70 °C for 72 h. The resulting mixture was diluted with water and extracted with EtOAc (3x) and washed with water (5x). The combined organic extracts were collected, dried over Na_2_SO_4_, filtered, and evaporated under reduced pressure. Purification via silica gel column chromatography furnished the title compound (0.036 g, 0.062 mmol, 91%). ^1^H NMR (600 MHz, CDCl_3_) δ 8.03 – 7.96 (m, 1H), 7.45 (d, *J* = 5.0 Hz, 1H), 7.12 (dd, *J* = 5.0, 3.7 Hz, 1H), 6.62 – 6.58 (m, 2H), 4.39 (s, 1H), 4.26 (q, *J* = 6.8 Hz, 1H), 4.01 (t, *J* = 6.0 Hz, 2H), 3.43 (t, *J* = 6.8 Hz, 2H), 2.90 (s, 3H), 2.05 (d, *J* = 5.3 Hz, 2H), 1.82 (dt, *J* = 13.8, 6.9 Hz, 1H), 1.44 (s, 9H), 1.11 (d, *J* = 6.8 Hz, 3H), 0.88 (dd, *J* = 6.8, 3.9 Hz, 6H) ppm. Malonate **75** and diazirine **77** were synthesized as previously published.

**4,6-Dichloro-5-(2,6-difluoro-4-iodophenyl)-2-(thiophen-2-yl)pyrimidine (76)**

Synthesized according to General procedures C using diethyl 2-(2,6-difluoro-4-iodophenyl)malonate (1.70 g, 4.27 mmmol, 1.00 eq), thiophene-2-carboximidamide hydrochloride (0.833 g, 5.12 mmol, 1.20 eq), and DBU (1.30 g, 8.54 mmol, 2.00 eq) to obtain the intermediate (1.85 g, 4.27 mmol, 99%), which was immediately followed by General procedure D by reacting with POCl_3_ (3.99 mL, 42.8 mmol, 10.0 eq) to provide the dichloride **76** (1.49 g, 3.17 mmol, 74%). ^1^H NMR (400 MHz, CDCl_3_*d*) δ 8.13 (dd, *J* = 3.8, 1.3 Hz, 1H), 7.61 (dd, *J* = 5.0, 1.3 Hz, 1H), 7.48 – 7.40 (m, 2H), 7.18 (dd, *J* = 5.0, 3.8 Hz, 1H) ppm.

**6-Chloro-5-(2,6-difluoro-4-iodophenyl)-N-((3-methyl-3H-diazirin-3-yl)methyl)-2-(thiophen-2-yl)pyrimidin-4-amine (78)**

Synthesized according to General procedure E using 4,6-Dichloro-5-(2,6-difluoro-4-iodophenyl)-2-(thiophen-2-yl)pyrimidine (0.150 g, 0.320 mmol, 1.00 eq), 3-((chloro-λ^5^-azaneyl)methyl)-3-methyl-3H-diazirine (0.043 g, 0.352 mmol, 1.10 eq), and Et_3_N (0.049 g, 0.352 mmol 1.50 eq) to furnish the desired intermediate (0.120 g, 0.240 mmol, 74%). ^1^H NMR (600 MHz, CDCl_3_) δ 8.08 (dd, *J* = 3.8, 1.2 Hz, 1H), 7.51 (dd, *J* = 5.0, 1.2 Hz, 1H), 7.47 (d, *J* = 6.2 Hz, 2H), 7.15 (dd, *J* = 5.0, 3.7 Hz, 1H), 4.49 (s, 1H), 3.49 (d, *J* = 6.1 Hz, 2H), 1.12 (s, 3H) ppm.

## ACKNOWLEDGEMENTS

The research reported was supported by the NIH grant R21AI156554 to CRC and CB. KRF acknowledges support of the CARING T32 Training Grant (T32AI007036).

